# Atropselective Oxidation of 2,2’,3,3’,4,6’-Hexachlorobiphenyl (PCB 132) to Hydroxylated Metabolites by Human Liver Microsomes and Its Implications for PCB 132 Neurotoxicity

**DOI:** 10.1101/373837

**Authors:** Eric Uwimana, Brianna Cagle, Coby Yeung, Xueshu Li, Eric V. Patterson, Jonathan A. Doorn, Hans-Joachim Lehmler

## Abstract

Polychlorinated biphenyls (PCBs) have been associated with neurodevelopmental disorders. Several neurotoxic congeners display axial chirality and atropselectively affect cellular targets implicated in PCB neurotoxicity. Only limited information is available regarding the atropselective metabolism of these congeners in humans and their atropselective effects on neurotoxic outcomes. Here we investigate the hypothesis that the oxidation of 2,2’,3,3’,4,6’-hexachlorobiphenyl (PCB 132) by human liver microsomes (HLMs) and their effects on dopaminergic cells in culture are atropselective. Racemic PCB 132 was incubated with pooled or single donor HLMs, and levels and enantiomeric fractions of PCB 132 and its metabolites were determined gas chromatographically. The major metabolite was either 2,2’,3,4,4’,6’-hexachlorobiphenyl-3’-ol (3’-140), a 1,2-shift product, or 2,2’,3,3’,4,6’-hexachlorobiphenyl-5’-ol (5’-132). The PCB 132 metabolite profiles displayed inter-individual differences and depended on the PCB 132 atropisomer. Computational studies demonstrated that 3’-140 is formed via a 3,4-arene oxide intermediate. The second eluting atropisomer of PCB 132, first eluting atropisomer of 3’-140, and second eluting atropisomer of 5’-132 were enriched in all HLM incubations. Enantiomeric fractions of the PCB 132 metabolites differed only slightly between the single donor HLM preparations investigated. Reactive oxygen species and levels of dopamine and its metabolites were not significantly altered after a 24 h exposure of dopaminergic cells to pure PCB 132 atropisomers. These findings suggest that there are inter-individual differences in the atropselective biotransformation of PCB 132 to its metabolites in humans; however, the resulting atropisomeric enrichment of PCB 132 is unlikely to affect neurotoxic outcomes associated with the endpoints investigated in the study.

## INTRODUCTION

PCB congeners with a 2,3,6-chlorine substitution pattern on one phenyl ring, such as PCB 132, are important components of commercial PCB mixtures (Kania-Korwel *et al*., 2016a). Food is the major source of exposure to these and other PCB congeners (Schecter *et al*., 2010; Su *et al*., 2012; Voorspoels *et al*., 2008). For example, PCB 132 has been detected in fish species caught for human consumption (Wong *et al*., 2001). PCB 132 is also present in the indoor air of U.S. schools (Thomas *et al*., 2012), raising concerns about inhalation exposure of school children, teachers and staff to PCBs (Herrick *et al*., 2016). PCB 132 and structurally related PCB congeners are present in human blood (DeCaprio *et al*., 2005; Jursa *et al*., 2006; Whitcomb *et al*., 2005), breast milk (Bordajandi *et al*., 2008; Bucheli *et al*., 2006; Glausch *et al*., 1995) and postmortem human tissue samples (Chu *et al*., 2003). Like several other PCB congeners with a 2,3,6-chlorine substitution pattern, PCB 132 is axially chiral because it exists as two stable rotational isomers, or atropisomers, which are non-superimposable mirror images of each other (Lehmler *et al*., 2010).

Exposure to PCBs has been implicated in the etiology of neurodevelopmental and neurodegenerative disorders (Hatcher-Martin *et al*., 2012; Jones *et al*., 2008; Pessah *et al*., 2010). In particular, PCB congeners with two or more *ortho* chlorine substituents are neurotoxic and, for example, have been associated with behavioral and cognitive deficits in animal models (Caudle *et al*., 2006; Schantz *et al*., 1995; Wayman *et al*., 2012). Similarly, animal studies with hydroxylated PCB metabolites reported impairments in behavioral and locomotor activity in rats and mice (Haijima *et al*., 2017; Lesmana *et al*., 2014). Mechanistic studies suggest that these PCBs affect the dopaminergic system. *ortho*-Chlorinated PCBs inhibited dopamine uptake in rat synaptosomes (Mariussen *et al*., 2001) and decreased the dopamine content in cells in culture, possibly due to inhibition of dopamine synthesis (Seegal, 1996). 2,2’,3,4’,6-Pentachlorobiphenyl (PCB 91) and 2,2’,3,5’,6-pentachlorobiphenyl (PCB 95) decreased dopamine content in rat synaptosomes by inhibiting vesicular monoamine transporter (VMAT) (Bemis *et al*., 2004). In contrast, PCB 95 increased intracellular dopamine and decreased dopamine in the medium by down-regulating VMAT2 expression in PC12 cells (Enayah *et al*., 2018). Striatal dopamine levels in male mice exposed to PCB mixtures decreased due to a decrease in the expression of the dopamine transporter (DAT) and VMAT2 (Richardson *et al*., 2004). Other studies suggest that, in addition to their effects on the dopaminergic system, PCB neurotoxicity can be mediated by altered intracellular Ca^2+^ signaling and/or disruption of thyroid and sex hormone homeostasis (reviewed in: (Kodavanti *et al*., 2010; Mariussen *et al*., 2006; Pessah *et al*., 2010).

Neurotoxic PCBs are metabolized to potentially neurotoxic hydroxylated metabolites (OH-PCBs) in animal models and humans (Grimm *et al*., 2015; Kania-Korwel *et al*., 2016a). In general, PCB congeners with H-atoms in vicinal *meta* and *para* positions are readily metabolized by cytochrome P450 (P450) enzymes, whereas congeners without adjacent *para* and *meta* positions are metabolized more slowly (Grimm *et al*., 2015). The oxidation of PCBs by P450 enzymes occurs by direct insertion of an oxygen atom into an aromatic C-H bond or via an arene oxide intermediate (Forgue *et al*., 1982; Forgue *et al*., 1979; Preston *et al*., 1983). Several P450 isoforms, including CYP2A and CYP2B enzymes, are involved in the metabolism of *ortho* chlorinated PCB congeners, such as PCB 132, in different species (Lu *et al*., 2013; McGraw *et al*., 2006; Nagayoshi *et al*., 2018; Ohta *et al*., 2012; Uwimana *et al*., 2019; Waller *et al*., 1999; Warner *et al*., 2009). Hydroxylated and methyl sulfone metabolites of PCB 132 have been detected in human blood, breast milk and feces (Haraguchi *et al*., 2004; Haraguchi *et al*., 2005). Similar to animal studies (Norstroem *et al*., 2006), PCB 132 undergoes atropisomeric enrichment in humans (Bordajandi *et al*., 2008; Bucheli *et al*., 2006; Chu *et al*., 2003; Glausch *et al*., 1995; Zheng *et al*., 2016) due to atropselective metabolism of PCB 132.

Biotransformation studies reveal species differences in the atropselective metabolism of PCBs to chiral hydroxylated metabolites (Kania-Korwel *et al*., 2016a); however, only a few PCB metabolism studies in humans have been reported (Schnellmann *et al*., 1983; Uwimana *et al*., 2016, 2018; Wu *et al*., 2014). Because PCB 132 is present in the environment, displays atropisomeric enrichment in human samples, and represent a largely unexplored environmental and human health concern, the objective of this study was to investigate the atropselective metabolism of PCB 132 to OH-PCB metabolites by HLMs and assess if PCB 132 atropselectively affects neurotoxic outcomes in dopaminergic cells in culture.

## MATERIALS AND METHODS

### Chemicals and materials

Sources and purities of racemic PCB 132 (**Table S1**), PCB metabolite standards, chemicals, and other reagents; information regarding the HLM preparations; and a description of the separation and characterization of PCB 132 atropisomers are presented in the Supplementary Material. Gas chromatograms and the corresponding mass spectrum of PCB 132 are shown in **Figs. S1** and **S2**. The chemical structures and abbreviations of OH-PCB 132 metabolites are shown in **Fig. 1**. Rat-tail collagen was obtained from Sigma Aldrich (St Louis, MO). All other cell culture components such as fetal bovine serum (FBS), horse serum (HS), RPMI 1640 medium and penicillin-streptomycin were obtained from Life Technologies (Carlsbad, CA). 3,4-Dihydroxyphenylacetic acid (DOPAC) was obtained from Sigma Aldrich (St Louis, MO). Dihydroxyphenyl ethanol (DOPET) was obtained by reducing 3,4-dihydroxyphenylaldehyde (DOPAL) in a 10-fold excess of sodium borohydride (Jinsmaa *et al*., 2009). DOPAL was obtained synthetically from epinephrine by the method of Fellman (Anderson *et al*., 2011; Fellman, 1958).

**Fig. 1.**
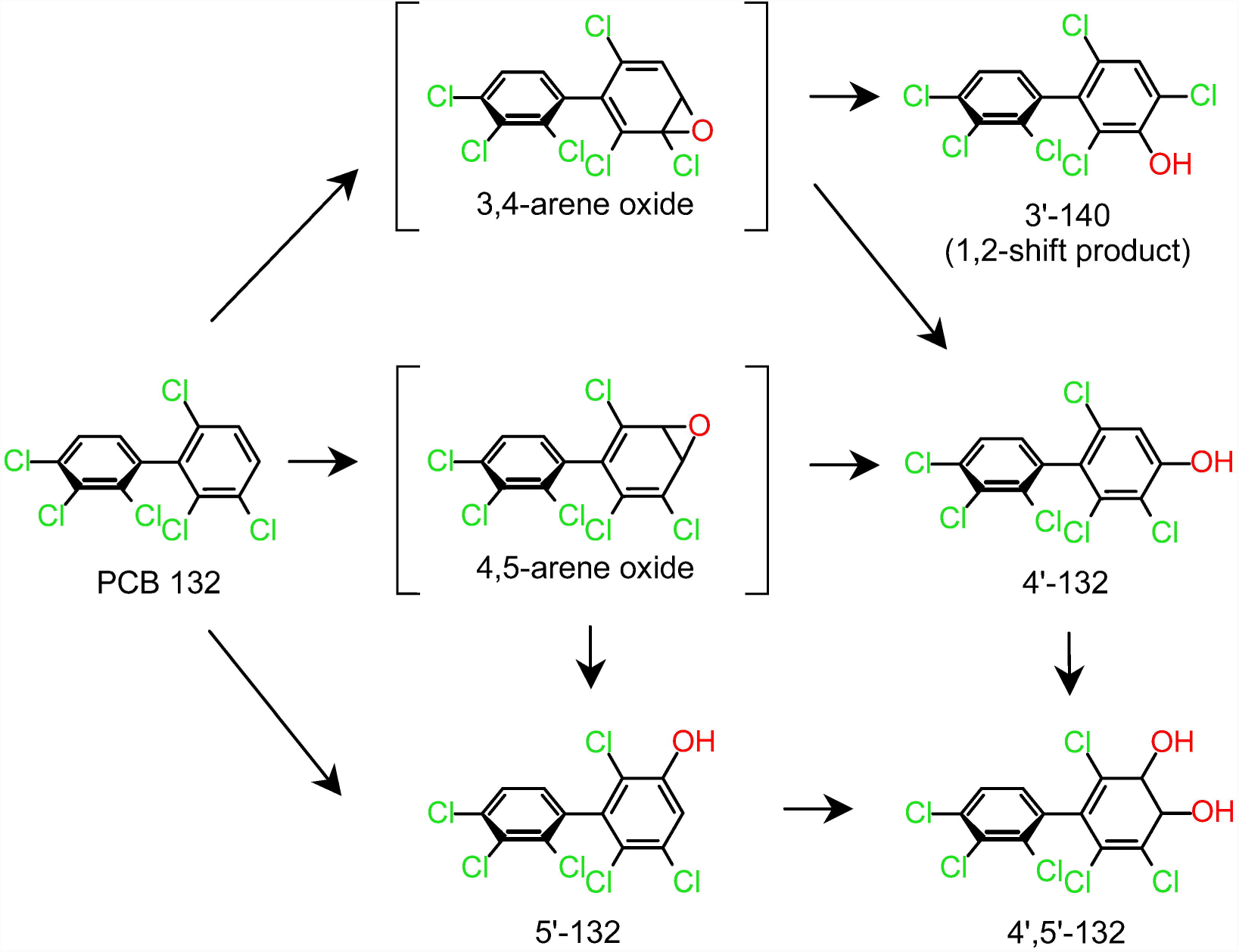
Proposed metabolism scheme showing the chemical structures of metabolites of PCB 132 identified in incubations with HLMs. Only one atropisomer of each metabolite is shown for clarity reasons. Abbreviations: 2,2’,3,3’,4,6’-hexachlorobiphenyl, PCB 132; 2,2’,3,4,4’,6’-hexachlorobiphenyl-3-ol, 3’-140; 2,2’,3,3’,4,6’-hexachlorobiphenyl-4’-ol, 4’-132; 2,2’,3,3’,4,6’-hexachlorobiphenyl-5’-ol, 5’-132; cytochrome P450 enzymes, P450.

### Microsomal incubations

The metabolism of PCB 132 was investigated in an incubation system containing sodium phosphate buffer (0.1 M, pH 7.4), magnesium chloride (3 mM), pooled human liver microsomes (pHLMs) or single donor HLMs (0.1 mg/mL), and NADPH (1 mM) (Kania-Korwel *et al*., 2011; Uwimana *et al*., 2017; Wu *et al*., 2011). The incubation mixtures were preincubated for 5 min, and PCB 132 (50 µM in DMSO; ≤ 0.5% *v/v*) was added to give a final volume of 2 mL. The mixtures were maintained for 10, 30 or 120 min at 37°C. Incubations with (−)-PCB 132, (+)-PCB132 or racemic PCB 132 (50 µM in DMSO; ≤ 0.5% *v/v*) were carried out analogously for 30 min at 37°C. The formation of PCB 132 metabolites was linear with time for up to 30 min. To terminate the enzymatic reaction, ice-cold sodium hydroxide (2 mL, 0.5 M) was added to each sample. Subsequently, the incubation mixtures were heated at 110 °C for 10 min. Phosphate buffer blanks and control incubations without PCB accompanied each microsomal preparation. The following incubations were performed in parallel with each experiment to control for the abiotic transformation of PCB 132: Incubations without NADPH, without microsomes, and with heat-inactivated microsomes. No metabolites were detected in the control samples. All incubations were performed in triplicate, if not stated otherwise.

### Extraction of PCB 132 and its hydroxylated metabolites

PCB 132 and its hydroxylated metabolites were extracted from the incubation mixtures as reported previously (Kania-Korwel *et al*., 2011; Uwimana *et al*., 2016; Wu *et al*., 2011). Briefly, PCB 117 (200 ng) and 4’-159 (68.5 ng) were added to each sample as surrogate recovery standards, followed by hydrochloric acid (6 M, 1 mL) and 2-propanol (5 mL). The samples were extracted with a hexane-MTBE mixture (1:1 v/v, 5 mL) and re-extracted once with hexane (3 mL). The organic layers were washed with an aqueous potassium chloride (1%, 4 mL), the organic phase was transferred to a new vial, and the KCl mixture was re-extracted with hexane (3 mL). The organic layers were evaporated to dryness under a gentle stream of nitrogen and reconstituted with hexane (1 mL). After derivatization with diazomethane (0.5 mL in diethyl ether) for approximately 16 h at 4 °C (Kania-Korwel *et al*., 2008), all extracts were subjected to sulfur and sulfuric acid clean-up steps before gas chromatographic analysis (Kania-Korwel *et al*., 2005; Kania-Korwel *et al*., 2007).

### Identification of PCB 132 metabolites

The identity of the hydroxylated PCB 132 metabolites was confirmed by high-resolution gas chromatography with time-of-flight mass spectrometry (GC-TOF/MS) (Uwimana *et al*., 2016, 2018). To obtain metabolite levels sufficient for GC-TOF/MS analysis, incubations were performed using the following experimental conditions: 50 µM racemic PCB 132, 0.3 mg/mL microsomal protein and 1 mM NADPH for 90 min at 37 °C. Metabolites were extracted and derivatized as described above before analysis on a Waters GCT Premier gas chromatograph (Waters Corporation, Milford, MA, USA) combined with a time-of-flight mass spectrometer in the High-Resolution Mass Spectrometry Facility of the University of Iowa (Iowa City, IA, USA) (Uwimana *et al*., 2016, 2018). Measurements were performed with and without heptacosafluorotributylamine as lock mass to determine the accurate mass of [M]^+^ and obtain mass spectra of the metabolites. PCB 132 metabolites were identified using the following criteria: The average relative retention times (RRT) of the OH-PCB 132 metabolites, calculated relative to PCB 51 as internal standard, were within 0.5% of the RRT of the authentic standard (European Commission, 2002); experimental accurate mass determinations were within 0.003 Da of the theoretical mass of [M]^+^; and the deviation between experimental and theoretical isotope pattern of [M]^+^ was < 20%.

### Quantification of PCB 132 metabolite levels

Levels of OH-PCB 132 metabolites in extracts were quantified as methylated derivatives on an Agilent 7890A gas chromatograph with a ^63^Ni-micro electron capture detector (µECD) and a SPB-1 capillary column (60 m length, 250 µm inner diameter, 0.25 µm film thickness; Supelco, St Louis, MO, USA) as reported earlier (Uwimana *et al*., 2016; Uwimana *et al*., 2017; Wu *et al*., 2013b). PCB 204 was added as internal standard (volume corrector) prior to analysis, and concentrations of OH-PCB 132 metabolites (as methylated derivatives) were determined using the internal standard method (Kania-Korwel *et al*., 2011; Wu *et al*., 2016; Wu *et al*., 2011). Levels of PCB and its metabolites were not adjusted for recovery to facilitate a comparison with earlier studies (Uwimana *et al*., 2016, 2018; Uwimana *et al*., 2019; Wu *et al*., 2016). The average RRTs of the metabolites, calculated relative to PCB 204, were within 0.5% of the RRT for the respective standard (European Commission, 2002). Metabolite levels and formation rates for all experiments described in the manuscript are summarized in the Supplementary Material (**Tables S2-S4**).

### Ring opening calculations

The ring opening reactions of selected PCB arene oxide intermediates were modeled using density functional theory at the M11/def2-SVP level of theory (Peverati *et al*., 2011; Weigend *et al*., 2005), coupled with the SMD aqueous continuum solvation model (Marenich *et al*., 2009), to assess if the formation of specific OH-PCB metabolites is energetically favored. The reactions were examined under neutral conditions, with two explicit water molecules complexed to the system to serve as a proton shuttle. To ascertain steric and electronic effects, calculations were carried out on arene oxides formed from 1,2,4-trichlorobenzene (TCB), 2,3,6-trichlorobiphenyl, PCB 91, PCB 95, PCB 132, and PCB 136. Transition states were characterized by normal mode analysis and confirmed via intrinsic reaction coordinate (IRC) calculations. All calculations were performed using Gaussian 16, Rev. A.01 (Frisch *et al*., 2016). Additional details including computational data are available in the Supplementary Material.

### Atropselective PCB analyses

Atropselective analyses were carried out with extracts from long-term incubations (*i.e.*, 5 or 50 µM PCB 132, 120 minutes at 37 °C, 0.5 mg/mL protein, and 0.5 mM NADPH) and extracts from the 30 min incubations described above. Hydroxylated metabolites were analyzed as methylated derivatives after derivatization with diazomethane. Analyses were performed on an Agilent 6890 gas chromatograph equipped with a µECD detector and a CP-Chirasil-Dex CB (CD) (25 m length, 250 µm inner diameter, 0.12 µm film thickness; Agilent, Santa Clara, CA, USA) or a ChiralDex G-TA (GTA) capillary column (30 m length, 250 µm inner diameter, 0.12 µm film thickness; Supelco, St Louis, MO, USA) (Kania-Korwel *et al*., 2011; Uwimana *et al*., 2017). The following temperature program was used for the atropselective analysis of PCB 132 and 5’-132 on a CD column: initial temperature was 50 °C, hold for 1 min, ramped at 10 °C/min to 160 °C, hold for 220 min, ramped at 20 °C/min to 200 °C, and hold for 10 min. To analyze 3’-140 on a GTA column, the temperature program was as follows: initial temperature was 50 °C, hold for 1 min, ramped at 10 °C/min to 150 °C, hold for 400 min, ramped at 10 °C/min to 180 °C, and hold for 40 min. The helium flow was 3 mL/min for all atropselective analyses. To facilitate a comparison with earlier studies (Kania-Korwel *et al*., 2016b; Uwimana *et al*., 2016; Uwimana *et al*., 2017; Wu *et al*., 2011), enantiomeric fractions (EFs) were calculated by the drop valley method (Asher *et al*., 2009) as EF = Area E_1_/(Area E_1_ + Area E_2_), with Area E_1_ and Area E_2_ denoting the peak area of the first (E_1_)and second (E_2_) eluting atropisomer on the respective column. All EF values are summarized in the Supplementary Material (**Table S6**).

### Quality Assurance and Quality Control

The response of the µECDs used in this study was linear for all analytes up to concentrations of 1,000 ng/mL (R^2^ ≥ 0.999). The recoveries of PCB 117 were 93 ± 13% (n = 120). Recoveries of 4’-159 could not be determined due to co-elution with 4’,5’-132. The limits of detection (LOD) of the PCB 132 metabolites were calculated from blank buffer samples as LOD = mean of blank samples + k × standard deviation of blank samples, where k is the student’s t value for a degree of freedom n-1 = 5 at 99% confidence level) (Kania-Korwel *et al*., 2011; Uwimana *et al*., 2016; Wu *et al*., 2016; Wu *et al*., 2011). The LODs were 0.03, 0.1 and 0.21 ng for 3’-140, 5’-132 and 4’-132, respectively. The background levels for 3’-140, 5’-132 and 4’-132 in control (DMSO) incubations with HLMs (n=6) were 0.17, 0.13 and 0.19 ng, respectively. The resolution of the atropisomers of PCB 132 and 5’-132 on the CD column was 1.05 and 2.02, respectively. The resolution of the atropisomers of 3’-140 on the GTA column was 1.18. The EF values of the racemic standards of PCB 132, 3’-140 and 5’-132 were 0.51 ± 0.01 (n=2), 0.50 ± 0.01 (n=3) and 0.49 ± 0.01 (n=3), respectively.

### Cell culture

PC12 cells were purchased from American Tissue Culture Collection (Manassas, VA, USA) and maintained in RPMI 1640 medium with 10% HS, 5% FBS, and 1% penicillin-streptomycin. Cells were kept in a humidified incubator at 37°C with 5% CO_2_ in a 100 mm^2^ tissue culture petri dish at a density of 5 × 10^6^ cells. For experiments, cells were plated in 6 well plates in 2 mL medium at a density of 1.5 × 10^6^ cells per well, or in a Collagen I coated 96-well plate at 50.5 × 10^3^ cells per well. N27 cells, a generous gift from Dr. Jau-Shyong at the National Institute of Environmental Health Sciences, were maintained in RPMI 1640 with 10% HS, and 1% penicillin-streptomycin. Cells were maintained in a humidified environment at 37°C with 5% CO_2_ in a 100 mm^2^ collagen-coated tissue culture petri dish at a density of 1 × 10^6^ cells. For experiments, cells were plated in collagen-coated 96-well plates at 5 × 10^3^ cells per well.

### Cell viability and production of reactive oxygen species (ROS)

Cell viability was determined in N27 and PC12 cells using the MTT (3-(4,5-dimethylthiazol-2-yl)-2,5-diphenyltetrazolium bromide) assay (van Meerloo *et al*., 2011). The formation of ROS was measured in N27 cells using the DCFDA and dihydroethidium (DHE) fluorescent probes (Chen *et al*., 2010; Speisky *et al*., 2009; Zhao *et al*., 2003). Details regarding these assays are provided in the Supplementary Material.

### Dopamine metabolite analysis using HPLC

Intracellular and extracellular dopamine metabolites were measured in PC12 lysate samples via an Agilent 1100 Series capillary HPLC system with an ESA Coulochem III coulometric electrochemical detector (Enayah *et al*., 2018). After seeding the cells in 6-well plates, they were kept under the same growth conditions for 48 h. Media was then removed and replaced with HEPES buffer (115 mM NaCl, 5.4 mM KCl, 1.8 mM CaCl_2_, 0.8 mM MgSO_4_, 5.5 mM glucose, 1 mM NaH_2_PO_4_, and 15 mM HEPES, pH = 7.4). The cells were then exposed to the PCB 132 atropisomers at 10 and 25 μM and incubated for 24 h at which point cell lysate was collected. After removing the extracellular buffer, 400 μL lysis buffer (10 mM potassium phosphate, 0.1% triton-X, pH = 7.4) was added to each well. Then a cell scraper was used to collect the cell lysate which was stored at −20°C until analysis. On the day of HPLC analysis, the samples were centrifuged at 10,000g for 10 min, and 3 μL of supernatant injected into the HPLC to measure intracellular dopamine and its metabolites. Separation was achieved with a Phenomenex Synergi C18 column (2 × 150 mm, 80 Å) using an isocratic mobile phase (50 mM citric acid, 1.8 mM sodium heptane sulfonate, 0.2% trifluoroacetic acid, 2% acetonitrile, pH = 3.0) at 200 μL/min. For electrochemical detection of catechol-containing compounds, the following setings were used: guard = +350, E1 = −150, and E2 = +200 mV, respectively. The amount of dopamine, DOPAC, and DOPET was determined by comparison to a standard curve and presented as pmol of analyte per mg of protein. Details regarding the measurement of dopamine and its metabolites in extracellular media are provided in the Supplementary Material. A bicinchoninic acid (BCA) assay was used on the PC12 lysate samples as previously described to attain the protein concentration (Walker, 1994).

## RESULTS

### Identification and quantification of PCB 132 metabolites in incubations with pHLMs

Several human biomonitoring studies have reported an atropisomeric enrichment of PCB 132 in human samples (**Table S8**) (Lehmler *et al*., 2010); however, the atropselective oxidation of this PCB congener to hydroxylated metabolites by HLMs has not been studied to date (Grimm *et al*., 2015; Kania-Korwel *et al*., 2016a). Our GC-TOF/MS analyses revealed the formation of three monohydroxylated and one dihydroxylated PCB metabolite in incubations of racemic PCB 132 with pHLMs (**Fig. 2**; **Figs. S4**-**S11**). The structure of these metabolites is shown in the simplified metabolism scheme in **Fig. 1**. Their identification was based on accurate mass determinations, the chlorine isotope patterns of their molecular ion (analyzed as methylated derivatives) and their fragmentation patterns; for additional discussion, see the Supplementary Material. PCB 132 was oxidized by pooled and individual donor HLMs in the *meta* position, with 5’-132 and 3’-140 being major metabolites (**Figs. 3-4**; **Table S2**). 3’-140 is a 1,2-shift product (Guroff *et al*., 1967) that is formed *via* an arene oxide intermediate, followed by a 1,2-shift of the 3’-chlorine substituent to the *para* position. 4’-132 was only a minor metabolite. 4’,5’-132 could not be quantified in this study due to co-elution with the recovery standard, 4’-159; however, this metabolite was only a minor metabolite. PCB 132 was not oxidized in the 2,3,4-chloro substituted ring.

**Fig. 2.**
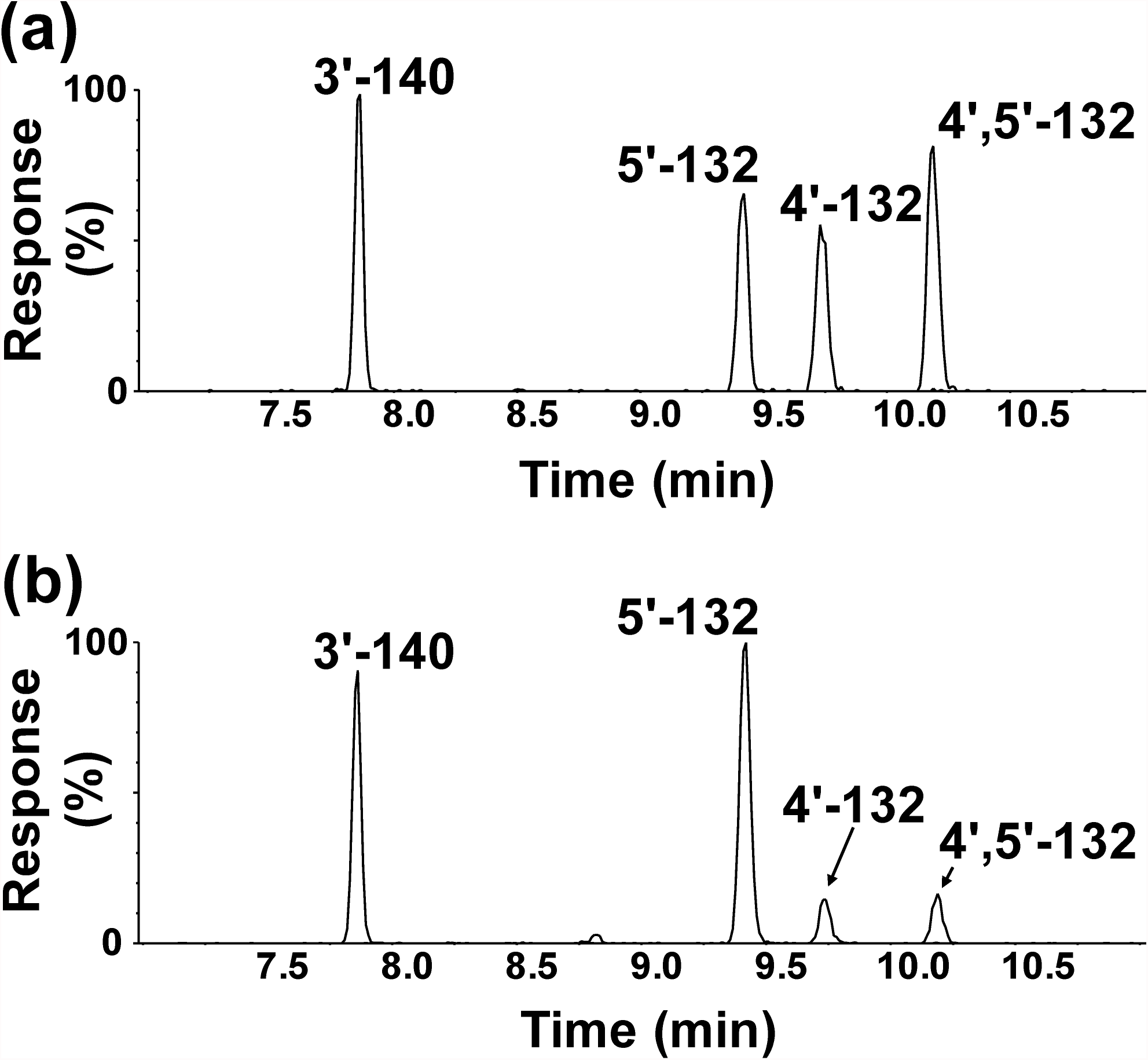
Three monohydroxylated (*m/z* 387.9) and one dihydroxylated (*m/z* 417.9) metabolite were identified in incubations of racemic PCB 132 with pHLMs. Representative gas chromatograms showing (a) the reference standard containing four hydroxylated PCB 132 metabolites and (b) an extract from a representative incubation of racemic PCB 132 with pHLMs. All metabolites were analyzed as the corresponding methylated derivatives. Incubations were carried out with 50 µM racemic PCB 132, 0.3 mg/mL microsomal protein and 1 mM NADPH for 90 min at 37 °C (Uwimana *et al*., 2016, 2018). Analyses were performed by GC-TOF/MS as described under Materials and Methods. The metabolites were identified based on their retention times relative to the authentic standard and their *m/z*. For the corresponding mass spectra, see **Figs. S4** to **S11**.

**Fig. 3.**
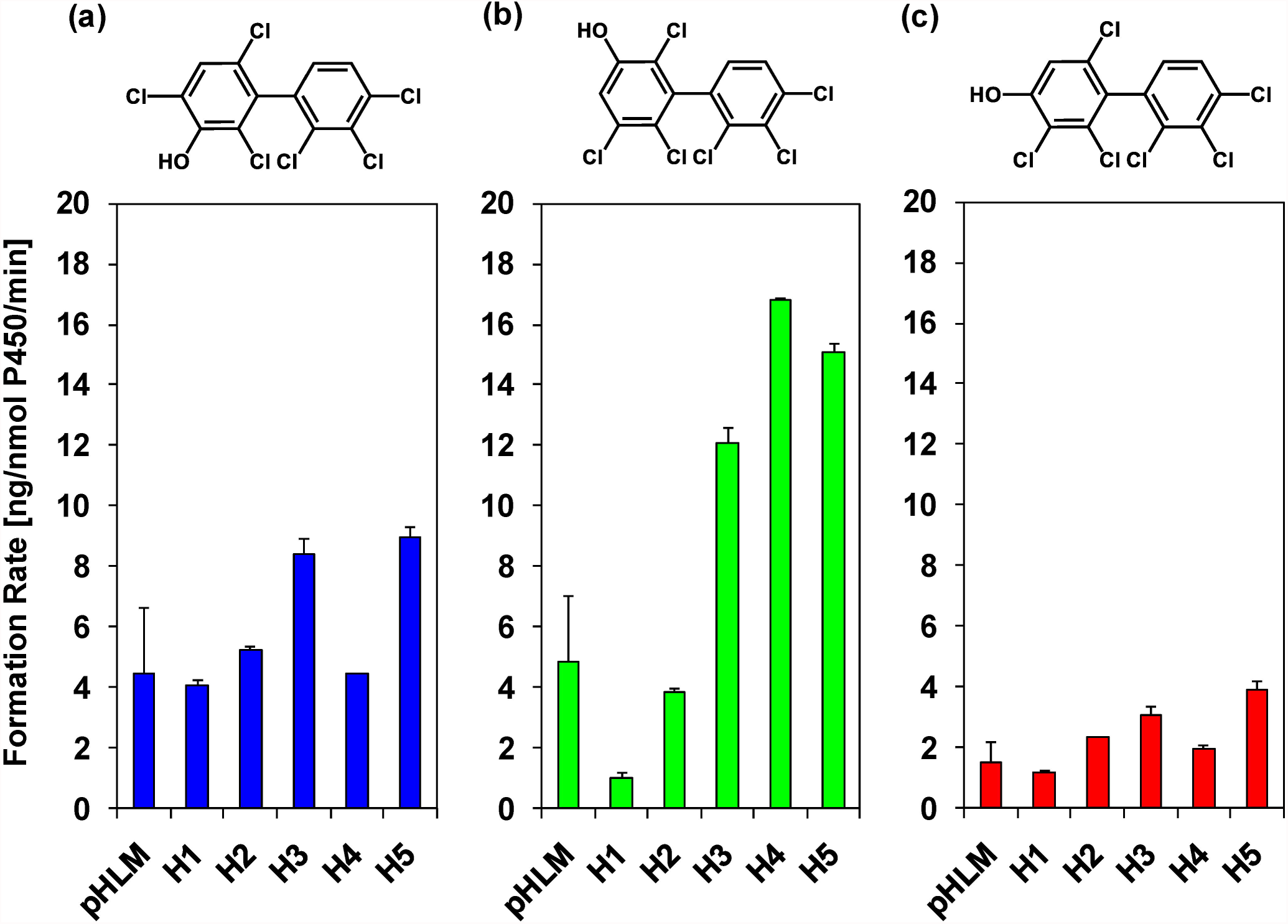
Formation rates of PCB 132 metabolites by different HLM preparations display inter-individual differences, with (a) the *meta* hydroxylated metabolites, 3’-140 (1,2-shift product), and (b) 5’-132, being major metabolites, and (c) the *para* hydroxylated metabolite, 4’-132, being a minor metabolite. Incubations were carried out with 50 µM racemic PCB 132, 0.1 mg/mL microsomal protein and 1 mM NADPH for 10 min at 37 °C (**Tables S3** and **S4**) as reported earlier (Uwimana *et al*., 2016, 2018). Extracts from the microsomal incubations were derivatized with diazomethane and analyzed by GC-μECD; see Materials and Methods for additional details. Data are presented as mean ± standard deviation, n = 3.

**Fig. 4.**
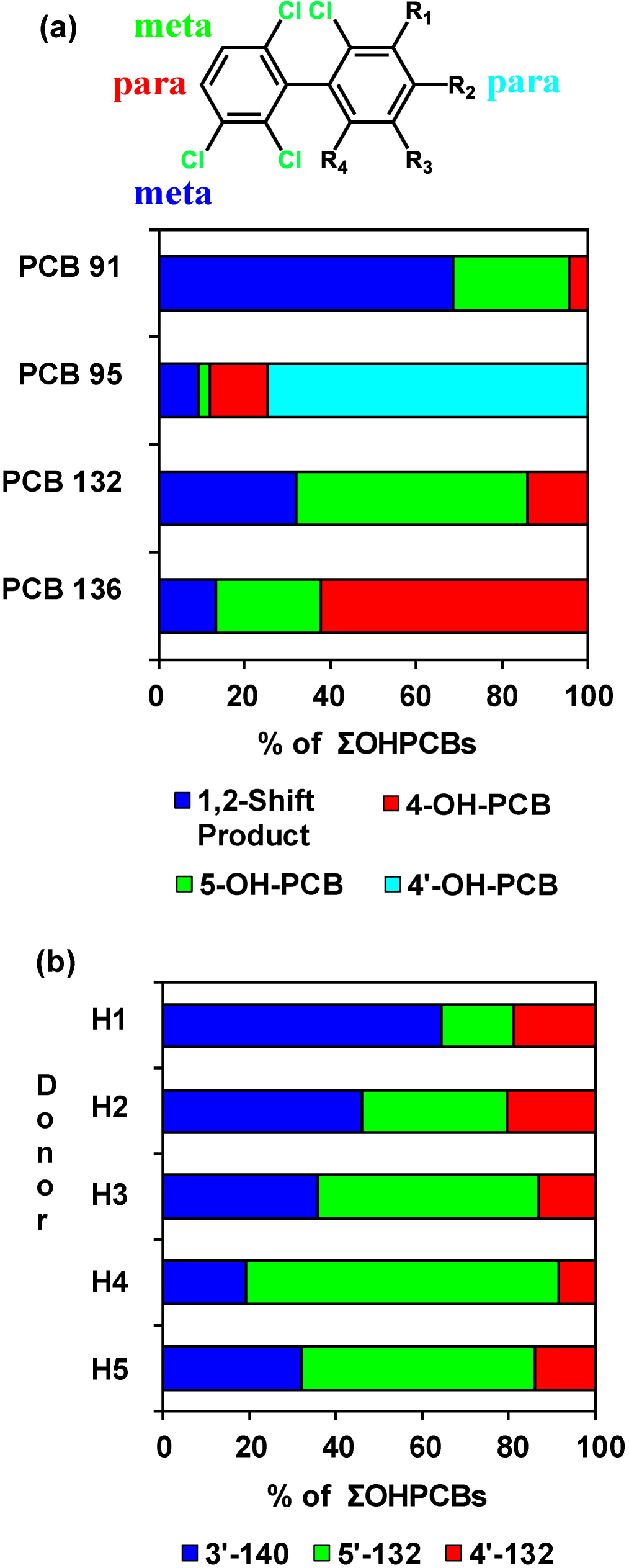
The OH-PCB metabolite profile of PCB 132 formed by pHLMs is (a) distinctively different from the published metabolite profiles of structurally related PCB congeners (*i.e.,* PCB 91, PCB 95 and PCB 136) and (b) shows considerable inter-individual variability. (a) Bar diagram comparing the profiles of hydroxylated metabolites of PCB 91, PCB 95, PCB 132 and PCB 136 formed in incubations with pHLMs. PCB 91 (Uwimana *et al*., 2018) and PCB 132 (this study) are preferentially oxidized in *meta* position, including the formation of 1,2-shift products, whereas PCB 95 (Uwimana *et al*., 2016) and PCB 136 (Wu *et al*., 2014) are hydroxylated in *para* position. In the case of PCB 95, the oxidation occurs preferentially in the *para* position of the lower chlorinated 2,5-dichlorophenyl ring (Uwimana *et al*., 2016). Incubations were carried out with 50 µM PCB, 0.1 mg/mL microsomal protein and 1 mM NADPH for 5 min (PCB 91 and PCB 95) or 10 min (PCB 132 and PCB 136) at 37 °C using the same pHLM preparation. (b) Bar diagram showing inter-individual differences in profiles of hydroxylated metabolites of PCB 132 formed in incubations with different single donor HLM preparations. Incubations were carried out with 50 µM PCB 132, 0.1 mg/mL microsomal protein and 1 mM NADPH for 10 min at 37 °C (Uwimana *et al*., 2016, 2018). Extracts from the microsomal incubations were derivatized with diazomethane and analyzed by GC-μECD; see Materials and Methods for additional details.

### Metabolism of PCB 132 in comparison to structurally related PCBs

As shown in **Fig. 4a**, the PCB 132 metabolite profiles formed by HLMs differs considerably from the profiles of structurally related PCB congeners with a 2,3,6-trichlorophenyl group (*i.e.*, PCB 91, PCB 95 and 2,2’,3,3’,6,6’-hexachlorobiphenyl [PCB 136]). HLMs oxidized PCB 132 in the *meta* position to yield the 1,2-shift product, 3’-140, and 5’-132. PCB 91, which is structurally similar to PCB 132, was also primarily metabolized to a 1,2-shift product by HLMs (Uwimana *et al*., 2018). PCB 95 was preferentially oxidized by HLMs in the *para* position with the lower chlorinated, 2,5-dichloro substituted phenyl ring, and only traces of a 1,2-shift product were formed by different HLM preparations (Uwimana *et al*., 2016). PCB 136 was metabolized in both the *meta* and *para* position to yield comparable levels of 5’-136 and 4’-136 (Schnellmann *et al*., 1983; Wu *et al*., 2014).

To assess if the congener-specific differences in OH-PCB metabolite profiles are due to different arene oxide intermediates or different substitution patterns in the second phenyl ring of chiral PCB congeners, the energetics of the 3,4-arene oxide ring opening reactions were determined for 1,2,4-trichlorobenzene, 2,3,6-trichlorobiphenyl, PCB 91, PCB 95, PCB 132, and PCB 136 (**Fig. 5**; **Fig. S12** and **Table S5**). Also, complete ring-opening pathways were computed for the 3,4-, 4,5-, and 5,6-arene oxides of a single atropisomer of PCB 132 and PCB 136, another environmentally relevant PCB congener (**Table S5**). Because the 3,4-, 4,5- and 5,6-arene oxides may open in two directions, six pathways were computed for both PCB 132 and PCB 136.

**Fig. 5.**
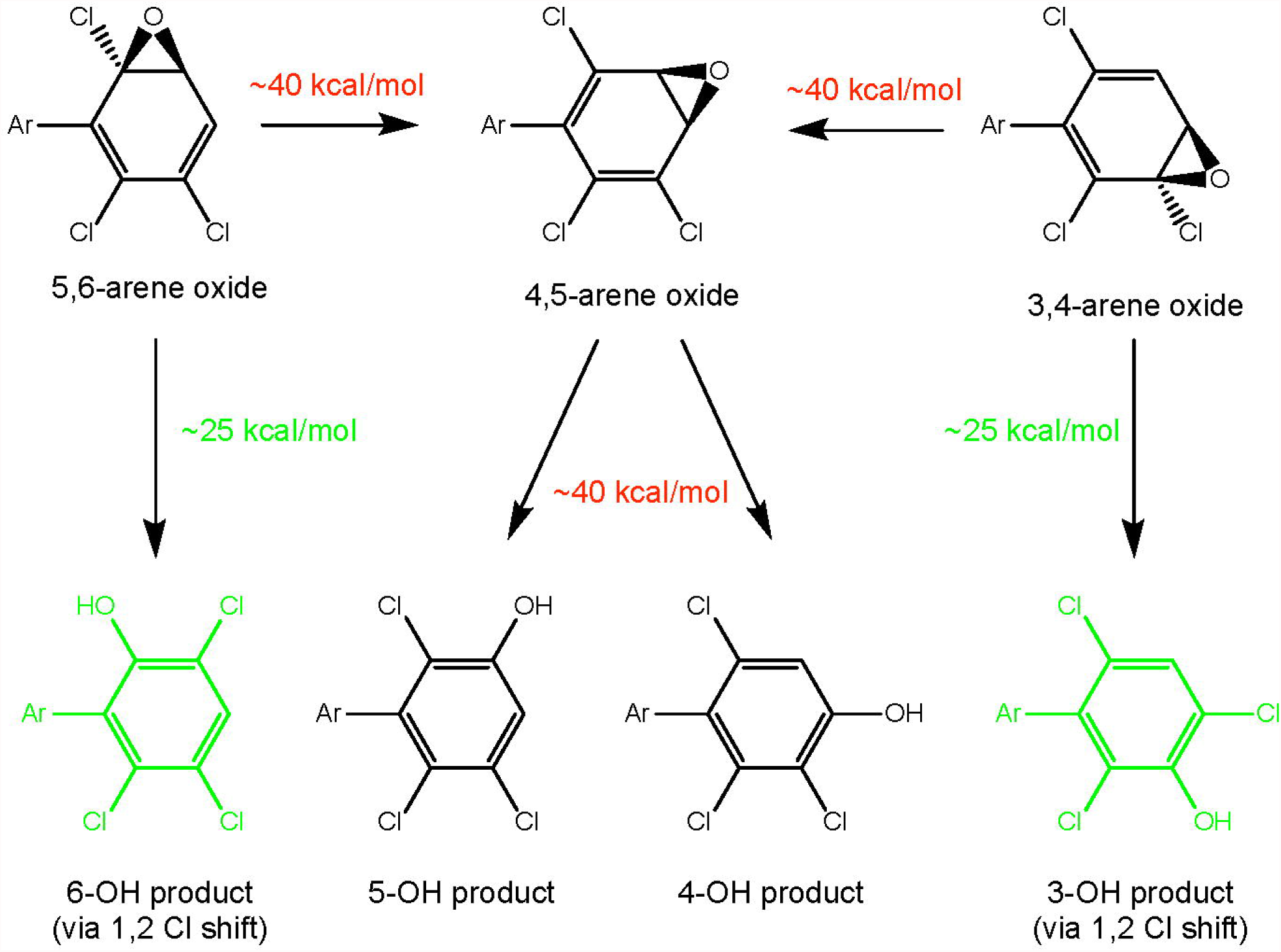
The formation of 1,2 chlorine shift metabolites via arene oxide intermediates of chiral PCBs is energetically favored. Energies shown are free energies of activation at the M11/def2-SVP level of theory with the SMD aqueous continuum solvation model. Ar: 2’,4’-dichlorophenyl for PCB 91; 2’,5’-dichlorophenyl for PCB 95; 2,3,4-trichlorophenyl for PCB 132; and 2’,3’,6’-dichlorophenyl for PCB 136.

In the case of the 3,4- and 5,6-arene oxides, the arene oxide opening towards the C-Cl group and the 1,2-shift of the chlorine were concerted, with barriers to the opening of approximately 25 kcal/mol (**Fig. S12**). The proton transfer step to form the phenol and restore aromaticity proceeded with a negligible barrier and was essentially spontaneous. Partial pathways opening toward C-Cl were further explored for the 3,4-arene oxides of 1,2,4-trichlorobenzene, 2,3,6-trichlorobiphenyl, PCB 91 and PCB 95. The presence or absence of the substituent aryl ring had no effect on the computed mechanism or energetics for an opening toward C-Cl, nor did the substitution pattern on the aryl substituent.

The energy barrier to arene oxide opening was approximately 40 kcal/mol for both arene oxides when the 3,4- and 5,6-arene oxides open towards the aromatic C-H bond (**Fig. 5**). For both the 3,4- and 5,6-arene oxides, this mode of opening leads to formation of the 4,5-arene oxide as confirmed by IRC calculations, and not the formation of OH-PCB. Because both possible openings of the 4,5-arene oxide also proceed towards C-H, it is not surprising that the barriers for 4,5-arene oxide opening were also approximately 40 kcal/mol. Here, IRC calculations revealed concerted 1,2-hydride shifts analogous to the 1,2-chloride shift observed for 3,4-arene oxide opening toward C-Cl. Overall, the computational results demonstrate that the formation of 1,2-shift products, such as 3’-140, is energetically favored for all PCB arene oxides investigated.

### Inter-individual differences in PCB 132 metabolism by HLMs

In experiments with single donor HLMs, the formation rates of the monohydroxylated PCB 132 metabolites displayed inter-individual variation (**Figs. 3** and **4b**; **Tables S3** and **S4**). The sum of OH-PCBs (ΣOH-PCBs) formed by single donor HLM preparations followed the following rank order: donor H5 > H4 ∼ H3 > H2 ∼ pHLM > H1. Notably, levels of ΣOH-PCB were 4.5-fold higher in incubations with HLMs from donors H5 *vs.* H1. The rate of 5’-132 formation differed 16-fold for incubations with HLM preparations from donors H1 *vs.* H4. The rate of 3’-140 and 4’-132 formation differed 2.2 and 3.3-fold, respectively, for incubations with HLM preparations from donors H1 *vs.* H5. As a consequence, the OH-PCB metabolite profiles differed across the HLM preparations investigated (**Fig. 4b**). Briefly, 3’-140 and 5’-132 were formed in a 1:1 ratio by pHLMs. In incubations with HLMs from donors H1 and H2, the 3’-140 to 5’-132 ratios were 3.9:1 and 1.4:1 for donors H1 and H2, respectively. In contrast, 5’-132, and not 3’-140, were major in incubations with HLMs from donors H3, H4 and H5 (3’-140 to 5’-132 ratios were 0.7:1, 0.3:1 and 0.6:1 for donors H3, H4, and H5, respectively). Although only a small number of single donor HLM preparations were investigated in this and other studies, the profiles of OH-PCB metabolite formed in the liver likely display considerable variability in humans.

### Atropisomeric enrichment of PCB 132

Because chiral PCBs undergo atropisomeric enrichment *in vivo* and affect toxic endpoints in an atropselective manner (Kania-Korwel *et al*., 2016a; Lehmler *et al*., 2010), the enrichment of PCB 132 atropisomers in incubations of racemic PCB 132 with HLMs was investigated with atropselective gas chromatography. E_2_-PCB 132 (EF = 0.39), which corresponds to (+)-PCB 132 (Haglund *et al*., 1996), was enriched in experiments with low PCB 132 concentration (5 µM) and long incubation time (120 min) (**Fig. 6**, **Table S6**). EF values were near racemic in incubations with higher PCB 132 concentrations (50 µM) because the large amount of racemic PCB 132 masked the atropselective depletion of one PCB 132 atropisomer over the other (Uwimana *et al*., 2016, 2018).

**Fig. 6.**
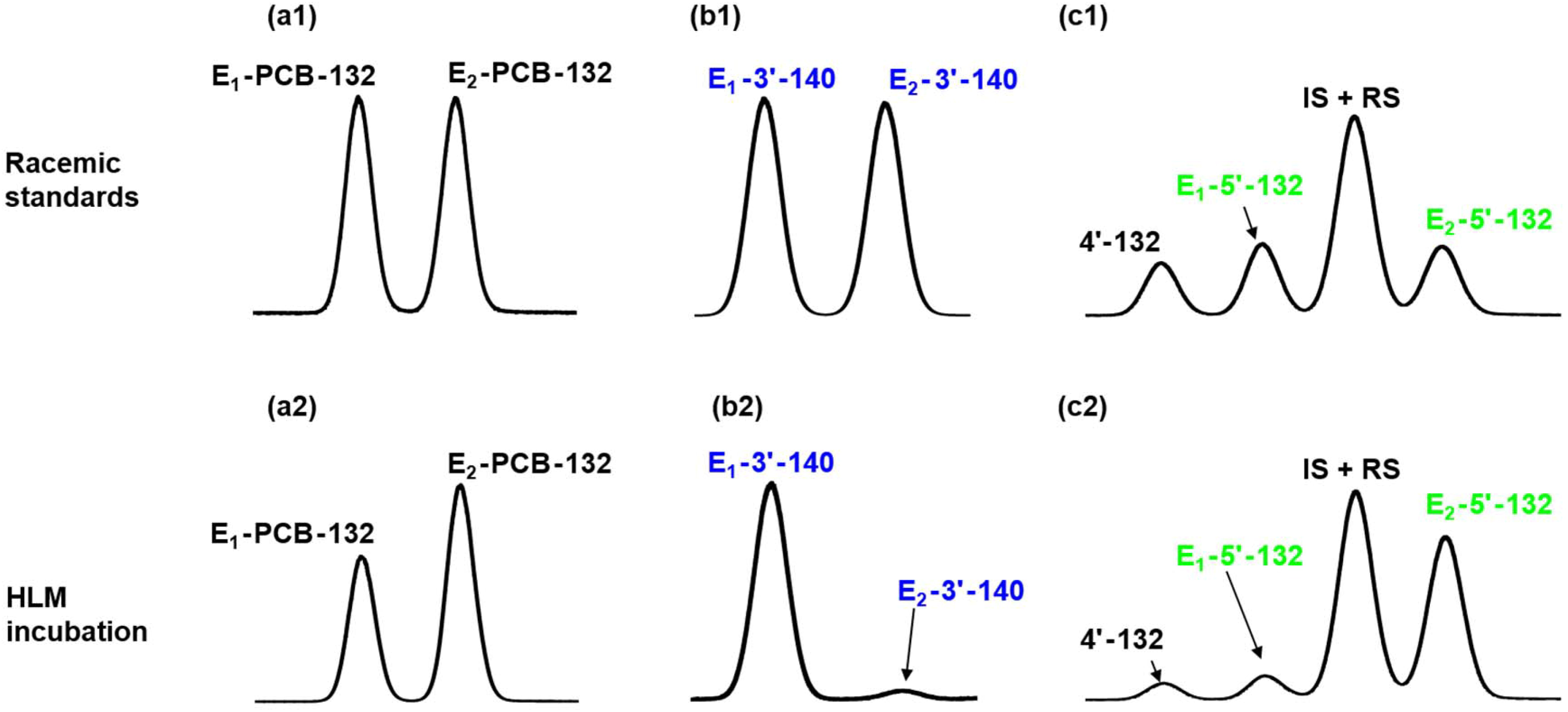
Atropselective gas chromatographic analysis revealed the atropselective metabolism of racemic PCB 132 to 5’-132 and 3’-140 in incubations with pHLMs. Representative gas chromatograms of racemic standards (top panels) *vs.* PCB 132, 5’-132 and 3’-140 in incubations of racemic PCB 132 with HLMs (bottom panels) show a depletion of E_1_-PCB 132 (panels a1 *vs.* a2) and atropselective formation of E_1_-3’-140 (panels b1 *vs.* b2) and E_2_-5’-132 (panels c1 *vs.* c2). To assess the atropselective depletion of PCB 132, incubations were carried out with 5 µM PCB 132, 0.5 mg/mL microsomal protein (pHLMs only) and 0.5 mM NADPH for 120 min at 37 °C. To study the atropselective formation of the PCB 132 metabolites, incubations were carried out with 50 µM racemic PCB 132, 0.1 mg/mL microsomal protein and 1 mM NADPH for 30 min at 37 °C (donor H3 shown here; see Fig. 7 for results from incubations using other HLM preparations) (Uwimana *et al*., 2016, 2018). Metabolites were analyzed as the corresponding methylated derivatives after derivatization with diazomethane. Atropselective analyses of 3’-140 were performed with a GTA column at 150 °C, and atropselective analyses of PCB 132 and 5’-132 were carried out with a CD column at 160 °C (Kania-Korwel *et al*., 2011).

### Atropselective formation of OH-PCB 132 metabolites

Several studies have reported the atropselective formation of chiral OH-PCB metabolites, both from chiral and prochiral PCBs, in *in vitro* and *in vivo* studies (Kania-Korwel *et al*., 2016a; Lehmler *et al*., 2010; Uwimana *et al*., 2017). Consistent with these earlier studies, we observed the atropselective formation of different OH-PCB 132 metabolites in incubations with HLMs. Specifically, the atropselective analysis on the GTA column revealed the atropselective formation of E_1_-3’-140, with EF values > 0.8 (range: 0.84 to 0.95) (**Figs. 6-7**). E_2_-5’-132 was significantly enriched with EF values < 0.2 (range: 0.12 to 0.18) in incubations with HLMs. As reported previously, the atropisomers of 4’-132 could not be resolved on any of the chiral columns used (Kania-Korwel *et al*., 2011).

**Fig. 7.**
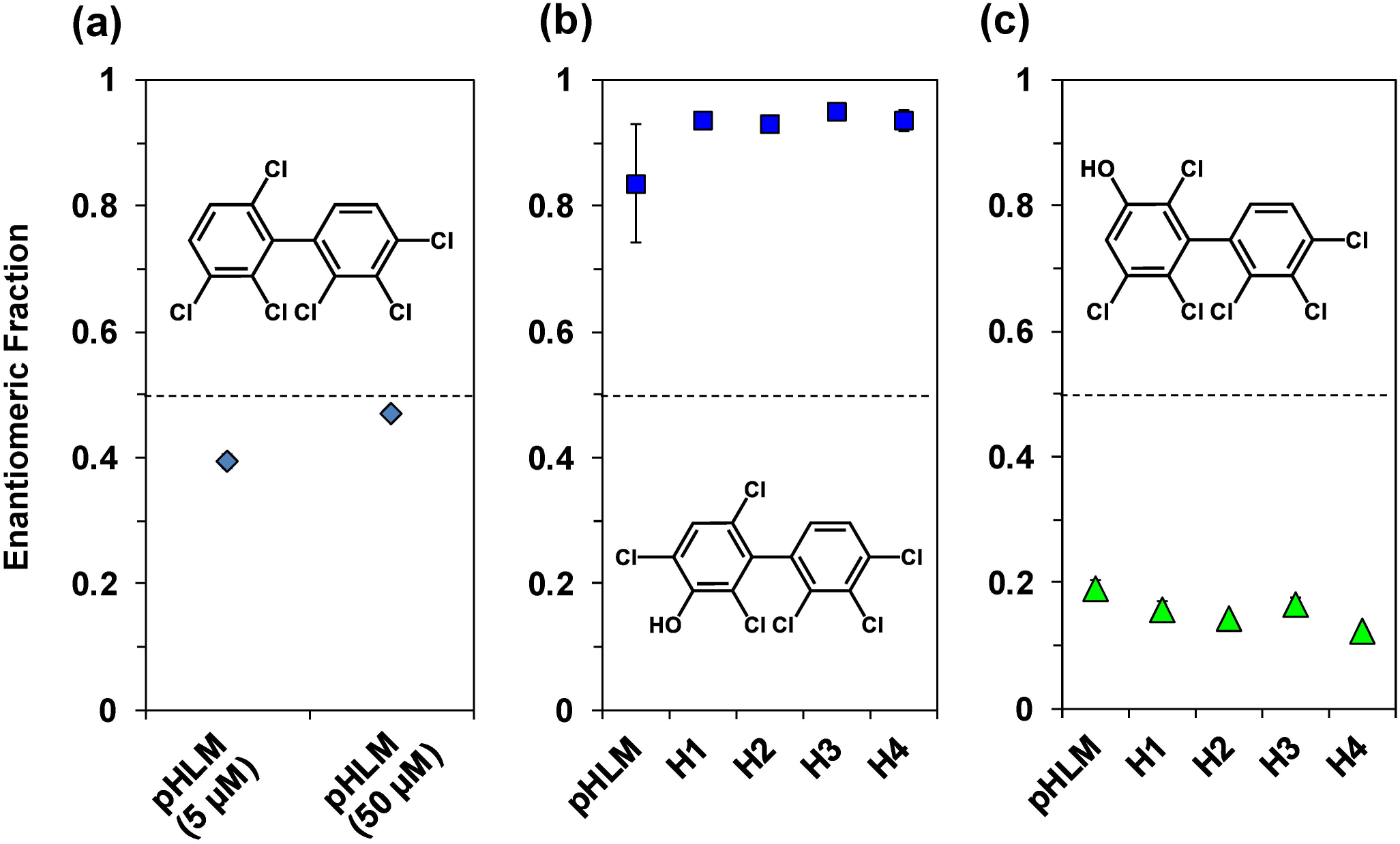
Enantiomeric fractions (EFs) of (a) parent PCB 132, (b) 3’-140 and (c) 5’-132 reveal only small inter-individual differences in the atropselective formation of both OH-PCB metabolites. (a) To assess the atropselective depletion of PCB 132, incubations were carried out with 5 µM or 50 µM PCB 132, 0.5 mg/mL microsomal protein (using pHLMs only), 0.5 mM NADPH for 120 min at 37 °C (see Fig. 6 for a representative chromatogram). To study the atropselective formation of (b) 3’-140 and (c) 5’-132, microsomal incubations were carried out with 50 µM racemic PCB 132, 0.1 mg/mL microsomal protein, 1 mM NADPH for 30 min at 37 °C (Uwimana *et al*., 2016, 2018). Metabolites were analyzed as the corresponding methylated derivatives. Atropselective analyses of 3’-140 were performed with a GTA column at 150 °C, and atropselective analyses of PCB 132 and 5’-132 were carried out with a CD column at 160°C (Kania-Korwel *et al*., 2011). EF values could not be determined in incubations with HLMs from donor H5 due to the low metabolite levels. Data are presented as mean ± standard deviation, n = 3. The dotted line indicates the EF values of the racemic standards.

To determine which of E_1_-OH-PCB and E_2_-OH-PCB atropisomer is formed from (−)- or (+)-PCB 132, we investigated the metabolism of (−)-, (±)-, and (+)-PCB 132 by pHLMs (**Fig. 8**, **Table S7**). The atropselective analysis showed that E_1_-5’-132 is formed from (+)-PCB 132. Conversely, E_1_-3’-140 and E_2_-5’-132 are formed from (−)-PCB 132. The ∑OH-PCBs formed from (−)-PCB 132 was 3-times the ∑OH-PCBs formed from (+)-PCB 132 (**Table S7**). A more complex picture emerges when individual OH-PCB 132 metabolites are analyzed (**Figs. 8e-g**). The levels of both major metabolites, 3’-140 and 5’-132, decreased in the order of (−)-PCB 132 > (±)-PCB 132 > (+)-PCB 132. In contrast, levels of the minor metabolite, 4’-132, decreased in the reverse order. As a consequence, the OH-PCB 132 metabolite profiles formed by HLMs differ considerably for incubations with (−)-, (±)-, and (+)-PCB 132 (**Fig. 8h**).

**Fig. 8.**
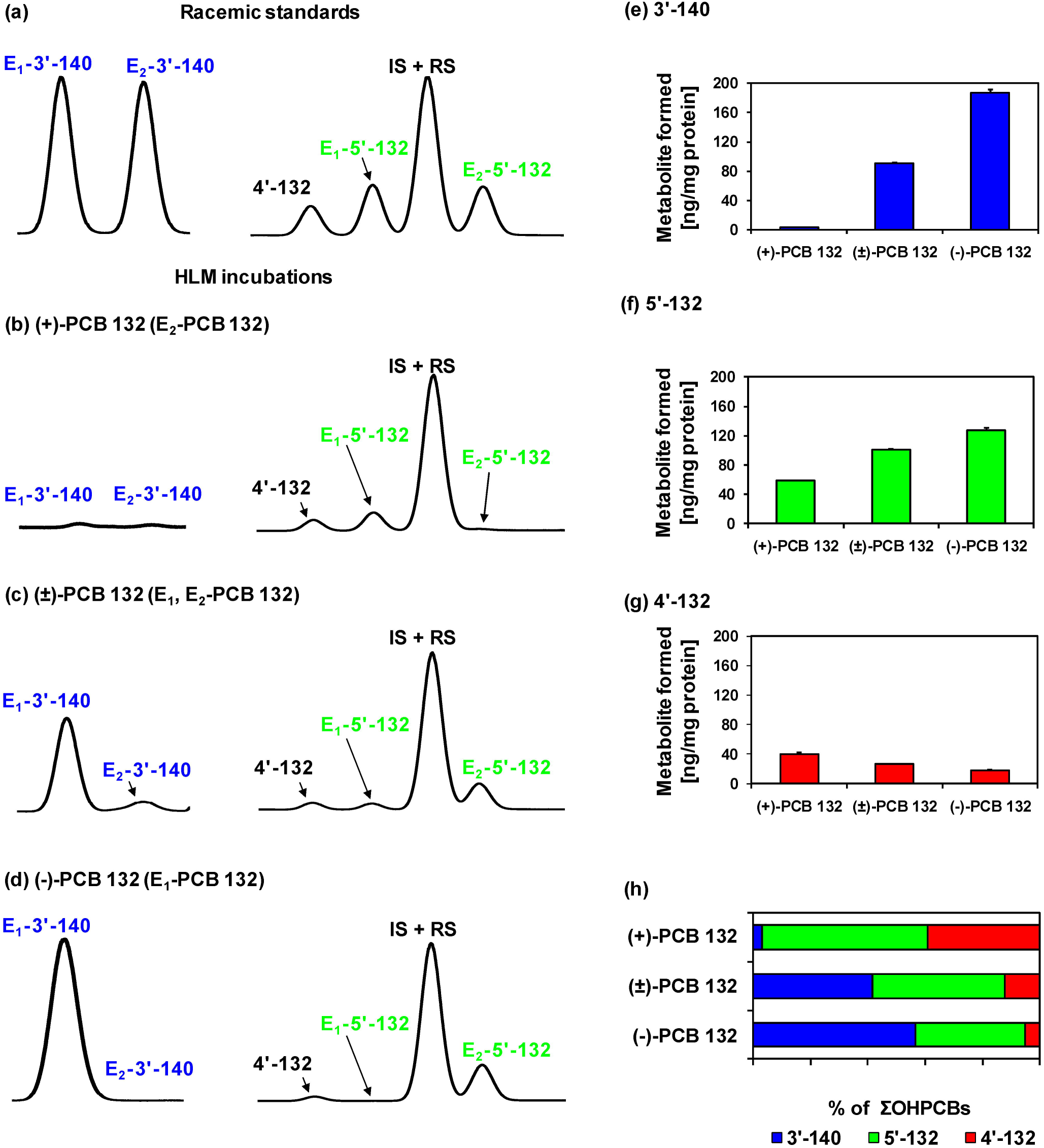
Comparison of representative gas chromatograms of (a) racemic OH-PCB metabolite standards with OH-PCB metabolites formed in incubations of (b) (+)-PCB 132, (c) (±)-PCB 132 or (d) (−)-PCB 132 with pHLMs reveals that E_1_-5’-132 is formed from (+)-PCB 132 and E_1_-3’-140, and E_2_-5’-132 are formed from (−)-PCB. Moreover, (e) 3’-140 and (f) 5’-132, but not (g) 4’-132 are formed more rapidly from (−)-PCB 132 than (+)-PCB 132, resulting (h) in distinct OH-PCB metabolite profiles formed from (+)-, (±)-, and (−)-PCB 132 in incubations with pHLMs. Incubations were carried out with 50 µM (+)-PCB 132, racemic PCB 132 or (−)-PCB 132, 0.1 mg/mL microsomal protein and 1 mM NADPH for 30 min at 37 °C (Uwimana *et al*., 2016). Metabolites were analyzed as the corresponding methylated derivatives after derivatization with diazomethane. Atropselective analyses of 3’-140 were performed with a GTA column at 150 °C, and atropselective analyses of PCB 132 and 5’-132 were carried out with a CD column at 160°C. 4’-132 was not resolved on any of the columns used in this study (Kania-Korwel *et al*., 2011). Data are presented as mean ± standard deviation, n = 3.

### Cytotoxicity and oxidative stress of PCB 132 atropisomers

It is currently unknown if pure PCB atropisomers differentially cause cytotoxicity or atropselectively increase the production of reactive oxygen species in dopaminergic cells. In our study, PCB 132 atropisomers did not elicit overt cytotoxicity in immortalized dopaminergic cell lines as determined by MTT analysis (**Fig. S13**). Both (+)-PCB 132 and (−)-PCB 132 exerted measurable toxicity to N27 cells only at the highest dose (100 μM) with a 24 h exposure. Analogously, PC12 cells showed no significant cell loss following PCB 132 atropisomer exposure. We observed a slight increase in overall reactive oxygen species and hydrogen peroxide, measured with DCFDA and DHE, respectively, after 24 h exposure of N27 cells to PCB 132 atropisomers; however, these changes did not reach statistical significance (**Fig. S14**). Similarly, no statistically significant, time-dependent changes in the production of reactive oxygen species were observed at earlier time intervals (data not shown). There were also no significant differences in the effects of both PCB 132 atropisomers on dopaminergic cellular toxicity and levels of reactive oxygen species.

### Effect of PCB 132 on levels of dopamine and its metabolites

PCBs, including chiral PCB 95, have been shown to affect the dopaminergic system, including by altering the expression of dopamine receptors and by changing dopamine levels (Bavithra *et al*., 2012; Enayah *et al*., 2018; Seegal *et al*., 2002); however, it is unknown if PCBs atropselectively affect the dopaminergic system and dopamine metabolism. PC12 cells were exposed to PCB atropisomers for 24 h, and intracellular dopamine and its metabolites were measured via HPLC analysis. Dopamine is converted by monoamine oxidase to DOPAL which is further metabolized by aldehyde dehydrogenase to DOPAC or by aldehyde reductase to DOPET. Intracellular levels of dopamine, DOPAC, and DOPET showed no significant differences from control. (**Fig. 9**). Extracellular levels of dopamine and DOPAC was also measured in media with similar results (**Fig. S15**).

**Fig. 9.**
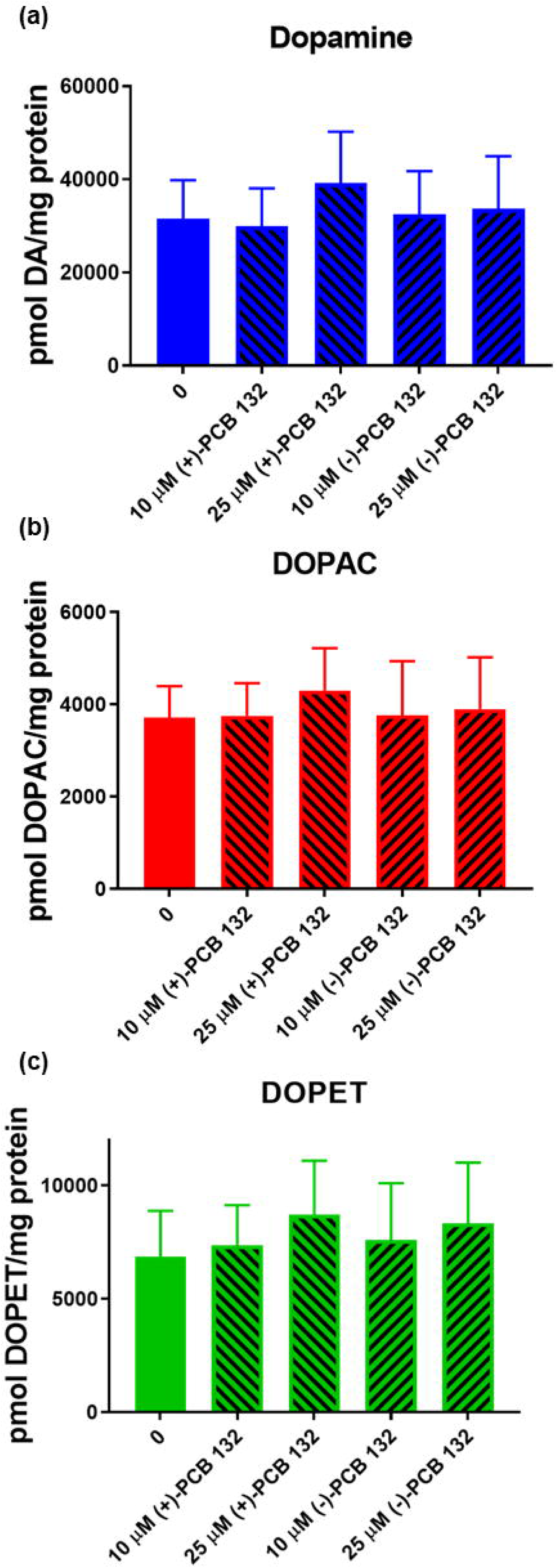
Exposure of PC12 cells for 24 h to (+)- or (−)-PCB 132 did not significantly alter intracellular levels of (a) dopamine, (b) 3,4-dihydroxyphenylacetic acid (DOPAC), or (c) 3,4-dihydroxyphenyl ethanol (DOPET). Intracellular measurements of dopamine and its metabolites were performed by HPLC analysis with electrochemical detection. Data are represented as mean ± standard error, n = 4.

## DISCUSSION

PCB 132 is readily metabolized by HLMs to a 1,2-shift product, 3’-140, and a second *meta* hydroxylated metabolite, 5’-132. Several structurally related PCB congeners with multiple *ortho* chlorine substituents are also metabolized by HLMs to OH-PCBs (Nagayoshi *et al*., 2018; Schnellmann *et al*., 1983; Wu *et al*., 2014). For example, PCB 91 is primarily metabolized to a 1,2-shift product by HLMs (Uwimana *et al*., 2018), whereas PCB 95 and PCB 136 are oxidized by HLMs with different regioselectivity (Schnellmann *et al*., 1983; Uwimana *et al*., 2016; Wu *et al*., 2014). Based on published structure-metabolism relationships (Grimm *et al*., 2015), these congener-specific differences in their metabolism are likely due to the presence of a *para* chlorine substituent in PCB 91 and PCB 132, but not PCB 95 and PCB 136. The formation of a 1,2-shift product indicates that similar to PCB 91, PCB 132 is oxidized by human P450 enzymes to an arene oxide intermediate that subsequently undergoes a 1,2 chlorine shift to 3’-140. Importantly, different OH-PCB profiles have been reported for the metabolism of these PCB congeners in rodent models used to study the neurotoxicity of PCBs (Kania-Korwel *et al*., 2017; Schantz *et al*., 1997; Wayman *et al*., 2012). For example, 5’-132, and not 3’-140, was the major metabolite formed from PCB 132 in experiments with rat liver microsomes (Kania-Korwel *et al*., 2011), recombinant rat CYP2B1 (Lu *et al*., 2013), and precision-cut mouse liver tissue slices (Wu *et al*., 2013a). 1,2-Shift products were only minor metabolites of PCB 132 and other chiral PCBs in *in vitro* and *in vivo* studies in rodent models (Kania-Korwel *et al*., 2011; Lu *et al*., 2013; Wu *et al*., 2013a).

The preferential formation of different PCB arene oxide intermediates in rodent vs. humans is one explanation for the species differences in the oxidation of PCB 132 and structurally related PCB congeners. We employed a computational approach to study the energetics of the ring opening of the three possible arene oxides on the 2,3,6-trichloro-substituted phenyl ring to explore this possibility. Independent of the position of the arene oxide, our calculations demonstrated that (1) the ring opening leading to a 1,2-shift of a chlorine substituents is energetically favorable over a ring opening toward C-H and (2) the presence of the second aromatic ring does not perturb the energetics of the arene oxide ring opening. Based on our computational results, the 1,2-shift products of PCB 132 (this study) and PCB 91 (Uwimana *et al*., 2018) are formed via a 3,4-arene oxide intermediate which opens toward C-Cl to form the 1,2-shift metabolite. Moreover, our computational results suggest that the formation of *para*-substituted metabolites of PCB 95 and PCB 136 is not merely due to the energetically favored opening of the 3,4- or 4,5-arene oxide intermediates. Instead, congener-specific differences in the steric or electronic interaction of both PCB congeners with P450 isoforms involved in their metabolism (Uwimana *et al*., 2019) likely affect the ring opening of the putative arene oxide intermediates to OH-PCBs. In contrast, the formation of OH-PCB metabolites of PCB 132 and related PCB congeners in rodents may or may not involve an arene oxide intermediate. Indeed, the *meta* oxidation of PCB 52 by rat CYP2B1 does not involve an arene oxide but occurs via the insertion of an oxygen atom into the aromatic C-H bond (Preston *et al*., 1983).

In addition to congener and species-dependent differences in the OH-PCB metabolite profiles, we observed considerable inter-individual variability in the formation of PCB 132 metabolites by HLM preparations from different donors. For example, differences as high as 16-fold were found in the formation of 5’-132 by different HLM preparations. Similarly, we reported inter-individual differences in the formation of OH-PCB metabolites of PCB 91 and PCB 95 with the same HLM preparations (Uwimana *et al*., 2016, 2018). This observation is not surprising considering the well-documented variability of P450 enzyme activities among humans (Guengerich, 2015). As we reported recently, both CYP2A6 and CYP2B6 are involved in the metabolism of PCB 132 (Uwimana *et al*., 2019). CYP2A6 primarily formed 3’-140, with 5’-132 and 4’-132 being minor metabolites. CYP2B6 primarily yielded 5’-132, with the 1,2-shift product being a minor metabolite. Differences in the levels of CYP2A6 and CYP2B6 in the HLM preparations investigated are one explanation for the inter-individual differences in the OH-PCB formation rates in this and our earlier studies. Moreover, both CYP2A6 and CYP2B6 are highly polymorphic enzymes (PharmVar, 2012, 2013). As has been shown for active site mutants of CYP2B1 (Waller *et al*., 1999), the rat ortholog of CYP2B6, polymorphisms can alter the regioselectivity of the oxidation of PCBs.

The biotransformation of PCB 132 by P450 enzymes results in non-racemic chiral signatures of both the parent PCB and its metabolites in environmental samples and mammals, including humans (Kania-Korwel *et al*., 2016b; Lehmler *et al*., 2010). Importantly, the enrichment of PCB atropisomers observed in *in vitro* studies typically predicts the atropisomeric enrichment in animal studies. (+)-PCB 132 was enriched in *in vitro* metabolism studies with precision-cut mouse liver tissue slices (Wu *et al*., 2013a), rat liver microsomes (Kania-Korwel *et al*., 2011), and recombinant rat CYP2B1 and human CYP2B6 (Lu *et al*., 2013; Warner *et al*., 2009). Consistent with these *in vitro* metabolism studies, (+)-PCB 132 was enriched in female mice (Kania-Korwel *et al*., 2010; Milanowski *et al*., 2010) and male Wistar rats (Norstrom *et al*., 2006). (−)-PCB 132 had a shorter half-life in a disposition study in mice (Kania-Korwel *et al*., 2010). In rats, levels of methyl sulfone metabolites of PCB 132 were higher in (−)-PCB 132 compared (+)-PCB 132 exposed animals, suggesting that (−)-PCB 132 is more rapidly metabolized in rats (Norstrom *et al*., 2006). Analogous to the laboratory studies, the preferential oxidation of (−)-PCB 132 by HLMs observed in this study is consistent with the enrichment of (+)-PCB 132 found in most human samples (**Table S8**). However, other factors may also contribute to the atropisomeric enrichment of PCB 132 in humans. For example, we cannot dismiss the possibility that exposure to atropisomerically enriched PCB 132 *via* the diet also contributes to the atropisomeric enrichment of PCB 132 in humans (Harrad *et al*., 2006; Vetter, 2016).

In addition to the parent PCB, and consistent with earlier studies reporting the atropselective formation of chiral OH-PCB metabolites from PCBs (Kania-Korwel *et al*., 2016a; Lehmler *et al*., 2010; Uwimana *et al*., 2017), we observed the atropselective formation of OH-PCB 132 metabolites in incubations with HLMs. Specifically, we noted the atropselective formation of E_1_-3’-140. This 3’-140 atropisomer was also enriched in experiments with rat liver microsomes (Kania-Korwel *et al*., 2011). E_2_-5’-132 was significantly enriched in incubations with HLMs. Similarly, metabolism of racemic PCB 132 resulted in an enrichment of E_2_-5’-132 in incubations with rat liver microsomes (Kania-Korwel *et al*., 2011) and mouse liver tissue slices (Wu *et al*., 2013a). In contrast, rat CYP2B1 metabolized PCB 132 preferentially to E_1_-5’-132, which indicates the involvement of other P450 isoforms in the atropselective oxidation of PCB 132 in incubations with rat liver microsomes (Lu *et al*., 2013). The atropselective formation of OH-PCBs by HLMs is important in the context of several findings from laboratory studies: Chiral OH-PCBs present in the developing brain of mice exposed developmentally to racemic PCBs (Kania-Korwel *et al*., 2017). Moreover, the position of the OH-group has a profound effect on the interaction of OH-PCBs with cellular targets implicated in PCB neurotoxicity (Kodavanti *et al*., 2003; Niknam *et al*., 2013). It is therefore likely that inter-individual differences in (1) PCB and OH-PCB levels and (2) chiral signatures influence toxic outcomes in humans.

Several studies have shown that PCB atropisomers atropselectively affect endpoints relevant to PCB neurotoxicity, including altered calcium homeostasis, estrogen receptor activation, and ROS, linked to PCB developmental neurotoxicity (Chai *et al*., 2016; Feng *et al*., 2017; Lehmler *et al*., 2005; Pencikova *et al*., 2018; Yang *et al*., 2014). In our preliminary assessment of the effects of PCB 132 atropisomers on endpoint implicated in PCB neurotoxicity, we saw increased general ROS levels, which are known to damage lipids and proteins, at sub-toxic PCB 132 concentrations in dopaminergic cells in culture; however, this effect did not reach statistical significance at the PCB 132 concentrations investigated. Exposure of rats to a PCB mixture containing racemic PCB 132 leads to dopaminergic neurodegeneration (Lee *et al*., 2012). Both PCB 132 atropisomers did not alter dopamine metabolism in our cell culture studies. A detailed structure-activity investigation in the same cell line showed that PCB congeners with *ortho* substituents result in a decrease in intracellular, total dopamine levels, but that chlorine substituents in *meta* position counteract this effect. The lack of an effect of PCB 132, which contains a *meta* chlorine substituent in both phenyl rings, on cellular dopamine metabolism is therefore not entirely surprising. It is noteworthy that exposure of PC12 cells to racemic PCB 95 decreases dopamine concentrations and increases levels of toxic dopamine metabolites (Enayah *et al*., 2018). A systematic study of other PCB atropisomers is therefore warranted to assess if PCBs atropselectively cause oxidative stress and alter dopamine homeostasis in the brain and, ultimately, if the atropselective metabolism of PCBs plays a currently unexplored role in PCB-mediated neurotoxicity.

## SUPPLEMENTARY DATA DESCRIPTION

Supplementary data, including source of chemicals and other materials, identification of PCB 132 metabolites by GC-TOF/MS, description and summary of cell viability studies and ROS measurements, purity determination of PCB 132, quantification of levels and formation rates of OH-PCB 132 metabolites, summary of relative free energies for arene oxide ring opening, enantiomeric fractions of PCB 132 and its metabolites, summary of chiral PCB 132 signatures in human samples, and representative mass spectra of PCB 132 metabolites are available online at http://toxsci.oxfordjournals.org/.

## ACKNOWLEDGMENTS

We thank Drs. S. Joshi, S. Vyas, and Y. Song (University of Iowa) for synthesizing the PCB standards and Mr. V. Parcel from the University of Iowa HRMS Facility for help with the GC-TOF/MS.

