## Supplemental material for "Atropselective Oxidation of 2,2’,3,3’,4,6’-Hexachlorobiphenyl (PCB 132) to Hydroxylated Metabolites by Human Liver Microsomes and Its Implications for PCB 132 Neurotoxicity"

Running Head: Atropselective metabolism and toxicity of PCB 132

Corresponding Author:

Dr. Hans-Joachim Lehmler

The University of Iowa

Department of Occupational and Environmental Health

University of Iowa Research Park, B164 MTF

Iowa City, IA 52242-5000

**Table of Content**

| Sources of chemicals and other materials | **S4** |
| --- | --- |
| HPLC separation of PCB 132 atropisomers | **S5** |
| Identification of PCB 132 metabolites by GC-TOF/MS | **S6** |
| Cell viability studies | **S7** |
| Detection of reactive oxygen species in N27 cells | **S8** |
| Dopamine metabolite analysis in PC12 media |  |
| **Table S1.** Purity determination of PCB 132 using GC-MS | **S10** |
| **Table S2.** Quantification of OH-PCB 132 metabolites (ng/mg microsomal protein) in incubations with different human liver microsome (HLM) preparations. | **S11** |
| **Table S3.** Rate of PCB 132 metabolite formation (ng OH-PCB/nmol P450/min) by different human liver microsome (HLM) preparations. | **S12** |
| **Table S4.** Rate of PCB 132 metabolite formation (ng OH-PCB/mg microsomal protein/min) by different human liver microsome (HLM) preparations. | **S13** |
| **Table S5.** Summary of relative free energies for epoxide ring opening. | **S14** |
| **Table S6.** Enantiomeric fractions (EFs) of PCB 132, 3'-140 and 5'-132 in incubations with human liver microsomes reveal the atropselective formation of the second eluting atropisomer of 5'-132 (E_2_-5'-132) and nearly enantiospecific formation the first eluting atropisomer of 3'-140 (E_1_-3'-140). | **S15** |
| **Table S7.** PCB 132 metabolite levels in incubations of individual atropisomers and racemic PCB 132 with pHLMs. | **S16** |
| **Table S8.** Summary of chiral PCB 132 signatures in human samples. | **S17** |
| **Figure S1.** GC-MS chromatogram and mass spectrum of the test compound, PCB 132, analyzed in the TIC mode. | **S18** |
| **Figure S2.** Enlarged chromatogram showing minor impurities (retention times 23.403, 23.483, 24.662, 25.320, and 28.793 min) present in PCB 132 by GC-MS (TIC). | **S19** |
| **Figure S3.** Enantioselective separation of (a) (-)-PCB 132, (b) racemic PCB 132 and (c) (+)-PCB 132 on a GC-µECD equipped with a ChiralsilDex-CD column. (-)-PCB 132 (EF = 0.97) and (+)-PCB 132 (EF = 0.05) correspond to E_1_-PCB 132 and E_2_-PCB 132, respectively. | **S20** |
| **Figure S4.** Mass spectrum of the authentic standard of methylated 3'-140 (RT 7.87 min; RRT 1.809). | **S21** |
| **Figure S5.** Mass spectrum of the authentic standard of methylated 5'-132 (RT 9.44 min; RRT 2.170). | **S22** |
| **Figure S6.** Mass spectrum of the authentic standard of methylated 4'-132 (RT 9.76 min; RRT 2.244). | **S23** |
| **Figure S7.** Mass spectrum of the authentic standard of methylated 4',5'-132 (RT 10.19 min; RRT 2.343). | **S24** |
| **Figure S8.** Mass spectrum of 3'-140 (RT 7.86 min; RRT 1.803) formed in a representative incubation with pooled human liver microsomes (analyzed as methylated derivative). | **S25** |
| **Figure S9.** Mass spectrum of 5'-132 (RT 9.42 min; RRT 2.161) formed in a representative incubation with pooled human liver microsomes (analyzed as methylated derivative). | **S26** |
| **Figure S10.** Mass spectrum of 4'-132 (RT 9.74 min; RRT 2.234) formed in a representative incubation with pooled human liver microsomes (analyzed as methylated derivative). | **S27** |
| **Figure S11.** Mass spectrum of 4',5'-132 (RT 10.19 min; RRT 2.337) formed in a representative incubation with pooled human liver microsomes (analyzed as methylated derivative). | **S28** |
| **Figure S12.** Ring-opening pathways determined for PCB 132 at the M11/def2-SVP + SMD level of theory. | **S29** |
| **Figure S13.** Exposure to PCB 132 atropisomers has little effect on the viability of N27 and PC12 cell, as determined with the MTT assay. | **S30** |
| **Figure S14.** Production of reactive oxygen species in N27 cells after 24 h exposure to PCB 132 atropisomers. | **S31** |
| **Figure S15.** Levels of (a) dopamine and (b) 3,4-dihydroxyphenylacetic acid (DOPAC) in media are not significantly changed following exposure of PC12 cells to PCB 132 atropisomers. | **S32** |
| Computational data | **S33** |
| References | **S71** |

**Sources of chemicals and other materials**

3'-Methoxy-2,2',3,4,4',6'-hexachlorobiphenyl (methylated derivative of 3'-140), 2,2',3,3',4,6'-hexachlorobiphenyl-4'-ol (4'-132), 2,2',3,3',4,6'-hexachlorobiphenyl-5'-ol (5'-132) and 4',5'-dimethoxy-2,2',3,3',4,6'-hexachlorobiphenyl (dimethylated derivative of 4',5'-132) were synthesized as described previously (Joshi *et al.*, 2011; Kania-Korwel *et al.*, 2008). The chemical structures and the abbreviations of the OH-PCB metabolites are shown in **Fig. 1** in the manuscript. PCB 132, 2,2',4,6'-tetrachlorobiphenyl (PCB 51; internal standard); 2,3,4',5,6-pentachlorobiphenyl (PCB 117; recovery standard), 2,2',3,4,4',5,6,6'-octachlorobiphenyl (PCB 204) and 2,3,3',4,5,5'-hexachlorobiphenyl-4'-ol (4'-159; recovery standard) were purchased from AccuStandard (New Haven, CT, USA). The purity of PCB 132 used in microsomal incubations was 99.8 % (**Table S1**), as determined on an Agilent 6890 chromatograph coupled with an Agilent 5975 Inert Mass Selective Detector (Agilent Technologies, CA, USA) operated in electron ionization mode and equipped with a SLB-5MS capillary column (30 m length, 250 µm inner diameter, 0.25 µm film thickness; Supelco. St, Louis, MO, USA) as described earlier (Holland *et al.*, 2017; Li *et al.*, 2018a; Uwimana *et al.*, 2018). The gas chromatogram and mass spectrum of PCB 132 are shown in **Figs. S1** and **S2**, respectively.

Solutions of diazomethane in diethyl ether for the derivatization of hydroxylated PCB metabolites to methoxylated PCB derivatives were synthesized from N-methyl-N-nitroso-p-toluenesulfonamide (Diazald) using an Aldrich mini Diazald apparatus (Milwaukee, WI, USA).

Dihydroethidium (DHE), dopamine, paraquat, and β-nicotinamide adenine dinucleotide 2'-phosphate reduced tetrasodium salt hydrate (NADPH) were purchased from Sigma-Aldrich (St. Louis, MO, USA). 2’,7’-Dichlorodihydrofluoroscein diacetate (DCFDA) and 3-(4,5-dimethylthiazol-2-yl)-2,5-diphenyltetrazolium bromide (MTT) were purchased from Invitrogen (Carlsbad, CA, USA). Salts for the preparation of buffers and reagent solutions (*e.g.*, sodium phosphate dibasic, sodium phosphate monobasic, magnesium chloride, tetrabutylammonium sulfite and sodium sulfite), acids (*e.g.*, concentrated sulfuric and hydrochloric acid) and solvents, including dimethyl sulfoxide (DMSO) and pesticide grade solvents, were obtained from Fisher Scientific (Pittsburgh, PA, USA). Ultrapure water (18 mΩ) was obtained from a Milli-Q Academic water purification system and was used to prepare all aqueous solutions.

Pooled human liver microsomes (pHLMs, catalog number H0620, pool of 50, mixed gender; lot numbers 0910398 or 1410013) and human liver microsomes from individual female donors (H1: catalog number H0531, lot number 0710185; H2, catalog number H0832, lot number 0910174; H3: catalog number H0779, lot number 0910130; H4: catalog number H0788, lot number 0910139; H5: catalog number H0444, lot number 0710042) were provided by Xenotech (Lenexa, KS, USA). The same microsomal preparations have been used in our earlier metabolism studies with PCB 91 and PCB 95 (Uwimana *et al.*, 2016, 2018).

**HPLC separation of PCB 132 atropisomers**

Semi-preparative separation of PCB 132 atropisomers was performed on a Shimadzu high performance-liquid chromatography system using two serially connected enantioselective Nucleodex β-PM columns (silica-based permethylated β-cyclodextrin, 200 mm length, 4 mm inner diameter, 5 μm particle size; Macherey-Nagel, Düren, Germany) with MeOH:H_2_O (85:15, v/v) as mobile phase and a flow rate of 0.23 mL/min at 12 °C (Haglund, 1996; Li *et al.*, 2018b). Briefly, 20 µL of a saturated solution of racemic PCB 132 in HPLC grade methanol was repeatedly injected into the HPLC system and fractions of both PCB 132 atropisomers were collected. Corresponding fractions were pooled, and the mobile phase was evaporated under a gentle stream of nitrogen. The residue was re-dissolved in 1 mL of hexane, the PCB 132 atropisomers were eluted with hexane through a glass pipette packed with 2 g of silica gel, and the solvent was removed under a gentle stream of nitrogen to give the pure (-)-PCB 132 and (+)-PCB 132 (**Fig. S3**). The overall purities were 99.97 % for (-)-PCB 132 and 99.98 % for (+)-PCB 132 (determined by GC-MS and based on the relative peak area).

**Identification of PCB 132 metabolites by GC-TOF/MS**

GC-TOF/MS analysis was used to identify the PCB 132 metabolites formed in incubations with HLMs. We observed the formation of three monohydroxylated and one dihydroxylated PCB metabolite in incubations of racemic PCB 132 with pHLMs (**Fig. 2**). The structure of these metabolites is shown in the simplified metabolism scheme shown in **Fig. 1** in the manuscript. Their identification was based on accurate mass determinations and the chlorine isotope patterns of their molecular ion (analyzed as methylated derivatives); see **Figs. S4** to **S11** for mass spectra of the authentic standards and metabolites extracted from incubations with HLMs). Moreover, the fragmentation patterns of the methylated monohydroxylated PCB 132 metabolites showed characteristic fragments, including [M-CH_3_]^+^, [M-CH_3_-CO] ^+^, [M-CH_3_- Cl]^+^, and [M-CH_3_-CO-Cl_2_]^+^, that are characteristic of *meta*- or *para*-methoxylated hexachlorobiphenyls (**Figs. S4** to **S6** and **S8** to **S10**, respectively). Similar fragmentation patterns have been reported by earlier studies for *meta* or *para*, but not *ortho* substituted derivatives of derivatives of monohydroxylated PCBs (Bergman *et al.*, 1995; Jansson *et al.*, 1974; Joshi *et al.*, 2011; Li *et al.*, 2009).

The mass spectrum of the dihydroxylated PCB 132 metabolite (analyzed as methylated derivative) showed characteristic fragments, for example [M-CH_3_]^+^, [M-CH_3_CO]^+^, [M-C_2_H_6_CO]^+^, [M-CH_3_COCl]^+^, [M-C_2_H_6_COCl]^+^, [M-CH_3_COCl-HCl]^+^ and [M-C_2_H_6_(CO)_2_Cl]^+^ (**Figs. S7** and **S11**). These fragments have been previously observed for 4',5'-132 (as methylated derivative) (Haraguchi *et al.*, 2004; Joshi *et al.*, 2011). The identity of this dihydroxylated PCB 132 metabolite, as well as the monohydroxylated metabolites discussed above, were further confirmed by comparing the relative retention times and mass spectral information to an authentic standard of each analyte (**Fig. 2**) and by GC-μECD analysis (see Materials and Methods for additional details).

**Cell viability studies**

To determine a non-toxic concentration of the PCB 132 atropisomers in N27 and PC12 cells, we employed the MTT assay as previously described (van Meerloo *et al.*, 2011). MTT is a tetrazolium salt that is reduced by metabolically active mitochondria to a purple, water insoluble formazan salt. The PCB congeners were dissolved in DMSO and cells exposed to a range of concentrations (1-100 μM) of the PCB 132 atropisomers for 4 or 24 h in cell medium. The final concentration of DMSO was $\leq$1.7% in N27 and $\leq$0.85% in PC12 cells. Vehicle controls of the relative %DMSO were used (i.e. $\leq$1.7%). After treatment with the PCB 132 atropisomers, the medium was removed. The cells were then incubated in MTT (Invitrogen) at a final concentration of 0.5 mg/ml in Hank’s Balanced Salt Solution (HBSS) with glucose (1 mg/mL) for 1 to 4 h. After incubation, the MTT-containing buffer was removed and DMSO added to the wells to solubilize the formazan product. The MTT reduction was measured at 570 nm and 650 nm with a Molecular Devices SpectraMax 190 plate reader. The values at 650 nm were subtracted from those at 570 nm and plotted as percent of control in order to show the formation of the formazan product alone.

**Detection of reactive oxygen species in N27 cells**

**DCFDA fluorescent probe:** Ninety-six well plates of N27 cells were washed with 1X DPBS. Then the cells were incubated in 25 μM DCFDA in glucose supplemented Hank’s Buffered Sodium Solution (HBSS) (1 mg/mL) for 45 min. After incubation, the cells were washed again with 1X DPBS and incubated in 100 μM dopamine for 1 h. Next the cells were washed with 1X DPBS and treated with the PCB 132 atropisomers at a range of concentrations (1-25 μM) or the positive control (200$\mu M$ H_2_O_2_). The fluorescence was then read with a Molecular Devices SpectraMax M5 plate reader over the course of 24 h with an excitation wavelength of 485 nm and an emission wavelength of 535 nm.

**DHE fluorescent probe:** Similar to the DCFDA experiment, N27 cells were rinsed with 1X DPBS and incubated in dihydroethidium (DHE) (25 μM) for 20 min. The cells were rinsed again and incubated with dopamine (100 μM) for 1 h. After incubation, the cells were rinsed and treated with the PCB 132 atropisomers or the positive control (100 μM) paraquat. The fluorescence was read with the following excitation/emission settings: 480/576 nm and 515/600 nm. The excitation/emission at 515/600 nm is intended to detect superoxide formation, while the excitation/emission at 480/576 nm detects general reactive oxygen species.

**Dopamine metabolite analysis in PC12 media**

We analyzed dopamine and dopamine metabolites via HPLC as previously described (Enayah *et al.*, 2018; Mexas *et al.*, 2011) utilizing an Agilent 1200 Series Capillary HPLC system coupled to a photodiode array detector set to absorbance at 202 and 280 nm. After seeding the cells in 6-well plates they were kept under the same growth conditions for 48h. Media was then removed to eliminate interactions with DA and subsequent metabolites, and cells placed in HEPES buffer (115 mM NaCl_2_, 5.4 mM KCl, 1.8 mM CaCl_2_, 0.8 mM MgSO_4_, 5.5 mM glucose, 1 mM NaH_2_PO_4_, and 15 mM HEPES, pH = 7.4). The cells were then treated with the PCB 132 atropisomers at 10 and 25 μM and incubated for 24 h. Aliquots of the extracellular buffer were taken at 0, 12, and 24 h by collecting 95 μL of HEPES buffer. Perchloric acid (5% v/v) was added to the aliquots to precipitate protein and samples were stored in -20˚C until analysis. On the day of HPLC analysis, the samples were centrifuged at 10,000g for 10 min, and 10 μL of supernatant injected into the HPLC to measure extracellular DA and metabolites. Separation was achieved with a Phenomenex Luna C18 column (1 x 150 mm, 100 Å) and an isocratic mobile phase of 0.1% TFA in HPLC H_2_O and 6% acetonitrile (ACN). Conversion of AUC to concentration was determined via calibration curves of standards.

**Table S1.** Purity determination of PCB 132 using GC-MS.^a^

| **PCB congener** | **Purity (peak area %)** |
| --- | --- |
| PCB 132 | 99.80 |
| PCB 84+92 | 0.01 |
| PCB 89 | 0.01 |
| PCB 136 | 0.01 |
| PCB 135+144 | 0.11 |
| PCB 174+181 | 0.05 |

^a^ PCB 132 (Accustandard, New Haven, CT, USA, Lot# GC950828R-AC) was analyzed by GC-MS in our laboratory as described previously (Holland *et al.*, 2017; Li *et al.*, 2018a; Uwimana *et al.*, 2018).

**Table S2.** Quantification of OH-PCB 132 metabolites (ng/mg microsomal protein) in incubations with different human liver microsome (HLM) preparations. Data are expressed as mean ± SD, n = 3.

| **Incubation time** | **HLM Preparation** | **3'-140 (ng/mg protein)** | **5'-132 (ng/mg protein)** | **4'-132 (ng/mg protein)** | **Conversion (%)^a^** |
| --- | --- | --- | --- | --- | --- |
| **10 min^b^** | **pHLM^c^** | 26 ± 10 | 28 ± 20 | 9 ± 4 | 0.03 |
|  | **H1** | 28 ± 1 | 7 ± 1 | 8 ± 1 | 0.02 |
|  | **H2**^d^ | 23 ± 1 | 17 ±1 | 10 ± 1 | 0.03 |
|  | **H3** | 40 ± 2 | 58 ± 4 | 15 ± 1 | 0.06 |
|  | **H4** | 35 ± 1 | 130 ± 10 | 15 ± 1 | 0.10 |
|  | **H5** | 40 ± 1 | 67 ± 2 | 17 ± 1 | 0.07 |
| **30 min^b^** | **pHLM^c^** | 75 ± 6 | 94 ± 9 | 23 ± 2 | 0.11 |
|  | **H1** | 83 ± 10 | 29 ± 4 | 22 ± 2 | 0.07 |
|  | **H2** | 70 ± 4 | 66 ± 5 | 29 ± 3 | 0.09 |
|  | **H3** | 130 ± 2 | 210 ± 6 | 43 ± 1 | 0.21 |
|  | **H4^d^** | 100 ± 1 | 420 ± 4 | 38 ± 1 | 0.31 |
|  | **H5** | 110 ± 5 | 250 ± 20 | 47 ± 4 | 0.23 |
| **120 min** | **pHLM^e,f^** | 15 ± 1 | 120 ± 10 | 10 ± 1 | 4.1 |
|  | **pHLM^f,g^** | 108 ± 14 | 300 ± 20 | 43 ± 4 | 1.2 |

Metabolites levels were quantified by GC-µECD using the internal standard method as described in the Materials and Methods section.

^a^ Percent of ΣOH-PCB of the total racemic PCB 132 added to each incubation.

^b^ Incubation conditions: 50 μM PCB 132; 10 or 30 minute incubation at 37 ºC; 0.1 mg/mL microsomal protein; and 1 mM NADPH. These experimental conditions were optimized with regard to incubation time, and the metabolite formation was linear up to 30 min.

^c^ Lot number 1410013.

^d^ n = 2

^e^ Incubation conditions: 5 μM PCB 132; 120 minute incubation at 37 ºC; 0.5 mg/mL microsomal protein; and 0.5 mM NADPH.

^f^ Lot number 0910398.

^g^ Incubation conditions: 50 μM PCB 132; 120 minute incubation at 37 ºC; 0.5 mg/mL microsomal protein; and 0.5 mM NADPH.

**Table S3.** Rate of PCB 132 metabolite formation (ng OH-PCB/nmol P450/min) by different human liver microsome (HLM) preparations. Data are expressed as mean ± SD, n = 3.^a,b,c^

| **HLM Preparation** | **3'-140**  (ng/nmol P450/min) | **5'-132**  (ng/nmol P450/min) | **4'-132**  (ng/nmol P450/min) |
| --- | --- | --- | --- |
| **pHLM^d^** | 4.5 ± 2.2 | 4.8 ± 3.0 | 1.5 ± 0.7 |
| **H1** | 4.1 ± 0.2 | 1.0 ± 0.1 | 1.2 ± 0.1 |
| **H2**^e^ | 5.3 ± 0.1 | 3.8 ± 0.2 | 2.3 ± 0.1 |
| **H3** | 8.4 ± 0.5 | 12.1 ± 0.9 | 3.0 ± 0.3 |
| **H4** | 4.4 ± 0.1 | 16.8 ± 1.3 | 1.9 ± 0.1 |
| **H5** | 9.0 ± 0.3 | 19.1 ± 0.5 | 3.9 ± 0.3 |

Metabolites levels were quantified by GC-µECD using the internal standard method as described in the Materials and Methods section.

^a^ Microsomal metabolism studies were performed using the following incubation conditions: 50 μM PCB 132; 10 minute incubation at 37 ºC; 0.1 mg/mL microsomal protein; and 1 mM NADPH.

^b^ Rates of metabolism were calculated as described previously (Uwimana *et al.*, 2018).

^d^ Lot number 1410013.

^e^ n = 2.

**Table S4.** Rate of PCB 132 metabolite formation (ng OH-PCB/mg microsomal protein/min) by different human liver microsome (HLM) preparations. Data are expressed as mean ± SD, n = 3.^a,b^

| **HLM Preparation** | **3'-140**  (ng/mg protein/min) | **5'-132**  (ng/mg protein/min) | **4'-132**  (ng/mg protein/min) |
| --- | --- | --- | --- |
| **pHLM^c^** | 2.6 ± 1.3 | 2.8 ± 1.7 | 0.9 ± 0.4 |
| **H1** | 2.8 ± 0.1 | 0.7 ± 0.1 | 0.8 ± 0.1 |
| **H2**^d^ | 2.3 ± 0.1 | 1.7 ± 0.1 | 1.1 ± 0.1 |
| **H3** | 4.1 ± 0.2 | 5.8 ± 0.4 | 1.5 ± 01 |
| **H4** | 3.5 ± 0.1 | 13 ± 1 | 1.5 ± 0.1 |
| **H5** | 4.0 ± 0.1 | 6.7 ± 0.2 | 1.7 ± 0.1 |

Metabolites levels were quantified by GC-µECD using the internal standard method as described in the Materials and Methods section.

^a^ Microsomal metabolism studies were performed using the following incubation conditions: 50 μM PCB 132; 10 minute incubation at 37 ºC; 0.1 mg/mL microsomal protein; and 1 mM NADPH.

^b^ Rates of metabolism were calculated as described previously (Uwimana *et al.*, 2018).

^c^ Lot number 1410013.

^d^ n = 2

**Table S5.** Summary of relative free energies for epoxide ring opening. A blank cell indicates that the species was not computed.^a,b^

| **3,4-arene oxides** | **1,2,4-Trichlorobenzene^c^** | | **2,3,6-Trichloro-biphenyl** | | **PCB 91** | | **PCB 95** | | **PCB 132** | | **PCB 136** | |
| --- | --- | --- | --- | --- | --- | --- | --- | --- | --- | --- | --- | --- |
| **ΔG (kcal/mol)** |  |  |  |  |  |  |  |  |  |  |  |  |
| **Toward** | C-Cl | C-H | C-Cl | C-H | C-Cl | C-H | C-Cl | C-H | C-Cl | C-H | C-Cl | C-H |
| *Reactant* | 0 | 0 | 0 |  | 0 |  | 0 |  | 0 | 0 | 0 | 0 |
| *TS1* | 22 | 42 | 23 |  | 23 |  | 23 |  | 23 | 42 | 23 | 44 |
| *Intermediate* | -21 | -27 |  |  | -21 |  | -20 |  | -21 | 1 | -20 | 1 |
| *TS2* | -16 | -20 |  |  | -17 |  | -16 |  | -16 | - | -16 | - |
| *Phenol product* | -48 | -36 |  |  | -48 |  | -48 |  | -47 | - | -50 | - |
| **4,5-arene oxides** | *Energies relative to the 3,4-arene oxide* | | | | | | | | | | | |
| **ΔG (kcal/mol)** |  |  |  |  |  |  |  |  | **PCB 132** | | **PCB 136** | |
| **Toward** |  |  |  |  |  |  |  |  | C-H_4_ | C-H_5_ | C-H_4_ | C-H_5_ |
| *Reactant* |  |  |  |  |  |  |  |  | 1 | -1 | 1 | 0 |
| *TS1* |  |  |  |  |  |  |  |  | 38 | 42 | 39 | 42 |
| *Intermediate* |  |  |  |  |  |  |  |  | -26 | -27 | -25 | -26 |
| *TS2* |  |  |  |  |  |  |  |  | -20 | -20 | -20 | -21 |
| *Phenol product* |  |  |  |  |  |  |  |  | -49 | -49 | -48 | -48 |
| **5,6-arene oxides** | *Energies relative to the 3,4-arene oxide* | | | | | | | | | | | |
| **ΔG (kcal/mol)** |  |  |  |  |  |  |  |  | **PCB 132** | | **PCB 136** | |
| **Toward** |  |  |  |  |  |  |  |  | C-Cl | C-H | C-Cl | C-H |
| *Reactant* |  |  |  |  |  |  |  |  | 0 | -1 | 0 | 0 |
| *TS1* |  |  |  |  |  |  |  |  | 26 | 39 | 27 | 40 |
| *Intermediate* |  |  |  |  |  |  |  |  | -22 | 0 | -22 | -1 |
| *TS2* |  |  |  |  |  |  |  |  | -18 | - | -17 | - |
| *Phenol product* |  |  |  |  |  |  |  |  | -48 | - | -48 | - |

^a^ Structures and energies computed at the M11/def2-SVP level of theory with the SMD aqueous continuum solvation model. All calculations were performed using Gaussian 16, Rev. A.01 (Frisch *et al.*, 2016).

^b^ The differences in ΔG seen across each series are insignificant and within the error limits of the computational method applied.

^c^ The 3,4-arene oxide to 4,5-arene oxide rearrangement discussed for PCB 132 and PCB 136 described in the manuscript does not exist for 1,2,4-trichlorobenzene. Instead, a 1,2-hydride shift is predicted. However, the energetics was not affected by this variation in the mechanism.

**Table S6:** Enantiomeric fractions (EFs) of PCB 132, 3'-140 and 5'-132 in incubations with human liver microsomes reveal the atropselective formation of the second eluting atropisomer of 5'-132 (E_2_-5'-132) and nearly enantiospecific formation the first eluting atropisomer of 3'-140 (E_1_-3'-140). A depletion of the second eluting atropisomer of PCB 132 [*i.e.*, (+)-PCB 132] was observed. If not stated otherwise, data are expressed as mean ± SD, n = 3.^a^

| **HLM Preparation** | **EF** | | |
| --- | --- | --- | --- |
|  | **PCB 132**^b^ | **3'-140**^c^ | **5'-132**^d^ |
| **Pooled**^e,f^ | 0.49 ± 0.01 | 0.84 ± 0.09 | 0.19 ± 0.01 |
| **H1**^e^ | 0.48 ± 0.01 | 0.94 ± 0.01 | 0.16 ± 0.01 |
| **H2**^e^ | 0.49 ± 0.01 | 0.93 ± 0.01^g^ | 0.14 ± 0.01 |
| **H3**^e^ | 0.48 ± 0.01 | 0.95 ± 0.01 | 0.17 ± 0.01 |
| **H4**^e,h^ | 0.49 ± 0.01 | 0.94 ± 0.01 | 0.12 ± 0.01 |
| **H5**^e^ | Nd | Nd | Nd |
| **pHLM**^h,i^ | 0.39 ± 0.01 | 0.96 ± 0.01 | 0.19 ± 0.02 |
| **pHLM**^h,j^ | 0.47 ± 0.01 | 0.95 ± 0.04 | 0.14 ± 0.01 |

^a^ To allow a comparison with previously published data (Uwimana *et al.*, 2016, 2018; Uwimana *et al.*, 2017; Wu *et al.*, 2016; Wu *et al.*, 2011), EF values were calculated by the drop valley method (Asher *et al.*, 2009) using the following equation: EF = Area E_1_/(Area E_1_ + Area E_2_), where Area E_1_ and Area E_2_ denote the peak area of the first and second eluting atropisomer. Please see the Materials and Methods for additional details regarding the atropselective gas chromatographic analyses.

^b^ Resolution of PCB 132 atropisomers = 1.05.

^c^ Resolution of 3'-140 atropisomers = 1.18.

^d^ Resolution of 5'-132 atropisomers = 2.02.

^e^ Incubation conditions: 50 µM PCB 132; 30 minute incubation at 37 ºC; 0.1 mg/mL microsomal protein; and 1 mM NADPH.

^f^ Lot number 1410013.

^g^ n = 2.

^h^ Lot number 01320398.

^i^ Incubation conditions: 5 µM PCB 132; 120 minute incubation at 37 ºC; 0.5 mg/mL microsomal protein; and 0.5 mM NADPH.

^j^ Incubation conditions: 50 µM PCB 132; 120 minute incubation at 37 ºC; 0.5 mg/mL microsomal protein; and 0.5 mM NADPH.

nd = not detected

**Table S7.** PCB 132 metabolite levels in incubations of individual atropisomers and racemic PCB 132 with pHLMs (see **Fig. 8**).^a^

| **PCB 132 metabolites** | **(+)-PCB 132** | | **Racemic PCB 132**^b^ | | **(-)-PCB 132** | |
| --- | --- | --- | --- | --- | --- | --- |
|  | Concentration  (ng/mg protein) | EF | Concentration  (ng/mg protein) | EF | Concentration  (ng/mg protein) | EF |
| **3'-140** | 3.2 ± 0.2 | nd | 90 ± 1 | 0.89 ± 0.02 | 187 ± 4 | 0.99 ± 0.01 |
| **5'-132** | 58 ± 3 | 0.93 ± 0.01 | 100 ± 1 | 0.20 ± 0.3 | 130 ± 20 | 0.005 ± 0.002 |
| **4'-132** | 40 ± 2 | nr | 27 ± 1 | nr | 17 ± 1 | nr |
| **Σ-OH-PCBs** | 101 ± 6 |  | 217 ± 1 |  | 332 ± 12 |  |

^a^ Microsomal metabolism studies were performed using the following incubation conditions: 50 μM PCB 132; 30 minute incubation at 37 ºC; 0.1 mg/mL microsomal protein (Lot number 1410013); and 1 mM NADPH.

^b^ n = 2
nd: not detected

nr: not resolved

**Table S8.** Summary of chiral PCB 132 signatures in human samples.

| **Tissue** | **Country of Origin** | **Number of samples** | **EF values** | **Reference** |
| --- | --- | --- | --- | --- |
| Milk | Germany | 10 | 0.29–0.47^a^ | (Glausch *et al.*, 1995) |
| Milk | Spain | 11 | 0.35–0.48 | (Bordajandi *et al.*, 2008) |
| Milk | Swiss | 4 | 0.18–0.33**^b^** | (Bucheli *et al.*, 2006) |
| Brain | Belgium | 1 | 0.52^a^ | (Chu *et al.*, 2003) |
| Kidney | Belgium | 3 | 0.49–0.51^a^ | (Chu *et al.*, 2003) |
| Liver | Belgium | 11 | 0.32 to 0.49^a^ | (Chu *et al.*, 2003) |
| Muscle | Belgium | 3 | 0. 47–0.50^a^ | (Chu *et al.*, 2003) |
| Hair | China | 97 | 0.48±0.03 | (Zheng *et al.*, 2013) |

^a^ ER values were converted to EF values using the formula EF = 1/(1 + 1/ER) (Harner *et al.*, 2000).

^b^ Enantiomeric fractions (EF) values were recalculated as EF = E_1_/(E_1_+E_2_) to allow a direct comparison of EF values.

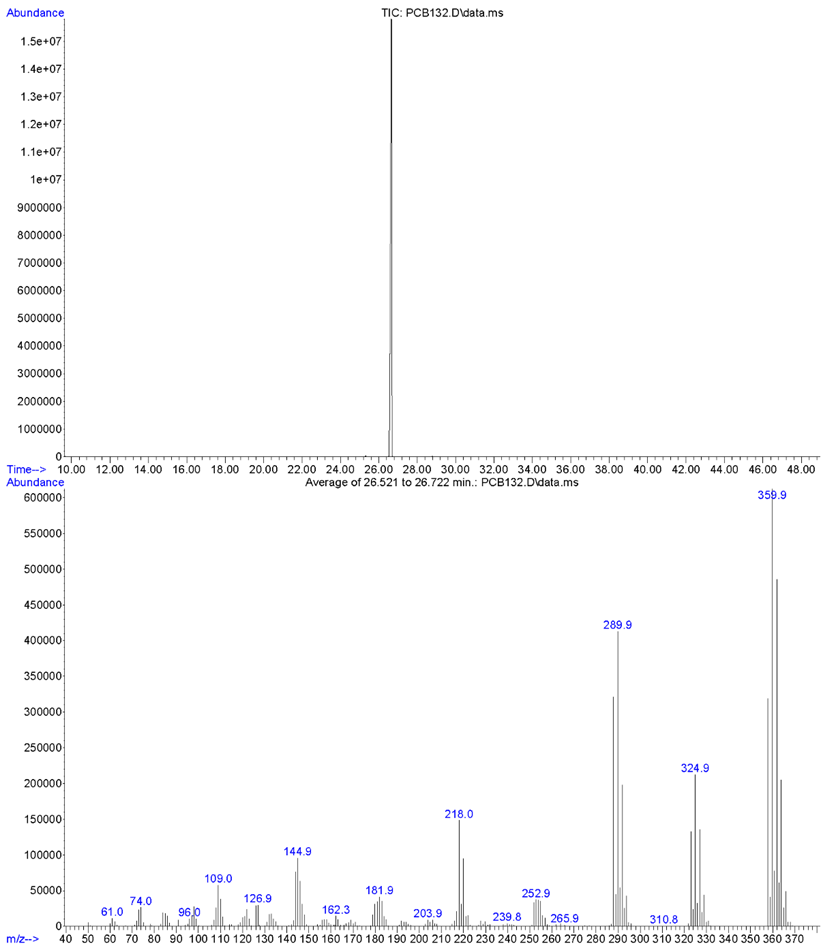

**Fig. S1.** GC-MS chromatogram and mass spectrum of the test compound, PCB 132, analyzed in the TIC mode. See description above for experimental details.

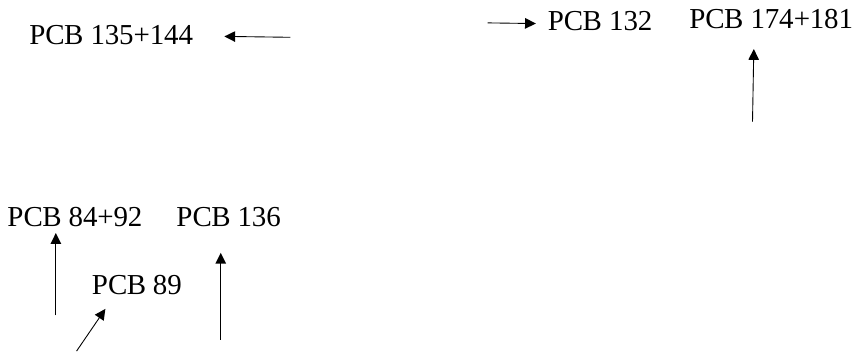

**Fig. S2.** Enlarged chromatogram showing minor impurities (retention times 23.403, 23.483, 24.662, 25.320, and 28.793 min) present in PCB 132 by GC-MS (TIC) (**Table S1**). The impurities were identified as specific PCB congeners using the congener specific analysis method mentioned above (see Table S1).

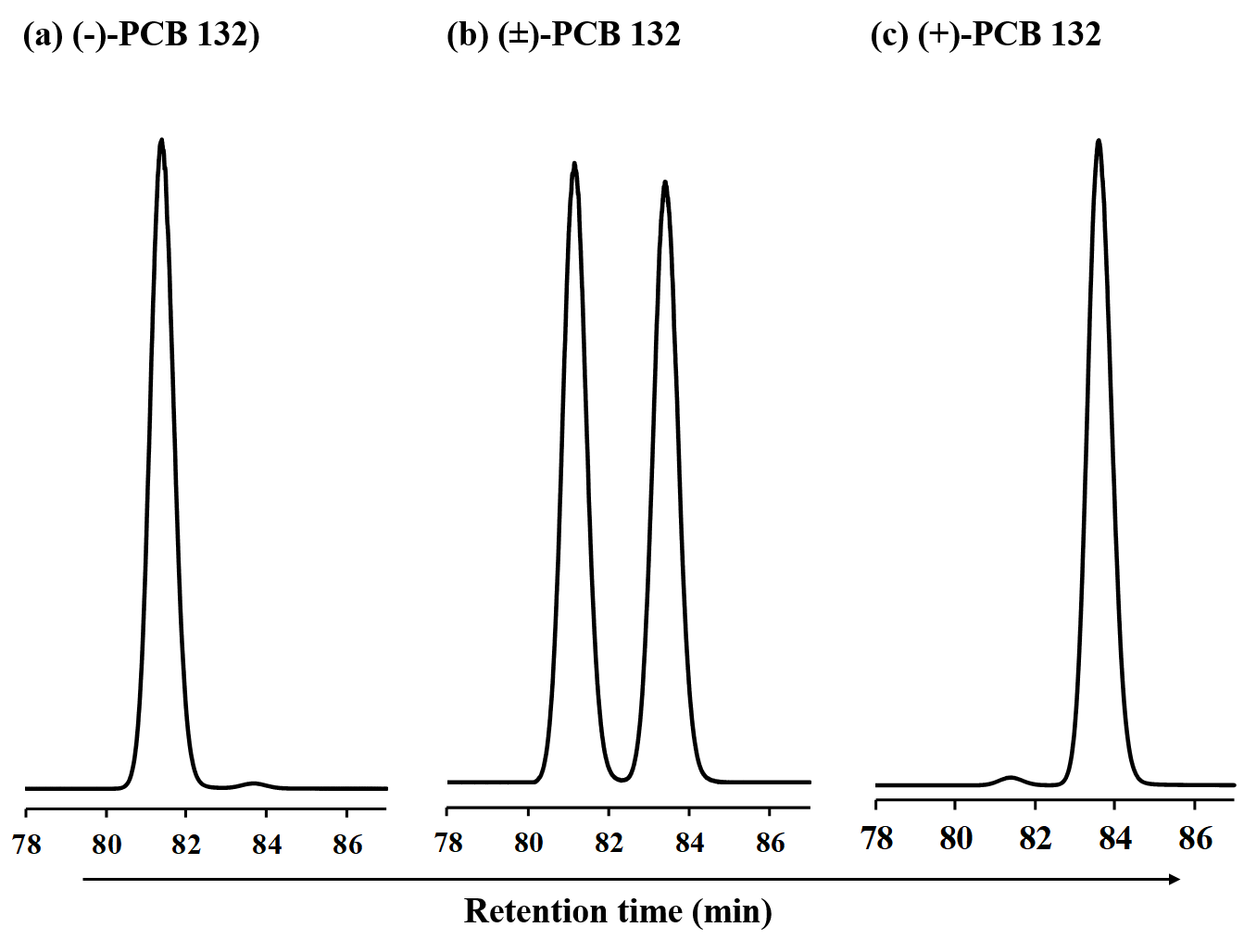

**Fig. S3.** Enantioselective separation of (a) (-)-PCB 132, (b) racemic PCB 132 and (c) (+)-PCB 132 on a GC-µECD equipped with a ChiralsilDex-CD column. (-)-PCB 132 (EF = 0.97) and (+)-PCB 132 (EF = 0.05) correspond to E_1_-PCB 132 and E_2_-PCB 132, respectively (Haglund *et al.*, 1996). For details regarding the separation of the pure PCB 132 atropisomers, see section “HPLC separation of PCB 132 atropisomers” above. Gas chromatographic separations were performed at 160 °C.

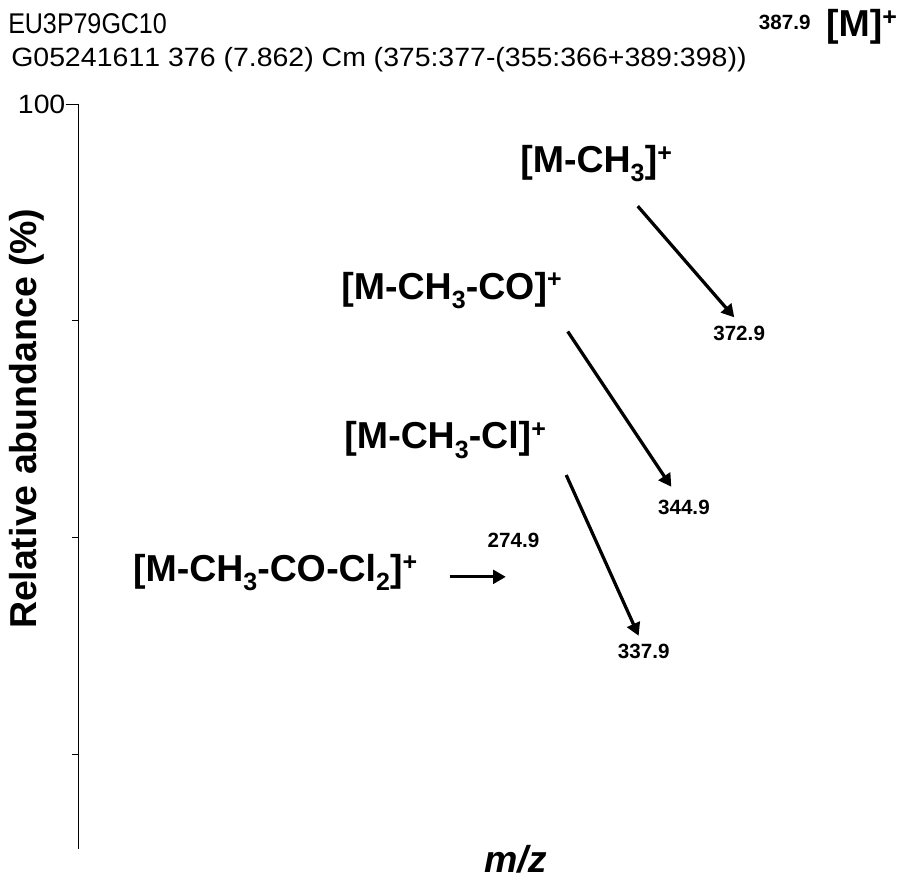
 **Fig. S4.** Mass spectrum of the authentic standard of methylated 3'-140 (RT 7.87 min; RRT 1.809). The accurate mass of the monoisotopic ion (*m/z* 387.8535 compared to theoretical *m/z* 387.8550 determined for C_13_H_6_O_1_^35^Cl_6_), the isotope pattern of the molecular ion (abundance ratio: 1:1.9:1.6:0.6:0.1 compared to predicted abundance ratio 1:1.9:1.6:0.7:0.2) and the fragmentation pattern are consistent with a monohydroxylated hexachlorobiphenyl (as the corresponding methylated derivative). The mass spectrum was recorded without the lock standard to improve the sensitivity, for additional information see manuscript. For the corresponding gas chromatogram of the authentic standards, see **Fig. 2a**.

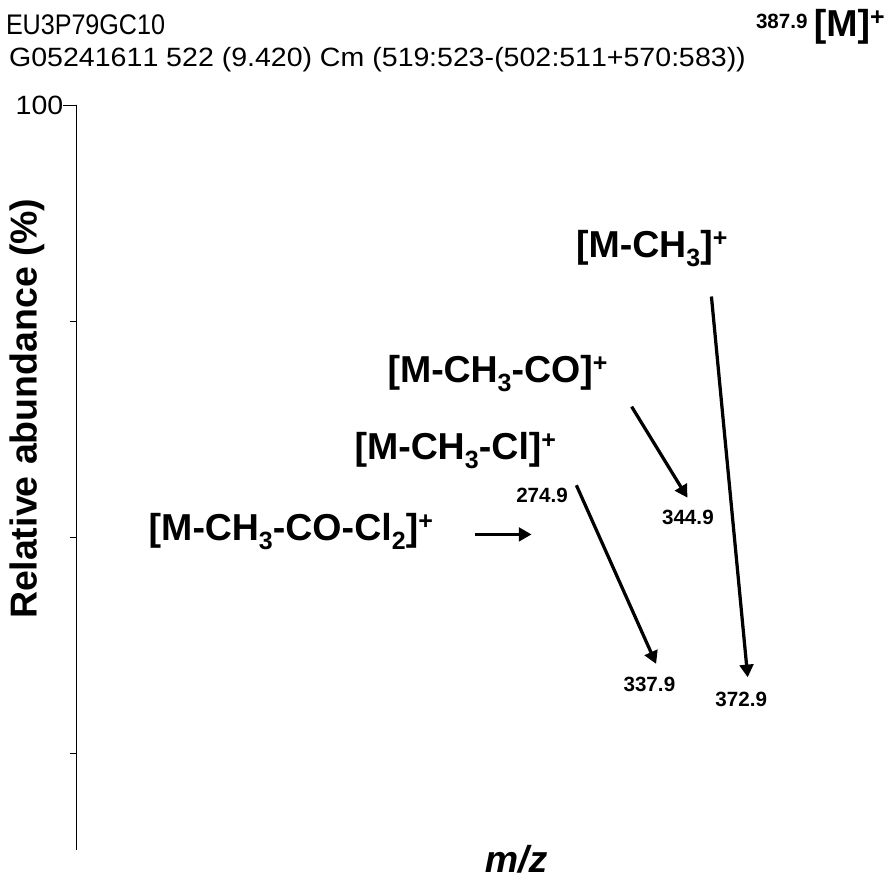
**Fig. S5.** Mass spectrum of the authentic standard of methylated 5'-132 (RT 9.44 min; RRT 2.170). The accurate mass of the monoisotopic ion (*m/z* 387.8557 compared to theoretical *m/z* 387.8550 determined for C_13_H_6_O_1_^35^Cl_6_), the isotope pattern of the molecular ion (abundance ratio: 1:1.9:1.5:0.7:0.2 compared to predicted abundance ratio 1:1.9:1.6:0.7:0.2) and the fragmentation pattern are consistent with a monohydroxylated hexachlorobiphenyl (as the corresponding methylated derivative). The mass spectrum was recorded without the lock standard to improve the sensitivity, for additional information see manuscript. For the corresponding gas chromatogram of the authentic standards, see **Fig. 2a**.

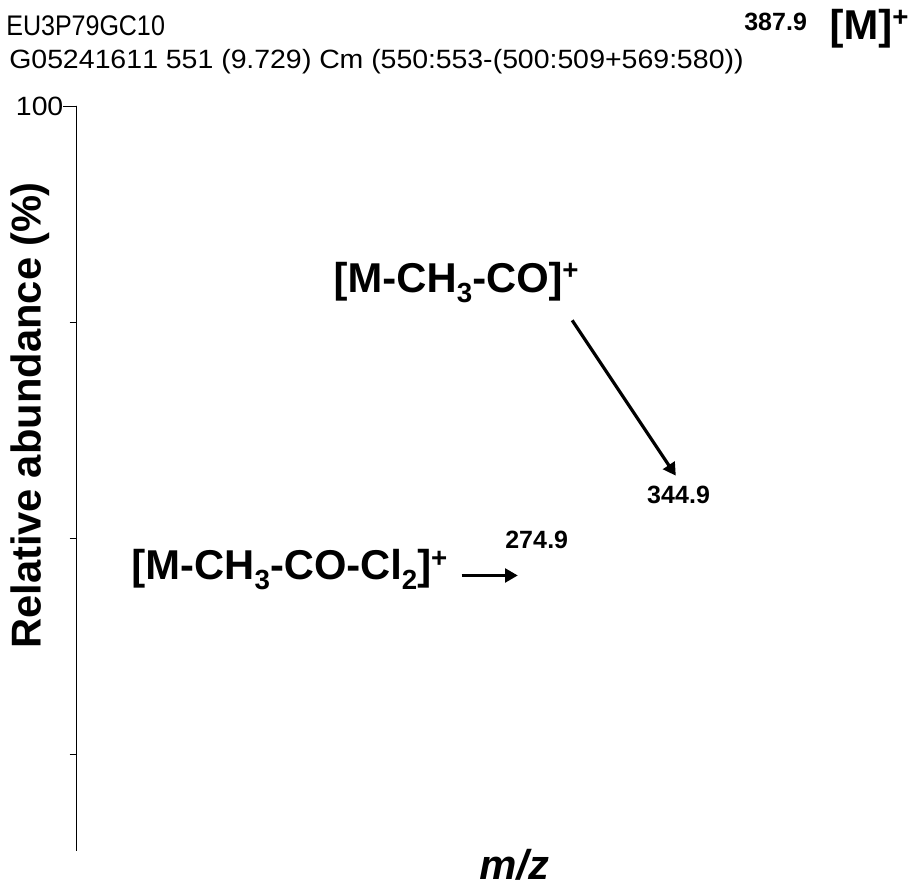
**Fig. S6.** Mass spectrum of the authentic standard of methylated 4'-132 (RT 9.76 min; RRT 2.244). The accurate mass of the monoisotopic ion (*m/z* 387.8522 compared to theoretical *m/z* 387.8550 determined for C_13_H_6_O_1_^35^Cl_6_), the isotope pattern of the molecular ion (abundance ratio: 1:2:1.6:0.7:0.2 compared to predicted abundance ratio 1:1.9:1.6:0.7:0.2) and the fragmentation pattern are consistent with a monohydroxylated hexachlorobiphenyl (as the corresponding methylated derivative). The mass spectrum was recorded without the lock standard to improve the sensitivity, for additional information see manuscript. For the corresponding gas chromatogram of the authentic standards, see **Fig. 2a**.

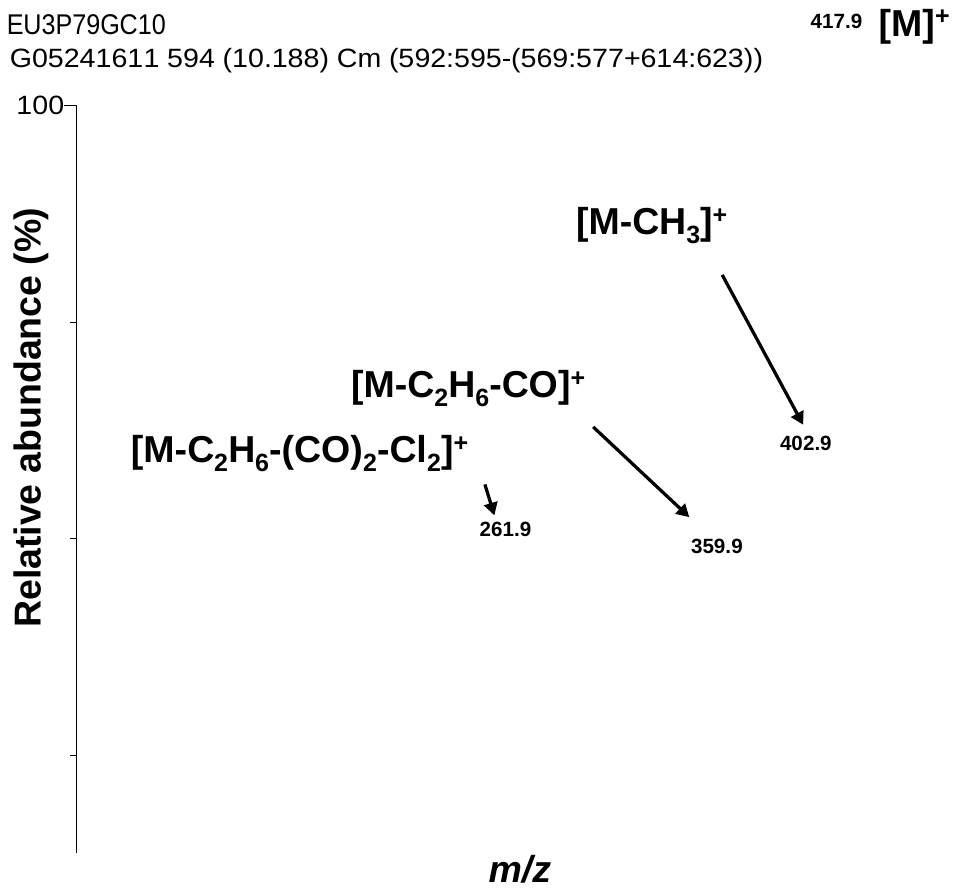
**Fig. S7.** Mass spectrum of the authentic standard of methylated 4',5'-132 (RT 10.19 min; RRT 2.343). The accurate mass of the monoisotopic ion (*m/z* 417.8635 compared to theoretical *m/z* 417.8655 determined for C_14_H_8_O_2_^35^Cl_6_), the isotope pattern of the molecular ion (abundance ratio: 1:2:1.6:0.7:0.2 compared to predicted abundance ratio 1:1.9:1.6:0.7:0.2) and the fragmentation pattern are consistent with a dihydroxylated hexachlorobiphenyl (as the corresponding methylated derivative). The mass spectrum was recorded without the lock standard to improve the sensitivity, for additional information see manuscript. For the corresponding gas chromatogram of the authentic standards, see **Fig. 2a**.

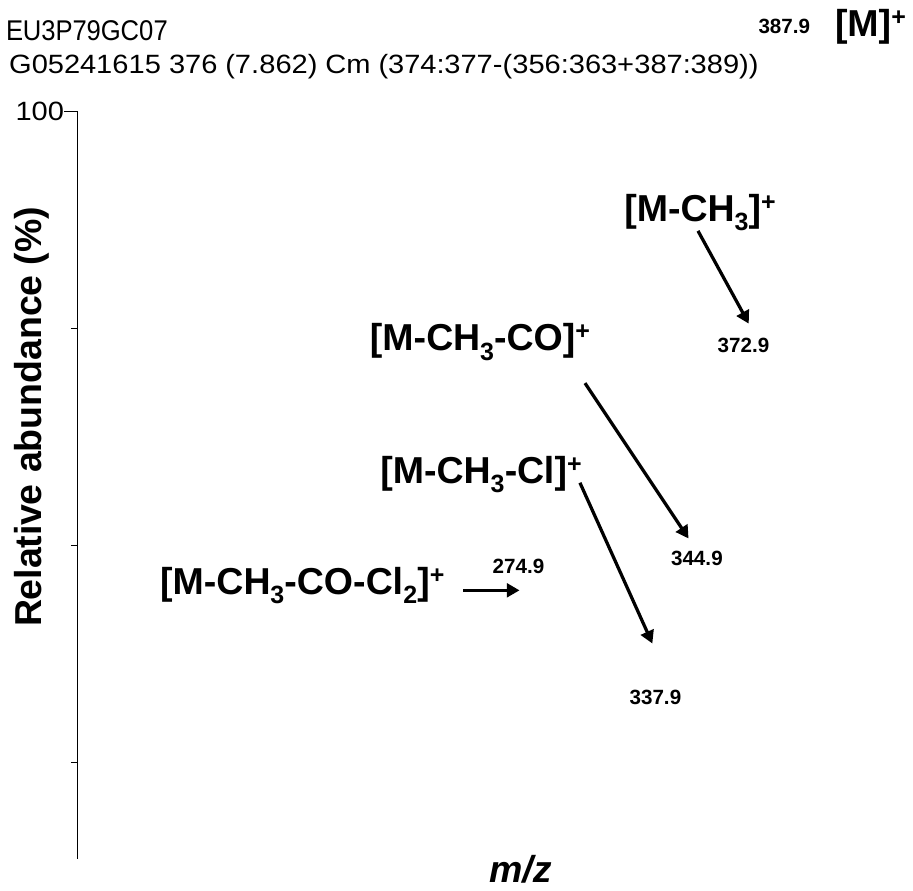
**Fig. S8.** Mass spectrum of 3'-140 (RT 7.86 min; RRT 1.803) formed in a representative incubation with pooled human liver microsomes (analyzed as methylated derivative). The accurate mass of the monoisotopic ion (*m/z* 387.8569 compared to theoretical *m/z* 387.8550 determined for C_13_H_6_O_1_^35^Cl_6_), the isotope pattern of the molecular ion (abundance ratio: 1:2:1.6:0.7:0.2 compared to predicted abundance ratio 1:1.9:1.6:0.7:0.2) and the fragmentation pattern are consistent with a monohydroxylated hexachlorobiphenyl (as the corresponding methylated derivative). The mass spectrum was recorded without the lock standard to improve the sensitivity, for additional information see manuscript. For the corresponding gas chromatogram, see **Fig. 2b**.

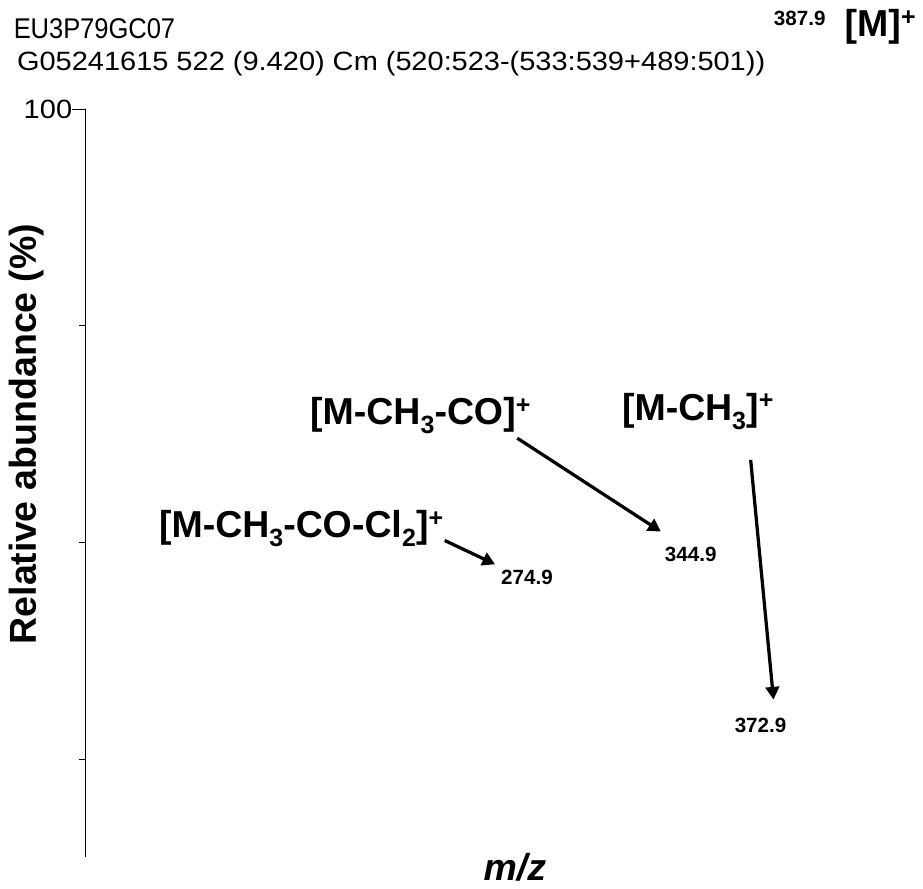
**Fig. S9.** Mass spectrum of 5'-132 (RT 9.42 min; RRT 2.161) formed in a representative incubation with pooled human liver microsomes (analyzed as methylated derivative). The accurate mass of the monoisotopic ion (*m/z* 387.8575 compared to theoretical *m/z* 387.8550 determined for C_13_H_6_O_1_^35^Cl_6_), the isotope pattern of the molecular ion (abundance ratio: 1:1.9:1.5:0.6:0.2 compared to predicted abundance ratio 1:1.9:1.6:0.7:0.2) and the fragmentation pattern are consistent with a monohydroxylated hexachlorobiphenyl (as the corresponding methylated derivative). The mass spectrum was recorded without the lock standard to improve the sensitivity, for additional information see manuscript. For the corresponding gas chromatogram, see **Fig. 2b**.

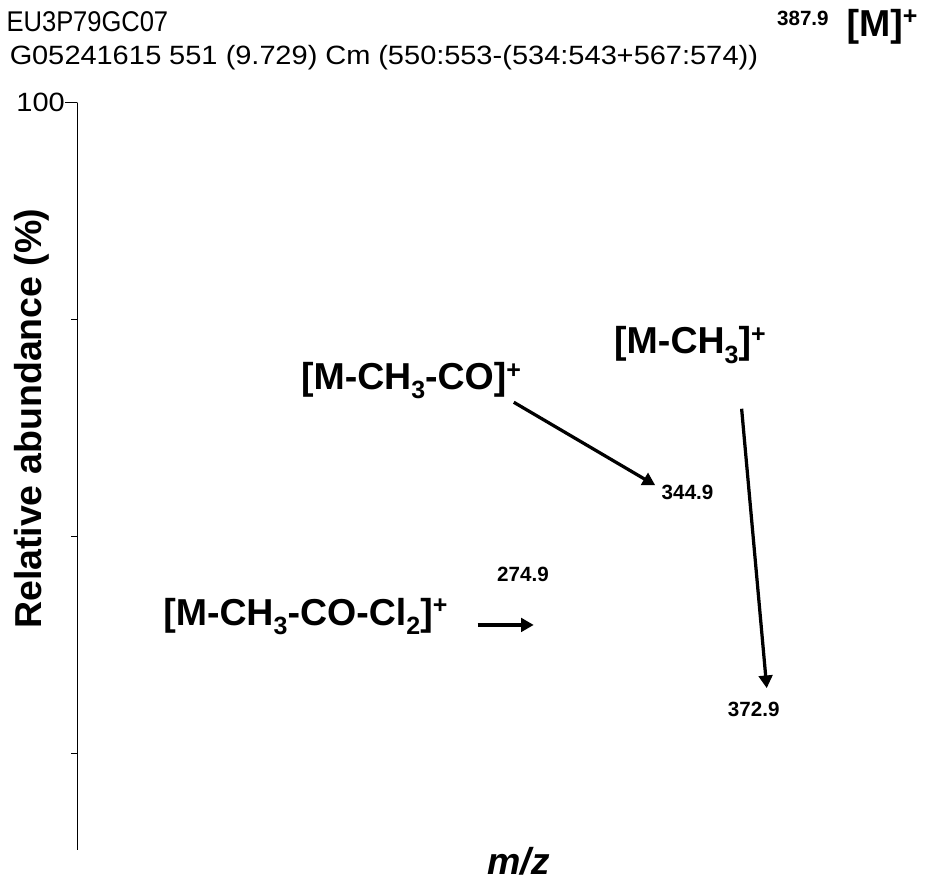
**Fig. S10.** Mass spectrum of 4'-132 (RT 9.74 min; RRT 2.234) formed in a representative incubation with pooled human liver microsomes (analyzed as methylated derivative). The accurate mass of the monoisotopic ion (*m/z* 387.8578 compared to theoretical *m/z* 387.8550 determined for C_13_H_6_O_1_^35^Cl_6_), the isotope pattern of the molecular ion (abundance ratio: 1:1.8:1.5:0.6:0.1 compared to predicted abundance ratio 1:1.9:1.6:0.7:0.2) and the fragmentation pattern are consistent with a monohydroxylated hexachlorobiphenyl (as the corresponding methylated derivative). The mass spectrum was recorded without the lock standard to improve the sensitivity, for additional information see manuscript. For the corresponding gas chromatogram, see **Fig. 2b**.

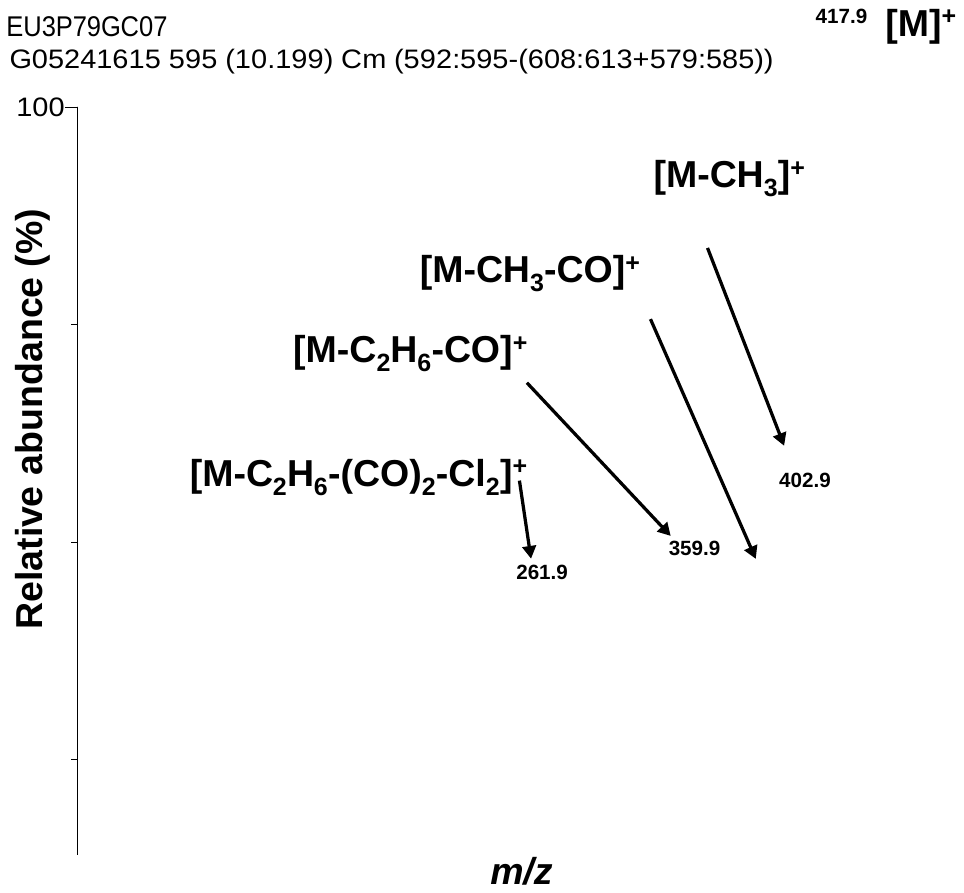
**Fig. S11.** Mass spectrum of 4',5'-132 (RT 10.19 min; RRT 2.337) formed in a representative incubation with pooled human liver microsomes (analyzed as methylated derivative). The accurate mass of the monoisotopic ion (*m/z* 417.8661 compared to theoretical *m/z* 417.8655 determined for C_14_H_8_O_2_^35^Cl_6_), the isotope pattern of the molecular ion (abundance ratio: 1:1.9:1.6:0.7:0.1 compared to predicted abundance ratio 1:1.9:1.6:0.7:0.2) and the fragmentation pattern are consistent with a dihydroxylated hexachlorobiphenyl (as the corresponding methylated derivative). The peak at *m/z* 257.3 corresponds to an impurity present in the background. The mass spectrum was recorded without the lock standard to improve the sensitivity, for additional information see manuscript. For the corresponding gas chromatogram, see **Fig. 2b**.

**Favored Opening Toward C-Cl**

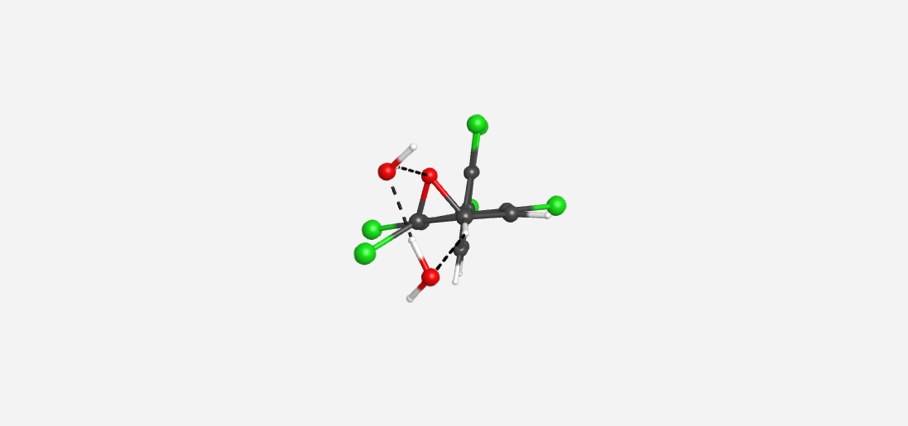

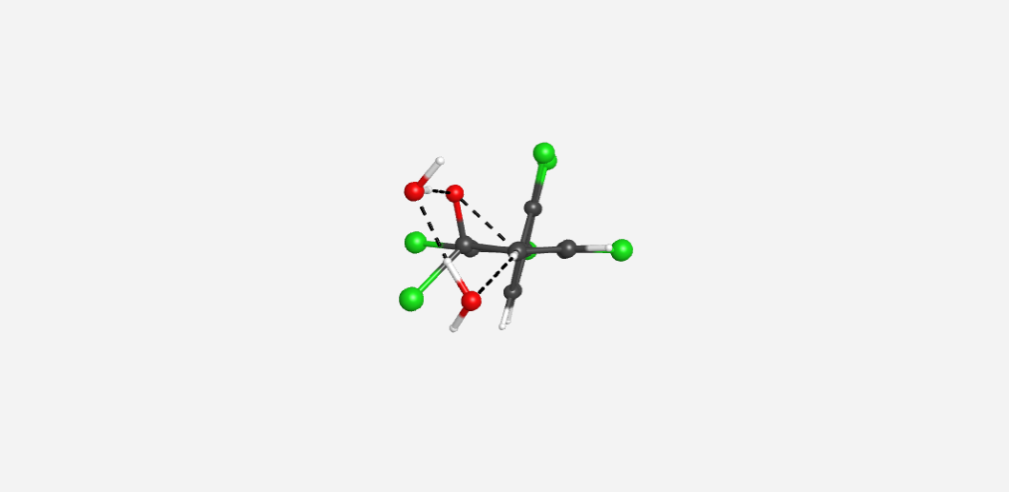

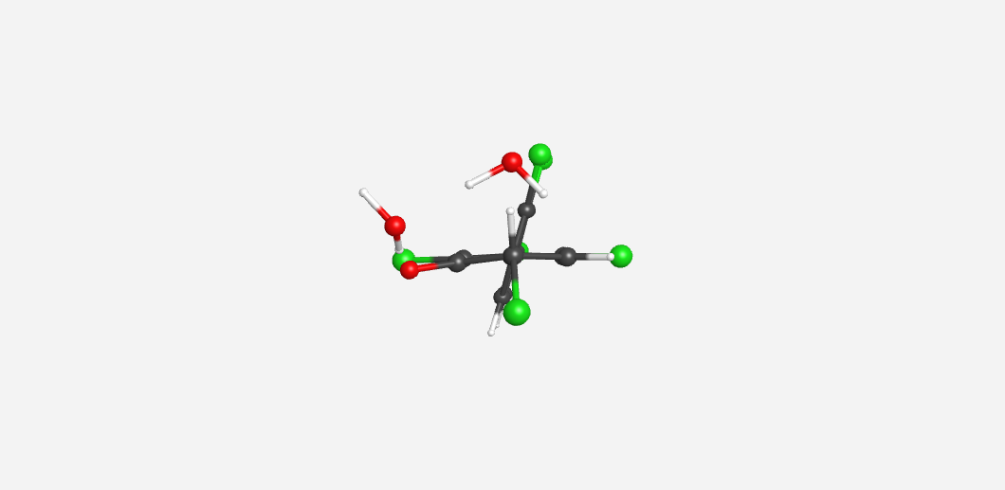

**PCB 132 3,4-arene oxide	Opening TS Toward C-Cl	Shifted Intermediate**

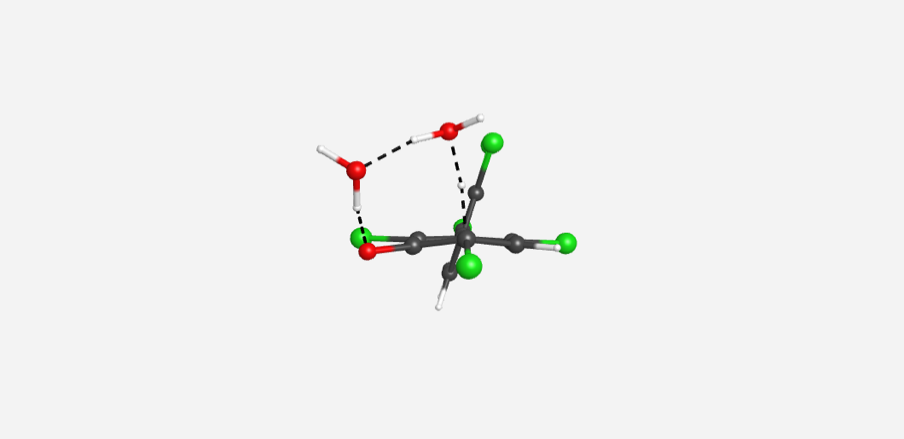

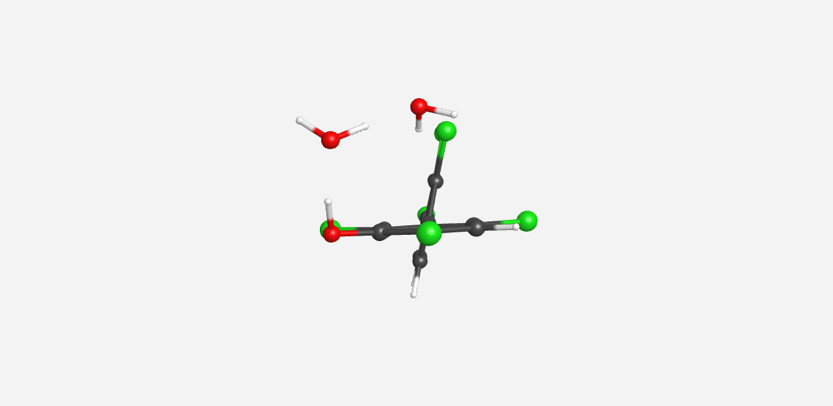

**TS for Rearomatization	3**'**-140**

**Disfavored Opening Toward C-H**

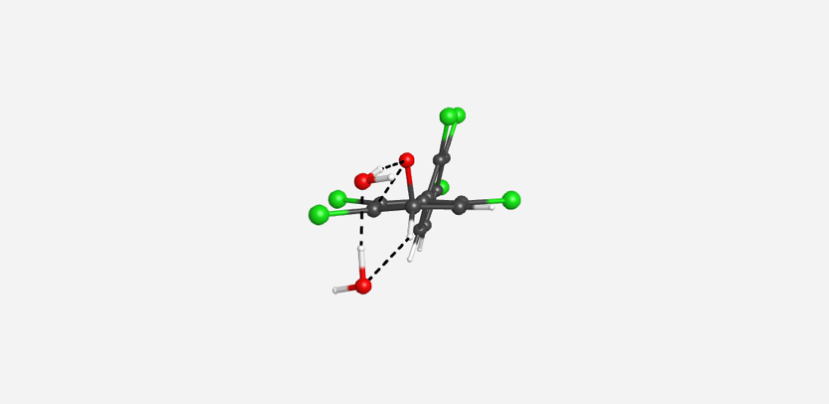

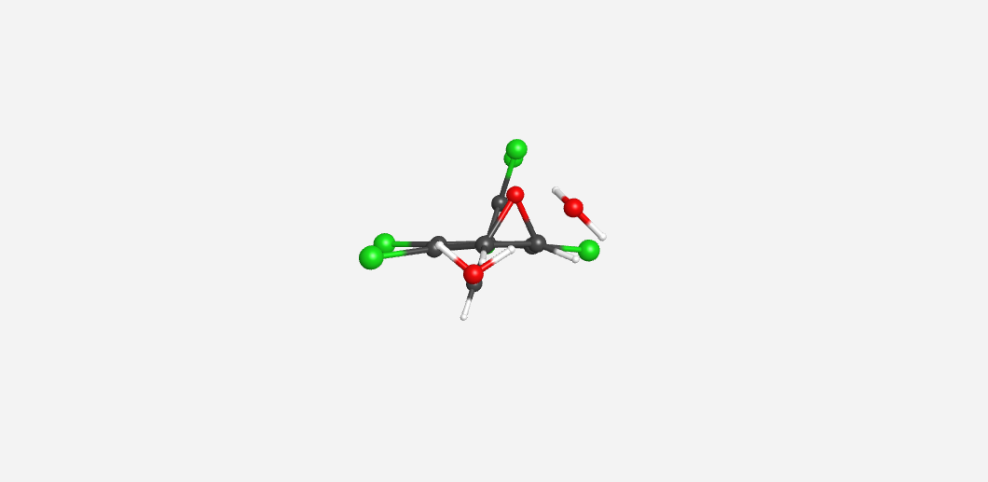

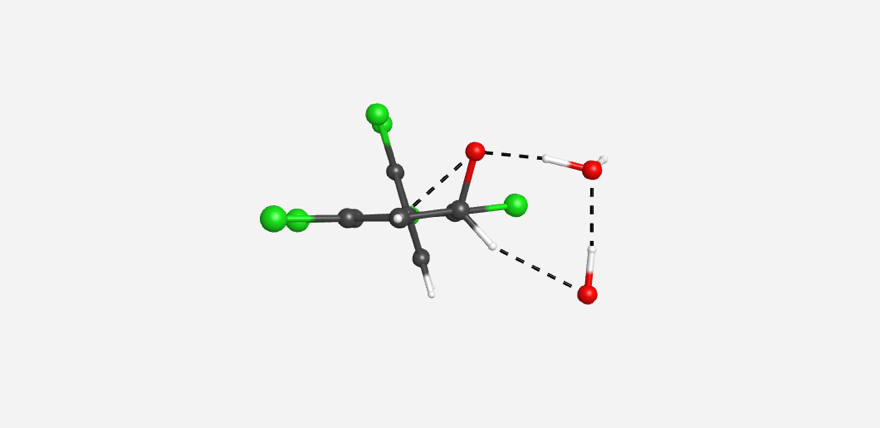

**Opening TS Toward C-H	PCB 132 4**'**,5**'**-arene oxide	TS for Opening**

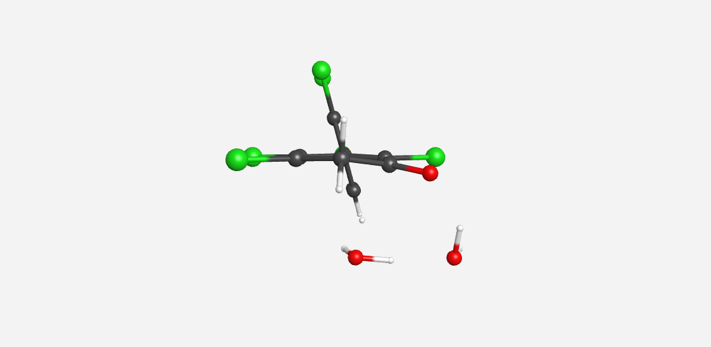

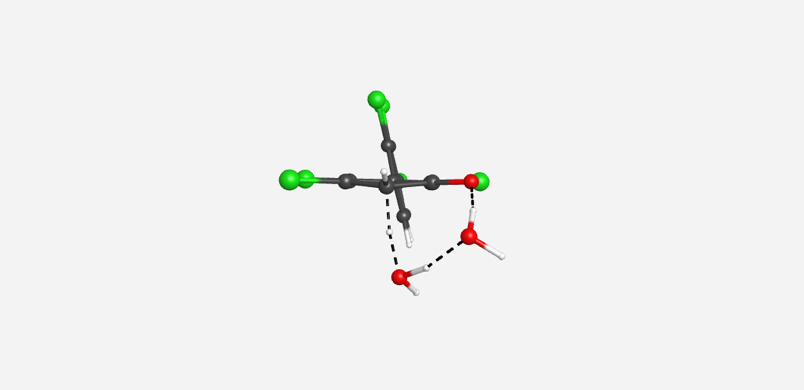

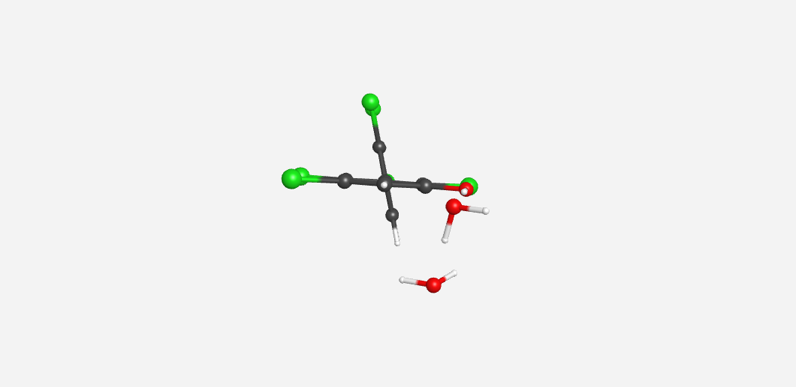

**Shifted Intermediate	TS for Rearomatization	5**'**-132**

**Fig. S12.** Ring-opening pathways determined for PCB 132 at the M11/def2-SVP + SMD level of theory. These pathways are general for all species considered herein.

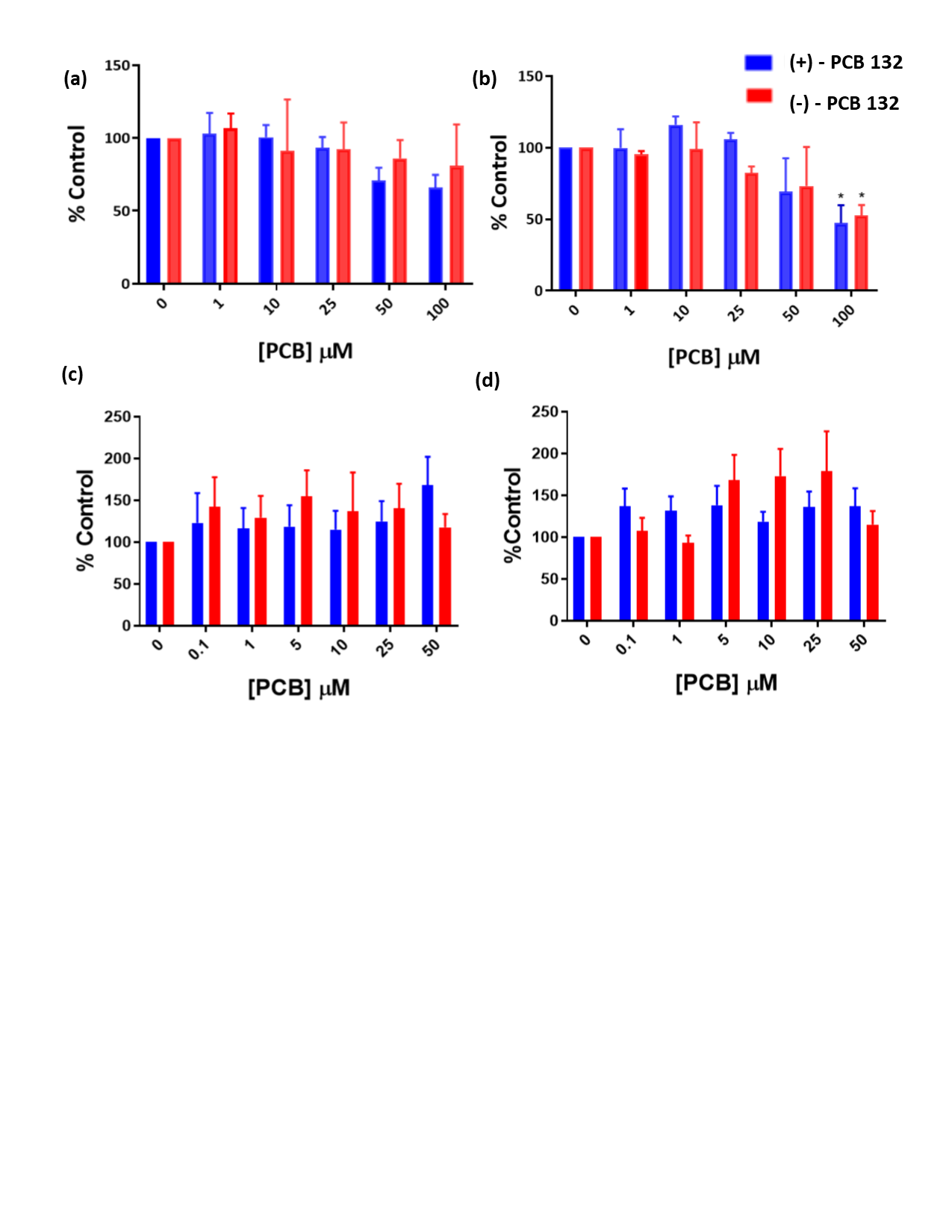

Fig. S13. Exposure to PCB 132 atropisomers has little effect on the viability of N27 and PC12 cell, as determined with the MTT assay. (a) N27 cell viability after 4 h exposure. (b) N27 cell viability after 24 h exposure. Exposure to 100 μM PCB 132 showed significant toxicity resulting in approximately 50% cell death. (c) PC12 cell viability after 4 h exposure. (d) PC12 cell viability after 24 h. Data are presented as mean ± standard error, n = 3. Significance determined by 2-way ANOVA, p < 0.05.

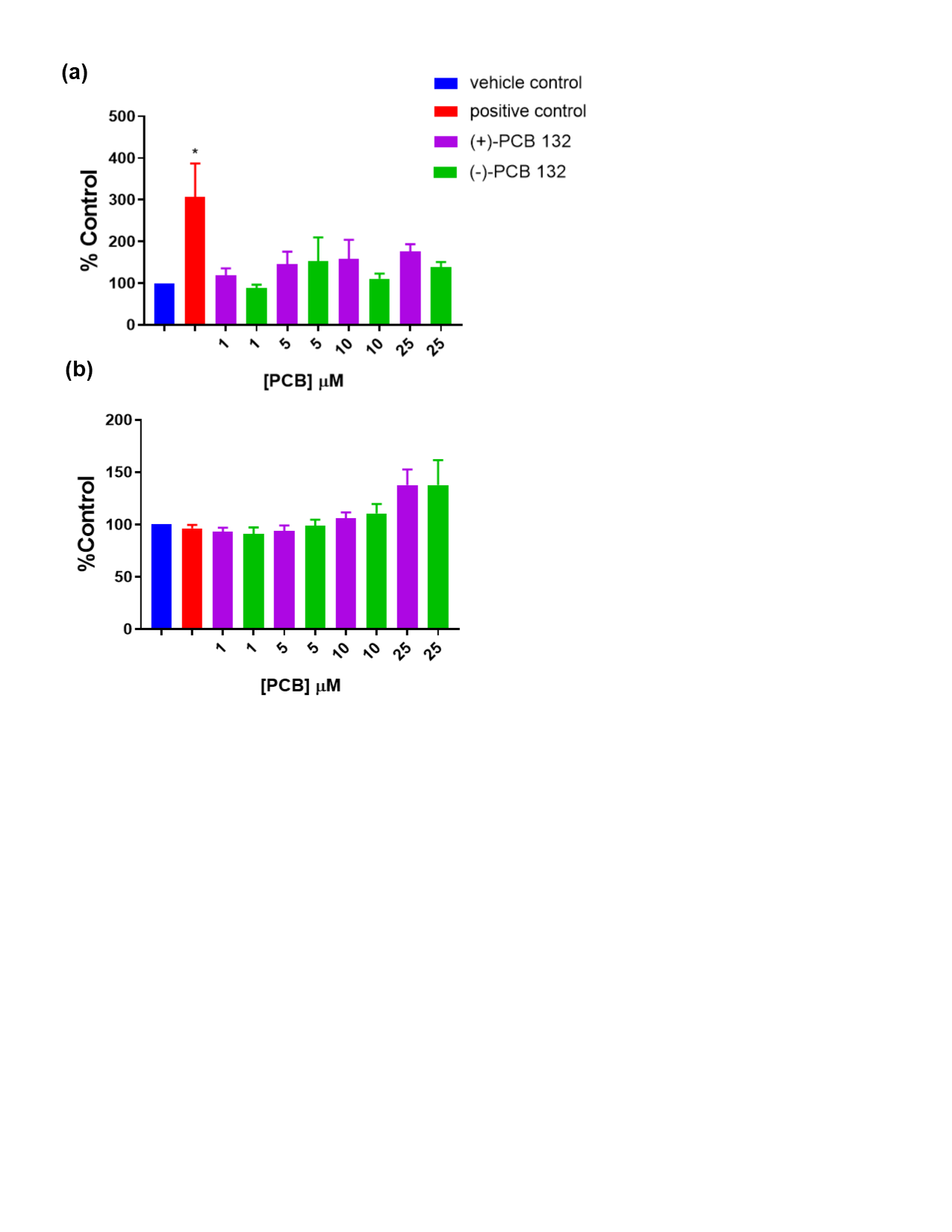

**Fig. S14.** Production of reactive oxygen species in N27 cells after 24 h exposure to PCB 132 atropisomers. (a) DCFDA detection of H_2_O_2_ revealed minimal increases in H_2_O_2_ production. These changes did not reach statistical significance. H_2_O_2_ (200 μM) was used as a positive control. (b) DHE detection of general ROS, with a positive control of 100 μM paraquat, which showed a moderate increase in ROS approximately 3h after treatment, but by 24 h this effect dissipated. A dose dependent increase in general ROS was observed with both PCB 132 atropisomers; however, this increase was not statistically significant. Data are present as mean ± standard error, n = 4.

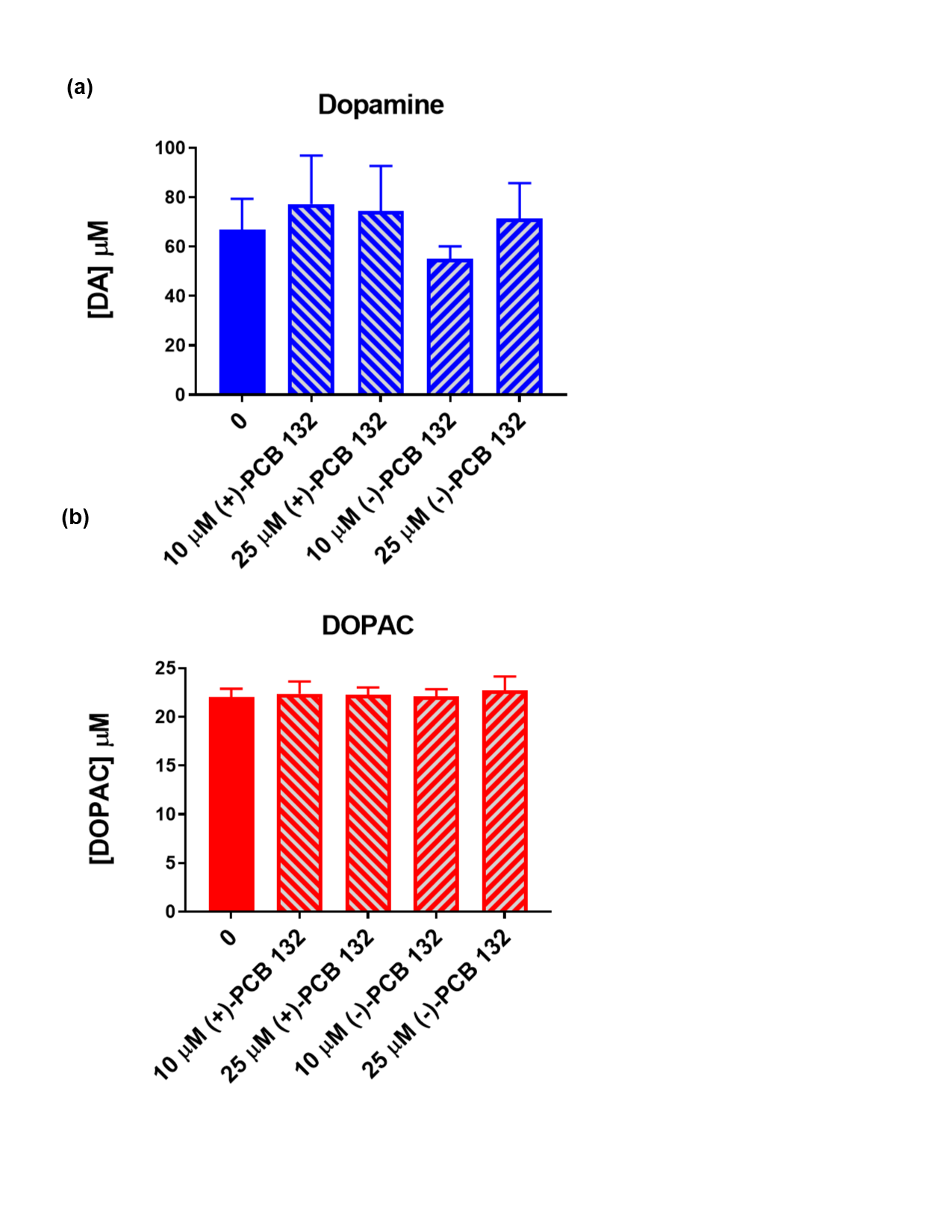

**Fig. S15. Levels of (a) dopamine (b) 3,4-dihydroxyphenylacetic acid (DOPAC) in media are not significantly changed following exposure of PC12 cells to PCB 132 atropisomers.** Extracellular measurements of dopamine and its metabolites were performed by HPLC analysis with photodiode array detection. DOPET presence in media was minimal and therefore not quantified here. Data are represented as mean ± standard error, n = 4. See **Fig. 9** for the corresponding intracellular levels of dopamine and its metabolites.

**Computational Data**

Absolute energies and cartesian coordinates for all species considered herein.

*Structures for 3,4-arene oxide mechanisms*

**R=H “3,4-arene oxide”**

Path Toward C-Cl

```
H 3,4 Reactant Complex (C-Cl same as C-H)
RM11 Energy	-1838.00794023 Hartree
Free Energy	-1837.923967 Hartree
6 0.497318 0.343222 0.038553
6 0.294046 -1.127606 -0.067567
6 -1.082473 -1.639308 -0.156028
6 -2.104326 -0.777583 -0.017595
17 -3.752337 -1.341601 -0.056688
6 -1.931338 0.667344 0.082264
6 -0.706024 1.216899 0.033450
17 -0.487594 2.933265 0.025648
1 -2.822191 1.303759 0.136715
1 -1.247941 -2.702577 -0.360607
1 1.116699 -1.716636 -0.500382
8 0.662258 -0.464513 1.163359
17 1.987691 1.042505 -0.610399
1 2.501580 -0.881643 1.545220
8 3.433120 -1.125985 1.402157
1 3.895552 -0.278273 1.326455
8 3.224347 -2.101649 -1.163085
1 3.126065 -1.299160 -1.693462
1 3.357301 -1.766981 -0.247397

H 3,4 TS1 (C-Cl)
RM11 Energy	-1837.97023307 Hartree
Free Energy	-1837.888569 Hartree
6 0.512861 0.215244 0.144288
6 0.155705 -1.206460 0.009360
6 -1.163245 -1.643736 0.067213
6 -2.155627 -0.684049 -0.001550
17 -3.792088 -1.180018 -0.014051
6 -1.916406 0.737109 -0.040384
6 -0.643807 1.178356 0.007957
17 -0.267978 2.842177 0.097996
1 -2.766235 1.428284 -0.070382
1 -1.410649 -2.709094 0.114002
1 1.013807 -1.892855 -0.062288
8 0.959421 0.070856 1.380038
17 1.806849 0.695971 -1.134351
1 2.619521 -0.406841 1.417020
8 3.565084 -0.649823 1.278299
1 3.914722 0.083202 0.750741
8 2.988276 -2.474063 -0.663800
1 2.831509 -1.892880 -1.421333
1 3.262877 -1.852566 0.051541

H 3,4 INT (C-Cl)
RM11 Energy	-1838.03972129 Hartree
Free Energy	-1837.957727 Hartree
6 -0.361760 0.894297 0.043015
6 -0.582293 -0.614165 0.006927
6 0.630005 -1.480435 0.002893
6 1.848421 -0.938552 -0.091172
17 3.274047 -1.935601 -0.161672
6 2.070389 0.505662 -0.139625
6 1.035827 1.362217 -0.085211
17 1.261273 3.077765 -0.104657
1 3.099493 0.879500 -0.211294
1 0.482659 -2.566656 0.036236
1 -1.189041 -0.825590 -0.900504
8 -1.294459 1.663106 0.150951
17 -1.665054 -1.079814 1.379525
1 -3.010281 1.019668 -0.091332
8 -3.863331 0.595678 -0.310233
1 -4.157125 0.206642 0.526366
8 -3.028556 -1.637224 -1.658390
1 -2.988930 -2.289750 -0.944847
1 -3.384054 -0.830379 -1.218538

H 3,4 TS2 (C-Cl)
RM11 Energy	-1838.03072424 Hartree
Free Energy	-1837.949308 Hartree
6 0.457018 -0.780964 -0.295376
6 0.599857 0.676746 -0.418711
6 -0.628712 1.413282 -0.090771
6 -1.824518 0.802286 0.103265
6 -1.936881 -0.627527 0.017786
6 -0.855150 -1.398970 -0.235226
1 -0.934040 -2.489867 -0.320348
17 -3.519663 -1.332583 0.234824
1 -2.719873 1.400805 0.313569
17 -0.497079 3.145159 -0.056143
8 1.668591 1.239769 -0.669560
1 0.893293 -0.855929 0.936962
17 1.701370 -1.711456 -1.167446
1 3.105652 0.459045 0.090052
8 3.685498 -0.005091 0.733347
1 4.037445 -0.764345 0.245109
8 1.627801 -0.923848 2.071282
1 1.749856 -1.864002 2.283528
1 2.504204 -0.600513 1.693759

H 3,4 Product Complex (C-Cl)
RM11 Energy	-1838.08376331 Hartree
Free Energy	-1838.000631 Hartree
6 0.356944 -0.759711 -0.638269
6 0.608342 0.618246 -0.510005
6 -0.494067 1.416889 -0.155468
6 -1.760120 0.890156 0.071293
6 -1.942857 -0.483777 -0.048903
6 -0.893217 -1.322512 -0.406368
1 -1.042464 -2.402885 -0.513247
17 -3.523588 -1.164782 0.242485
1 -2.590511 1.548917 0.348662
17 -0.245571 3.131724 0.008809
8 1.811220 1.165603 -0.712677
1 0.615011 -0.641084 2.156702
17 1.671068 -1.803435 -1.115875
1 2.522925 0.655264 -0.197829
8 3.429592 -0.058893 0.870064
1 3.913915 -0.785474 0.451291
8 1.397635 -1.204587 2.259799
1 1.155928 -2.031818 1.816046
1 2.735006 -0.495255 1.425090
```

**R=H**

Path Toward C-H

```
H 3,4 Reactant Complex (C-H same as C-Cl)
See above

H 3,4 TS1 (C-H)
RM11 Energy	-1837.93746772 Hartree
Free Energy	-1837.857243 Hartree
6 -0.416614 0.556317 0.078173
6 -0.467964 -0.928335 -0.087982
6 0.867914 -1.581478 0.037933
6 2.010522 -0.857174 0.092754
17 3.574441 -1.619929 0.121138
6 1.966788 0.565986 0.091965
6 0.772431 1.274462 0.049392
17 0.791541 3.005089 -0.007006
1 2.913611 1.125156 0.095582
1 0.891529 -2.678055 -0.019481
1 -1.186000 -1.361449 0.646588
8 -0.872316 -1.015679 -1.423835
17 -1.909887 1.344514 0.132440
1 -2.410713 -1.276180 -1.280247
8 -3.385156 -1.440210 -1.049356
1 -3.814566 -0.583656 -1.183004
8 -3.240656 -1.715091 1.644741
1 -3.215319 -0.794391 1.939285
1 -3.345613 -1.646729 0.663725

H 3,4 INT (C-H)
RM11 Energy	-1838.04959328 Hartree
Free Energy	-1837.966746 Hartree
6 -0.823154 0.271586 -0.508992
6 -0.239613 -1.044089 -0.861835
6 1.264054 -1.153189 -0.865712
6 1.984169 -0.033571 -0.200279
17 3.690822 -0.258189 0.009369
6 1.395443 1.107308 0.182683
6 -0.039605 1.260441 -0.019749
17 -0.711952 2.785340 0.429006
1 1.953198 1.929202 0.643352
1 1.579901 -1.196687 -1.929577
1 1.536930 -2.123007 -0.415311
8 -0.940511 -1.988681 -1.177105
17 -2.528496 0.445743 -0.720740
1 -2.348432 -2.202587 0.208676
8 -2.547226 -2.196426 1.161109
1 -3.119193 -1.425827 1.288411
8 0.037578 -1.517626 1.871254
1 -0.010213 -0.583958 2.118061
1 -0.895041 -1.765392 1.687831

H 3,4 TS2 (C-H)
RM11 Energy	-1838.03568112 Hartree
Free Energy	-1837.955340 Hartree
6 -0.899688 0.237367 -0.460589
6 -0.233026 -0.973136 -0.951160
6 1.191019 -1.051171 -0.644157
6 1.892707 0.148628 -0.265847
17 3.625751 0.071029 -0.187473
6 1.248751 1.281178 0.122623
6 -0.179320 1.291807 0.030183
17 -0.984278 2.730652 0.563591
1 1.780369 2.166859 0.483799
1 1.751628 -1.843885 -1.157495
1 0.940557 -1.522875 0.560198
8 -0.869620 -1.918901 -1.439044
17 -2.624920 0.291333 -0.618543
1 -1.857531 -2.398242 0.141845
8 -1.974843 -2.414611 1.111976
1 -2.561558 -1.667775 1.304153
8 0.526237 -1.887400 1.808772
1 0.528810 -1.048034 2.299189
1 -0.440548 -2.108031 1.649596

H 3,4 Product Complex (C-H)
RM11 Energy	-1838.06647643 Hartree
Free Energy	-1837.981605 Hartree
6 -0.549185 0.018366 -0.569833
6 0.028712 -1.268555 -0.562837
6 1.403528 -1.378296 -0.307478
6 2.155285 -0.247985 -0.021139
17 3.857412 -0.412879 0.301286
6 1.578469 1.019237 0.049621
6 0.221144 1.133634 -0.229171
17 -0.532125 2.699453 -0.106205
1 2.171534 1.901596 0.308083
1 1.855055 -2.375597 -0.318903
8 -0.684342 -2.366502 -0.787337
17 -2.210511 0.203517 -1.047563
1 -0.975977 0.335355 2.412256
1 -1.529581 -2.324551 -0.251661
8 -2.611730 -2.101758 0.964836
1 -3.513253 -1.967904 0.649019
8 -1.903656 0.303984 2.144706
1 -2.005243 1.030704 1.513385
1 -2.383632 -1.275347 1.447399
```

**R=Ph 3,4-arene oxide**

Path Toward C-Cl

```
Ph 3,4 Reactant Complex (C-Cl)
E(RM11) =  -2068.73951320
Sum of electronic and thermal Free Energies=        -2068.582639
 C,0,-1.5792703944,-0.3287825758,-0.114221297
 C,0,-1.9688041572,1.1026286325,-0.0929866756
 C,0,-0.8973159353,2.1069142365,-0.1139230907
 C,0,0.3837146662,1.7094931005,-0.0374498201
 Cl,0,1.6408890861,2.916129592,0.0195630143
 C,0,0.8094493648,0.2979507513,-0.0602746649
 C,0,-0.1317814014,-0.6661033123,-0.1691304715
 Cl,0,0.2965300021,-2.3377943436,-0.3178822957
 H,0,-1.1586866319,3.1660969882,-0.2110306292
 H,0,-2.9610540025,1.3632368573,-0.4907713779
 O,0,-2.0310799177,0.2486326413,1.0730581892
 Cl,0,-2.6943841702,-1.4897905805,-0.8520977766
 H,0,-3.8887308651,-0.0986758643,1.4529964186
 O,0,-4.8472005902,-0.1964704707,1.3126465643
 H,0,-4.9787213774,-1.1483235801,1.1902015482
 O,0,-5.057365676,0.9733146151,-1.1651176108
 H,0,-4.6490308139,0.3200408732,-1.7494533119
 H,0,-5.0403032786,0.5439012872,-0.2797114464
 C,0,2.2610294421,-0.0459716131,-0.0102421298
 C,0,2.7966680423,-0.6327016726,1.1401515102
 C,0,3.0853705446,0.2080195189,-1.1112680579
 C,0,4.1516356068,-0.9589468636,1.1903479524
 H,0,2.1429040688,-0.8357185976,1.9988022078
 C,0,4.4378696023,-0.1253555941,-1.060131103
 H,0,2.6615687588,0.6713477799,-2.012159602
 C,0,4.9727922069,-0.7063996743,0.0909492001
 H,0,4.5685199719,-1.4158106678,2.0965626492
 H,0,5.0801305985,0.0721442612,-1.9274276602
 H,0,6.0384262496,-0.9649917248,0.1308337667

Ph 3,4 TS1 (C-Cl)
E(RM11) =  -2068.70205520
Sum of electronic and thermal Free Energies=        -2068.546462
 C,0,-1.6254216681,-0.2203333335,0.1749546968
 C,0,-1.8903606241,1.2062159123,-0.0494510577
 C,0,-0.866133802,2.1449913036,-0.0384598207
 C,0,0.4363207713,1.6832066272,-0.0721906109
 Cl,0,1.689595791,2.842691053,-0.1830544015
 C,0,0.827143513,0.2822885068,-0.0098043267
 C,0,-0.1735859128,-0.6264899421,0.0884914092
 Cl,0,0.1351308686,-2.2919962485,0.3219488715
 H,0,-1.077734839,3.218638392,-0.0689563757
 H,0,-2.9531662403,1.4730015964,-0.156627433
 O,0,-2.1121460129,-0.2031319327,1.4048967825
 Cl,0,-2.5845076362,-1.2653864198,-1.0607031662
 H,0,-3.8225480258,-0.4295590299,1.4221758558
 O,0,-4.7858253875,-0.5926559462,1.2866975132
 H,0,-4.8135739476,-1.4485110201,0.8345561607
 O,0,-4.9960078705,1.1775310645,-0.7763106396
 H,0,-4.6075223949,0.6810316977,-1.5101919741
 H,0,-4.9899286995,0.5340281219,-0.028312093
 C,0,2.2635498149,-0.1191850191,-0.0147458859
 C,0,2.791843342,-0.7919115486,-1.1206806785
 C,0,3.0754625671,0.1577378447,1.0901167086
 C,0,4.1306560336,-1.1813998587,-1.1225323541
 H,0,2.1463071267,-1.0110354911,-1.9816287589
 C,0,4.4123048563,-0.2361945879,1.0850814211
 H,0,2.6548398964,0.6833994042,1.9578852481
 C,0,4.9407897302,-0.9039319002,-0.0209599477
 H,0,4.5436213693,-1.7068406636,-1.9925038278
 H,0,5.0463552019,-0.0203153259,1.9539024305
 H,0,5.9938431791,-1.2121222563,-0.023363746
```

**PCB 91 3,4-arene oxide**

Path Toward C-Cl

```
PCB 91 3,4 Reactant Complex (C-Cl)
E(RM11) =  -2987.61564014
Sum of electronic and thermal Free Energies=        -2987.480467
 C,0,2.3734557755,-0.4845762293,-0.0249695416
 C,0,2.8363287607,0.865945396,-0.4350230792
 C,0,1.8213430859,1.8853115948,-0.7329305574
 C,0,0.5236783869,1.539755184,-0.743614871
 Cl,0,-0.6802580321,2.7231913945,-1.1726492925
 C,0,0.0311263445,0.2133120207,-0.336076086
 C,0,0.9094058254,-0.7245552519,0.078770494
 Cl,0,0.3735958299,-2.2462482854,0.7017520709
 H,0,2.1368817526,2.9165485181,-0.9238441122
 H,0,3.8324982545,1.1805333409,-0.0894850511
 O,0,2.8733583631,-0.2830383535,-1.3115271247
 Cl,0,3.4117836129,-1.4408679198,1.0410111466
 H,0,4.7369802681,-0.7263579166,-1.4937742167
 O,0,5.6833157543,-0.80732506,-1.2799934609
 H,0,5.7667312031,-1.6900507303,-0.8899486822
 O,0,5.8779548735,0.9669675822,0.8076027094
 H,0,5.4332275671,0.49630012,1.5258242027
 H,0,5.8731262167,0.3232037122,0.0633260891
 C,0,-1.4339254484,-0.0540816143,-0.316533533
 C,0,-2.0353395649,-0.8132007993,-1.3226357452
 C,0,-2.2360095041,0.4421901247,0.7156430654
 C,0,-3.403654489,-1.0671815411,-1.3066161911
 H,0,-1.413629256,-1.2127312531,-2.1339676733
 C,0,-3.6055945213,0.2038144098,0.7580293179
 C,0,-4.1708010299,-0.5532000822,-0.2641904472
 H,0,-3.874950846,-1.6616239809,-2.0973649866
 H,0,-4.2175206868,0.6009489883,1.575566607
 Cl,0,-1.5025058687,1.3826429961,1.9875419856
 Cl,0,-5.8870306275,-0.8648253646,-0.2298000368

PCB 91 3,4 TS1 (C-Cl)
E(RM11) =  -2987.57707897
Sum of electronic and thermal Free Energies=        -2987.444131
 C,0,-1.6271565498,-0.2292697835,0.228125213
 C,0,-1.8941062207,1.1765671707,-0.1087683508
 C,0,-0.8729227792,2.1161315662,-0.1755207229
 C,0,0.4298020491,1.6559190321,-0.1766460472
 Cl,0,1.6896561832,2.8022022085,-0.3156757238
 C,0,0.8182837108,0.2605997839,-0.0286891679
 C,0,-0.174641121,-0.6429868803,0.1506893951
 Cl,0,0.1444670699,-2.2903941518,0.4708320597
 H,0,-1.085623525,3.1858353256,-0.2688787833
 H,0,-2.9590336849,1.4355410863,-0.2191664193
 O,0,-2.0995561886,-0.1272086938,1.4569598877
 Cl,0,-2.5931866534,-1.3630797155,-0.9214047652
 H,0,-3.8199330608,-0.3403845937,1.5013438331
 O,0,-4.7851513749,-0.4989694222,1.3790647565
 H,0,-4.8260637845,-1.3803531525,0.9800357044
 O,0,-5.0122352379,1.1507868359,-0.7769947981
 H,0,-4.6735655256,0.597199067,-1.4946061768
 H,0,-5.0016009557,0.5524471147,0.0075873646
 C,0,2.2553964472,-0.1275974727,-0.0263382549
 C,0,2.8908070007,-0.5041439756,1.1596345064
 C,0,2.9961710683,-0.1324527873,-1.213301995
 C,0,4.2313566141,-0.875626125,1.1659213126
 H,0,2.3181733949,-0.505302082,2.0960765931
 C,0,4.3375774433,-0.4984829628,-1.2359542022
 C,0,4.9366853717,-0.8679242469,-0.0351235797
 H,0,4.7291464613,-1.1702522987,2.0965160532
 H,0,4.9020583757,-0.4970564162,-2.1750804344
 Cl,0,2.2238284834,0.3332248801,-2.7053486147
 Cl,0,6.6162877585,-1.3345364103,-0.0433424732

PCB 91 3,4 INT (C-Cl)
E(RM11) =  -2987.64778472
Sum of electronic and thermal Free Energies=        -2987.513433
 C,0,2.3083706986,-0.9212924254,0.0181201421
 C,0,2.8739420462,0.4830590452,0.1560816924
 C,0,1.8924292212,1.5973592459,0.2374227114
 C,0,0.5843827889,1.3778593416,0.0759214183
 Cl,0,-0.5306814752,2.7152047695,0.0721941286
 C,0,0.0175854092,0.0299899031,-0.1332874331
 C,0,0.8446756266,-1.0380658278,-0.1737632375
 Cl,0,0.2564678536,-2.6523148357,-0.3723752563
 H,0,2.2840104685,2.6119241971,0.3797689128
 H,0,3.5399459379,0.6460167763,-0.7205207956
 O,0,3.0307659978,-1.8956517079,0.0480214349
 Cl,0,3.9848697067,0.5306573635,1.5823210981
 H,0,4.8570191871,-1.6242203376,-0.129579017
 O,0,5.7815262116,-1.3640960036,-0.3104958101
 H,0,6.1122479195,-1.0511893886,0.5439848008
 O,0,5.4150880246,1.0180673069,-1.6043305382
 H,0,5.6066935292,1.6609635615,-0.9066699285
 H,0,5.6067082725,0.1472223448,-1.1848524767
 C,0,-1.4520892464,-0.1313554701,-0.2959582083
 C,0,-2.000646678,-0.430399773,-1.5452591095
 C,0,-2.3101378479,0.0117833107,0.7985331915
 C,0,-3.3756329755,-0.5699245468,-1.704292273
 H,0,-1.3325328749,-0.5535021893,-2.4071928617
 C,0,-3.6877069588,-0.1263015534,0.6682390435
 C,0,-4.2010873035,-0.4135281715,-0.5933018702
 H,0,-3.8080192272,-0.7998717877,-2.6844718003
 H,0,-4.3458131054,-0.0134896883,1.5370372046
 Cl,0,-1.6372447178,0.3556383459,2.3689999442
 Cl,0,-5.9262154894,-0.5858388053,-0.7801601071

PCB 91 3,4 TS2 (C-Cl)
E(RM11) =  -2987.63877058
Sum of electronic and thermal Free Energies=        -2987.507451
 C,0,-2.0160572519,0.6442550833,-0.2679418546
 C,0,-1.564540037,-0.7338303688,-0.4921362323
 C,0,-0.1179794009,-0.9186071967,-0.2930190917
 C,0,0.7573406362,0.1165453782,-0.1501446578
 C,0,0.2573754292,1.4769359167,-0.1623569826
 C,0,-1.0633643275,1.7375710609,-0.2753738309
 H,0,-1.4369541219,2.7686523587,-0.2944178022
 Cl,0,1.4062339718,2.7838733495,-0.0289216541
 Cl,0,0.4391183724,-2.5610547964,-0.368484011
 O,0,-2.3271694014,-1.6762273548,-0.7126479105
 H,0,-2.3037234601,0.4736424774,0.9937614553
 Cl,0,-3.6137992771,1.0178350678,-0.9590046748
 H,0,-3.880708264,-1.5928145644,0.2273612474
 O,0,-4.5160514004,-1.4426393058,0.9605121049
 H,0,-5.2006045358,-0.8703840642,0.5830352171
 O,0,-2.8507285803,0.1593204008,2.195459867
 H,0,-3.3107616385,0.9470545097,2.5291912601
 H,0,-3.5638744189,-0.4731536277,1.8709347053
 C,0,2.2168062039,-0.13265352,0.0102158175
 C,0,3.0964608605,0.0660950824,-1.0575309764
 C,0,2.7356940629,-0.5644673197,1.2330330119
 C,0,4.4644392167,-0.1506410383,-0.9276743658
 C,0,4.1004257086,-0.7845049391,1.3928449561
 H,0,2.0516663045,-0.7288946201,2.0753319395
 C,0,4.9471038184,-0.5735335298,0.307261262
 H,0,5.1390240456,0.0077850305,-1.7763548309
 H,0,4.5078209396,-1.1184307263,2.3537987125
 Cl,0,2.466538698,0.5922054791,-2.5958432771
 Cl,0,6.6601576674,-0.848194363,0.4912719562

PCB 91 3,4 Product Complex (C-Cl)
E(RM11) =  -2987.69176651
Sum of electronic and thermal Free Energies=        -2987.556504
 C,0,2.8455271083,0.6680507111,0.2614939601
 C,0,2.354974123,-0.5885365498,0.6529769249
 C,0,0.971511527,-0.7947726112,0.4994022333
 C,0,0.110528056,0.1711598289,-0.029665332
 C,0,0.6709545501,1.3914789086,-0.4255636456
 C,0,2.027528502,1.6508985781,-0.2810201561
 H,0,2.4477273326,2.6160069088,-0.5851577055
 Cl,0,-0.350639409,2.6260811297,-1.1123189172
 Cl,0,0.3359833085,-2.3324085528,1.0062759489
 O,0,3.1326668552,-1.5543233206,1.1506102528
 H,0,2.6507856437,-0.599945493,-2.2312452094
 Cl,0,4.5424506573,1.0061338847,0.4721192764
 H,0,3.9541199687,-1.6813323183,0.5651118274
 O,0,4.9923625742,-1.9209015475,-0.5904063712
 H,0,5.7840211236,-1.3902738752,-0.4185004861
 O,0,3.5862652676,-0.5815325289,-2.4864968512
 H,0,3.8238003634,0.3577513933,-2.4549679178
 H,0,4.5274052997,-1.4562322788,-1.3313977472
 C,0,-1.3452852217,-0.0994765696,-0.1874172403
 C,0,-2.2911434081,0.4965739052,0.6532562879
 C,0,-1.8004887345,-0.9627854391,-1.1884832966
 C,0,-3.6546075071,0.25526675,0.5096837293
 C,0,-3.1569051484,-1.2216612632,-1.3565810495
 H,0,-1.0673550253,-1.4428843566,-1.8493953159
 C,0,-4.0679027498,-0.605348633,-0.5020392053
 H,0,-4.3797073754,0.7301648263,1.1796807399
 H,0,-3.5083046896,-1.8976371242,-2.1441909558
 Cl,0,-1.7646939695,1.5655389245,1.9263440925
 Cl,0,-5.7734580226,-0.9192912873,-0.695620871
```

**PCB 95 3,4-arene oxide**

Path Toward C-Cl

```
PCB 95 3,4 Reactant Complex (C-Cl)
E(RM11) =  -2987.61571829
Sum of electronic and thermal Free Energies=        -2987.480531
 C,0,-2.2088092227,-0.4731856501,0.0953093785
 C,0,-2.6809791138,0.7061325581,-0.6718333443
 C,0,-1.6708320889,1.616463215,-1.2296883286
 C,0,-0.375385,1.4272007719,-0.9323587357
 Cl,0,0.8214972626,2.5241135063,-1.5637703932
 C,0,0.125371795,0.27003537,-0.169113637
 C,0,-0.7436645704,-0.671649045,0.2567588705
 Cl,0,-0.2099782262,-2.0922627623,1.0869935739
 H,0,-1.9873664458,2.4164769433,-1.9074953852
 H,0,-3.6630084918,0.6271806105,-1.1626658624
 O,0,-2.7535173006,0.6211719811,0.7689275412
 Cl,0,-3.2106426588,-1.9307847663,0.1020015219
 H,0,-4.6175553927,0.3980248659,1.196328242
 O,0,-5.5584143766,0.1707372769,1.0914607235
 H,0,-5.6432348794,-0.7024303847,1.5016796276
 O,0,-5.6733611861,-0.2245091928,-1.6263259013
 H,0,-5.2140125997,-1.0669188867,-1.746549767
 H,0,-5.6936724152,-0.1004586756,-0.6503079322
 C,0,1.5891292903,0.1339535908,0.0724453898
 C,0,2.3380560264,-0.8059636383,-0.6373593912
 C,0,2.2362989828,0.9486615184,1.0066569026
 C,0,3.7052085795,-0.9100132902,-0.402473134
 H,0,1.8480783503,-1.456152504,-1.3726411221
 C,0,3.604518116,0.8401358669,1.2381970834
 C,0,4.3484308939,-0.0970022183,0.5278022956
 H,0,4.0856289689,1.4909213387,1.9774105802
 H,0,5.4269464217,-0.1968364358,0.6967277377
 Cl,0,1.30630749,2.1233472205,1.8992966507
 Cl,0,4.6320547915,-2.087625184,-1.2973901849

PCB 95 3,4 TS1 (C-Cl)
E(RM11) =  -2987.57718218
Sum of electronic and thermal Free Energies=        -2987.444065
 C,0,-1.633825303,-0.2301845118,0.1342479271
 C,0,-1.8918356769,1.214397027,0.0586799045
 C,0,-0.8675748324,2.1460149671,0.1774801963
 C,0,0.4333843377,1.6876891982,0.1003912537
 Cl,0,1.7007696174,2.8331643108,0.1325644951
 C,0,0.8132652817,0.2866645078,0.0074096906
 C,0,-0.1826108039,-0.6298445409,0.0069295865
 Cl,0,0.1349055596,-2.3061404139,0.0672371885
 H,0,-1.0767328461,3.2168566175,0.2666582885
 H,0,-2.9537900933,1.4938590094,-0.0229688298
 O,0,-2.1222418518,-0.3533412307,1.3545055887
 Cl,0,-2.5868944842,-1.1337662309,-1.2162940314
 H,0,-3.8383071227,-0.5781957202,1.3351857683
 O,0,-4.7976762967,-0.7269492631,1.1637947802
 H,0,-4.8125450787,-1.5254143241,0.6159208526
 O,0,-4.9718702381,1.2460287237,-0.7104171378
 H,0,-4.5640672471,0.8213391315,-1.4782442316
 H,0,-4.981979623,0.5333586698,-0.0281563034
 C,0,2.2478317345,-0.1107213208,-0.0263203782
 C,0,2.8309828322,-0.5329520641,-1.2217036289
 C,0,3.0240781497,-0.0828089313,1.1371352266
 C,0,4.1706994152,-0.9087014223,-1.2310473123
 H,0,2.2354663151,-0.5668160787,-2.1423522478
 C,0,4.3633967725,-0.4597515954,1.1211580435
 C,0,4.9438772923,-0.8763871751,-0.0730895016
 H,0,4.9500817586,-0.4279642881,2.0464309906
 H,0,5.9976817149,-1.1770032845,-0.1045334124
 Cl,0,2.2949001267,0.4364047949,2.633451355
 Cl,0,4.8938157997,-1.4295234617,-2.730602801

PCB 95 3,4 INT (C-Cl)
E(RM11) =  -2987.64767120
Sum of electronic and thermal Free Energies=        -2987.513163
 C,0,-2.1487158398,-0.8255479141,0.347565813
 C,0,-2.703536676,0.0571716955,-0.7581427268
 C,0,-1.7111260011,0.7525807837,-1.6213936356
 C,0,-0.40891696,0.7377568869,-1.3232333019
 Cl,0,0.7278020658,1.587948266,-2.3311764757
 C,0,0.1384552972,0.0003649226,-0.1667331057
 C,0,-0.6934319458,-0.7296186588,0.608465213
 Cl,0,-0.1280028714,-1.646185138,1.9615051715
 H,0,-2.0879281341,1.3085404314,-2.4883736258
 H,0,-3.3542565629,0.8104091296,-0.2602879347
 O,0,-2.871539347,-1.5427328408,1.0066173252
 Cl,0,-3.8414343285,-0.9081198682,-1.7756278896
 H,0,-4.7026576277,-1.284463875,0.9493867977
 O,0,-5.6365819778,-0.9984108207,0.9186765751
 H,0,-6.0027811869,-1.4576221673,0.1487828405
 O,0,-5.3255597728,1.5601989497,-0.0123618047
 H,0,-5.4466020318,1.448078777,-0.9659203821
 H,0,-5.4897904112,0.6601960828,0.3543364234
 C,0,1.5984146261,0.0624615769,0.1155677486
 C,0,2.4126201604,-1.0414012565,-0.141487626
 C,0,2.1708983087,1.2258305005,0.6387345813
 C,0,3.7765940702,-0.9555782702,0.1185801871
 H,0,1.9772594729,-1.9641351413,-0.5444481195
 C,0,3.5355126873,1.3034749811,0.8997346762
 C,0,4.3477379521,0.2050008503,0.6357048742
 H,0,3.9603321528,2.2258613549,1.3121537754
 H,0,5.4248266388,0.2490188013,0.8341142198
 Cl,0,1.1500921585,2.5973190909,0.9789635294
 Cl,0,4.7908810838,-2.3364291302,-0.2098031232

PCB 95 3,4 TS2 (C-Cl)
E(RM11) =  -2987.63901027
Sum of electronic and thermal Free Energies=        -2987.506452
 C,0,-2.0169137229,0.6486020701,-0.2510933024
 C,0,-1.5520519508,-0.7187238064,-0.5111335366
 C,0,-0.1047597155,-0.8975075308,-0.3115668456
 C,0,0.7599977256,0.1398855907,-0.1286426261
 C,0,0.2485028025,1.4955240162,-0.09269132
 C,0,-1.0733309076,1.7494333669,-0.2101986772
 H,0,-1.4561158957,2.7773814333,-0.1914377323
 Cl,0,1.385317054,2.8040733239,0.1101711158
 Cl,0,0.4680948581,-2.53059563,-0.4431116756
 O,0,-2.3047994356,-1.6605869205,-0.7648346566
 H,0,-2.3165076475,0.444676649,0.9988139826
 Cl,0,-3.6110466782,1.0264018986,-0.9484636362
 H,0,-3.8622580187,-1.6304088963,0.1761269538
 O,0,-4.5085951956,-1.4949690146,0.9022716123
 H,0,-5.1840599612,-0.9109599128,0.5260230739
 O,0,-2.8834781199,0.1068488088,2.189316753
 H,0,-3.3629440963,0.8851599216,2.5178429042
 H,0,-3.5807516523,-0.5300196933,1.839545258
 C,0,2.2217740479,-0.1048752958,0.0230479112
 C,0,2.728405983,-0.6029052415,1.2243444932
 C,0,3.1036581311,0.1540142091,-1.0302747624
 C,0,4.096594103,-0.8217353533,1.3494720527
 H,0,2.0499657323,-0.8207202828,2.0584306055
 C,0,4.4715317159,-0.0690288633,-0.8995068089
 C,0,4.9755244878,-0.5595044399,0.3011607788
 H,0,5.140918748,0.1411124459,-1.7417442929
 H,0,6.0497901153,-0.7407598474,0.4226277548
 Cl,0,4.7220740761,-1.4422479129,2.856038912
 Cl,0,2.4746151974,0.7581977075,-2.539862539

PCB 95 3,4 Product Complex (C-Cl)
E(RM11) =  -2987.69191399
Sum of electronic and thermal Free Energies=        -2987.557088
 C,0,2.7417298013,0.6598003336,-0.044699305
 C,0,2.2183474285,-0.3084649904,0.8279634608
 C,0,0.8169507365,-0.4365895835,0.8509969902
 C,0,-0.0310373044,0.3338968021,0.050210065
 C,0,0.5587312139,1.2644005315,-0.8131713135
 C,0,1.935599747,1.4368966751,-0.866534497
 H,0,2.3807976245,2.1761907845,-1.5416216635
 Cl,0,-0.4502862624,2.2407359048,-1.8467774988
 Cl,0,0.140556541,-1.6161238544,1.9361240455
 O,0,2.9845159543,-1.0804135837,1.6039854362
 H,0,2.2156704765,-1.4206606785,-1.8619397633
 Cl,0,4.4682482663,0.8942723165,-0.0921278424
 H,0,3.7266977405,-1.5013483763,1.050067285
 O,0,4.6128902584,-2.2536649646,-0.0031700801
 H,0,5.4529705881,-1.7822593282,-0.1038595953
 O,0,3.1212045249,-1.5891184841,-2.1658608611
 H,0,3.4249919382,-0.7315364624,-2.5003174892
 H,0,4.1061866465,-2.0555643025,-0.8314932004
 C,0,-1.5090977839,0.153941937,0.0976253059
 C,0,-2.0930918392,-0.972569967,-0.4862483603
 C,0,-2.3378632423,1.0955993042,0.7167879725
 C,0,-3.4740411911,-1.1321398032,-0.4454442352
 H,0,-1.4604269667,-1.7265458609,-0.9707640449
 C,0,-3.7203824621,0.9306820598,0.7551860064
 C,0,-4.2970649287,-0.1915159687,0.1693883628
 H,0,-4.3448037537,1.6846796901,1.2480431532
 H,0,-5.3832870219,-0.3370497523,0.1912966399
 Cl,0,-4.1882066366,-2.5450562641,-1.1822381623
 Cl,0,-1.6334510931,2.5056888856,1.4636181889
```

**PCB 132 3**'**,4**'**-arene oxide**

Path Toward C-Cl

**PCB 132 3**'**,4**' **Reactant Complex (C-Cl)**

```
RM11 Energy	-3447.04642054 Hartree
Free Energy	-3446.922920 Hartree
6 -2.688583 0.470052 0.119093
6 -1.248514 0.824291 0.008356
6 -0.355873 -0.078614 -0.450735
6 -0.833286 -1.366943 -0.982596
6 -2.129442 -1.712120 -1.038635
6 -3.137339 -0.836821 -0.423894
1 -4.185140 -0.910903 -0.752256
8 -6.284265 -0.195646 -1.100827
1 -5.890548 0.659014 -1.323077
1 -6.172929 -0.263711 -0.125527
8 -5.798862 -0.508957 1.584779
1 -6.003324 -1.444223 1.734487
1 -4.827571 -0.468308 1.622574
8 -2.994531 -0.577265 0.989508
1 -2.453996 -2.618722 -1.560519
17 0.370870 -2.422783 -1.668127
6 1.101764 0.228485 -0.479912
6 1.961978 -0.380648 0.438554
6 3.335005 -0.109906 0.423176
6 3.836050 0.782239 -0.531842
6 2.986529 1.393676 -1.449481
6 1.625506 1.116470 -1.419686
1 0.951166 1.596894 -2.139425
1 3.403677 2.088566 -2.186991
17 5.533585 1.138943 -0.584874
17 4.390038 -0.862981 1.567294
17 1.304697 -1.484048 1.605069
17 -0.756209 2.385167 0.566868
17 -3.838443 1.812488 0.141339
```

**PCB 132 3**'**,4**' **TS1 (C-Cl)**

```
RM11 Energy	-3447.00704296 Hartree
Free Energy	-3446.886097 Hartree
6 2.719384 -0.352919 0.209221
6 1.257509 -0.711985 0.076860
6 0.357097 0.143532 -0.461860
6 0.853703 1.419503 -0.949103
6 2.178720 1.811404 -0.916101
6 3.116539 0.890874 -0.467291
1 4.201530 1.065555 -0.524118
8 6.239017 0.392358 -0.986597
1 5.806483 -0.376858 -1.382898
1 6.144210 0.238795 -0.016483
8 5.768606 0.087795 1.674655
1 5.974281 0.973767 2.007189
1 4.783788 0.062866 1.642636
8 3.060869 0.068252 1.413198
1 2.478328 2.796956 -1.286418
17 -0.300501 2.506574 -1.585493
6 -1.095668 -0.178800 -0.514139
6 -1.973330 0.384437 0.417945
6 -3.338387 0.079298 0.389023
6 -3.811138 -0.803821 -0.589524
6 -2.942903 -1.371760 -1.517141
6 -1.589604 -1.059467 -1.475938
1 -0.898753 -1.505617 -2.201851
1 -3.338619 -2.060950 -2.271506
17 -5.497638 -1.203994 -0.657219
17 -4.416856 0.777282 1.544851
17 -1.347344 1.471900 1.615700
17 0.777547 -2.168365 0.825707
17 3.720633 -1.800076 -0.448740
```

**PCB 132 3**'**,4**' **INT (C-Cl)**

```
RM11 Energy	-3447.07787517 Hartree
Free Energy	-3446.955895 Hartree
6 -3.148408 -0.417974 -0.308186
6 -2.596351 0.955048 0.038708
6 -1.120889 1.097761 0.031465
6 -0.298667 0.119899 -0.407564
6 -0.874917 -1.150844 -0.891214
6 -2.186838 -1.402186 -0.874472
1 -2.585928 -2.355759 -1.240462
17 0.234371 -2.333784 -1.524923
6 1.176897 0.313804 -0.439837
6 1.990570 -0.355456 0.478450
6 3.380351 -0.194754 0.450606
6 3.942307 0.651458 -0.512680
6 3.136905 1.325908 -1.426008
6 1.758539 1.157992 -1.385832
1 1.117569 1.687570 -2.101577
1 3.601642 1.984310 -2.168474
17 5.661980 0.873606 -0.578504
17 4.380605 -1.027019 1.587889
17 1.253763 -1.385262 1.663945
17 -0.528367 2.625562 0.582931
8 -3.327547 1.874190 0.339044
1 -3.583493 -0.829294 0.629869
17 -4.568809 -0.225464 -1.404148
8 -5.390121 -1.558086 1.542781
1 -5.659185 -1.921260 0.687394
1 -5.615846 -0.601022 1.474832
8 -5.875607 1.126353 1.315584
1 -5.769858 1.472124 2.214315
1 -5.085938 1.449864 0.841987
```

**PCB 132 3**'**,4**' **TS2 (C-Cl)**

```
RM11 Energy	-3447.06931361 Hartree
Free Energy	-3446.948683 Hartree
6 -3.158086 -0.442674 -0.510320
6 -2.700608 0.935244 -0.302775
6 -1.235030 1.065301 -0.239552
6 -0.379452 0.063230 -0.587214
6 -0.916896 -1.213597 -1.015774
6 -2.247995 -1.443215 -1.034608
1 -2.650838 -2.404873 -1.375263
17 0.200019 -2.448861 -1.538914
6 1.095306 0.271846 -0.553257
6 1.869928 -0.329966 0.442369
6 3.257350 -0.145933 0.478016
6 3.858150 0.653594 -0.501142
6 3.092767 1.259418 -1.493602
6 1.717028 1.068808 -1.514970
1 1.106928 1.546788 -2.291410
1 3.587048 1.882683 -2.247245
17 5.575700 0.903739 -0.488093
17 4.209175 -0.891336 1.714094
17 1.091312 -1.305457 1.647584
17 -0.651805 2.628854 0.235766
8 -3.456244 1.882846 -0.084365
1 -3.273793 -0.737180 0.754879
17 -4.851533 -0.574921 -1.041894
8 -3.702550 -0.896942 2.037646
1 -4.179062 -1.742367 2.077862
1 -4.404243 -0.173065 2.059780
8 -5.380927 1.097739 1.747871
1 -5.227931 1.749068 2.449311
1 -4.893872 1.453151 0.974600
```

**PCB 132 3**'**,4**' **Product Complex (C-Cl) == 3**'**-140**

```
RM11 Energy	-3447.12335022 Hartree
Free Energy	-3446.998286 Hartree
6 -3.219217 -0.228782 -0.797361
6 -2.709541 0.993006 -0.327432
6 -1.310533 1.127520 -0.317297
6 -0.449726 0.109645 -0.737975
6 -1.021968 -1.084648 -1.189191
6 -2.398792 -1.262761 -1.229832
1 -2.832197 -2.202734 -1.588904
17 0.012980 -2.387884 -1.707488
6 1.027297 0.279640 -0.660800
6 1.755184 -0.334990 0.363371
6 3.144026 -0.178120 0.451455
6 3.796712 0.610339 -0.503041
6 3.080789 1.228348 -1.524191
6 1.703559 1.062725 -1.597194
1 1.133556 1.552052 -2.396609
1 3.614471 1.840891 -2.259525
17 5.516640 0.832865 -0.422330
17 4.035566 -0.941188 1.721912
17 0.923514 -1.301019 1.544427
17 -0.650006 2.614720 0.294563
8 -3.499702 1.975648 0.122703
1 -4.089751 1.593894 0.857852
17 -4.945466 -0.447239 -0.815556
1 -2.033029 -0.914726 1.706401
8 -2.842081 -1.235466 2.134071
1 -3.134189 -1.968367 1.570774
1 -4.035626 0.034666 2.140866
8 -4.637607 0.820381 2.123072
1 -4.359732 1.353939 2.883338
```

Path Toward C-H

**PCB 132 3**'**,4**' **Reactant Complex (C-H same as C-Cl)**

See above

**PCB 132 3**'**,4**' **TS1 (C-H)**

```
RM11 Energy	-3446.97455562 Hartree
Free Energy	-3446.855541 Hartree
6 2.588319 0.650510 -0.047573
6 1.228944 0.916127 -0.055135
6 0.348996 -0.048642 0.461444
6 0.833143 -1.308184 0.942108
6 2.155717 -1.602735 0.904401
6 3.156648 -0.691476 0.288729
1 4.038298 -0.553917 0.957858
8 6.133419 0.391036 1.441644
1 5.819720 1.295838 1.313128
1 6.131148 0.002152 0.532327
8 5.988648 -0.768582 -1.007397
1 6.359564 -1.642178 -0.817071
1 4.990772 -0.935484 -1.083574
8 3.453793 -1.206979 -0.975133
1 2.519608 -2.583486 1.238875
17 -0.311143 -2.456674 1.572169
6 -1.106524 0.241600 0.495470
6 -1.967037 -0.390785 -0.405123
6 -3.340867 -0.131552 -0.368691
6 -3.833315 0.765057 0.587651
6 -2.975239 1.401490 1.480678
6 -1.611040 1.147811 1.428539
1 -0.927234 1.650913 2.123522
1 -3.386856 2.102852 2.215043
17 -5.531312 1.100700 0.667189
17 -4.403755 -0.904885 -1.488165
17 -1.307885 -1.476876 -1.583453
17 0.613624 2.400843 -0.700243
17 3.704097 1.786551 -0.610322
```

**PCB 132 3**'**,4**' **Product Complex (C-H) == PCB 132 4**'**,5**'**-arene oxide**

```
RM11 Energy	-3447.04466215 Hartree
Free Energy	-3446.921255 Hartree
6 -2.462040 -1.256601 -0.615311
6 -1.008412 -1.043336 -0.757334
6 -0.396721 0.090137 -0.354777
6 -1.216392 1.222683 0.104379
6 -2.564162 1.164369 0.121022
6 -3.273397 -0.098913 -0.157272
1 -4.333187 -0.023379 -0.442671
8 -2.962973 -1.183610 0.712874
17 -3.555471 2.533089 0.485292
17 -0.390117 2.681557 0.547732
6 1.082826 0.243075 -0.449229
6 1.930719 -0.406941 0.453958
6 3.321716 -0.272076 0.352710
6 3.853251 0.524612 -0.667140
6 3.017190 1.175004 -1.570435
6 1.640804 1.033103 -1.456380
1 0.976860 1.546255 -2.163477
1 3.456887 1.792435 -2.361832
17 5.571848 0.711989 -0.821076
17 4.365293 -1.077304 1.472515
17 1.253455 -1.394878 1.709160
17 -0.110752 -2.353520 -1.450792
1 -2.926632 -2.028034 -1.247253
8 -6.561272 0.014211 -0.429589
1 -6.377815 0.510752 0.380501
1 -6.232351 -0.890206 -0.228607
8 -5.432194 -2.439076 0.113505
1 -5.140555 -2.822306 -0.726992
1 -4.606333 -2.184261 0.562749
```

**PCB 136 3,4-arene oxide**

Pathway Toward C-Cl

**PCB 136 3,4 Reactant Complex (C-Cl)**

RM11 energy (hartrees): -3447.0498104

Free Energy (hartrees): -3446.926824

6 2.312419 0.091420 -0.415334
6 0.894645 0.524723 -0.523530
6 -0.030734 0.041787 0.331588
6 0.386048 -0.794742 1.467396
6 1.664736 -1.119582 1.716167
6 2.703835 -0.752989 0.743815
1 3.752326 -0.697583 1.073292
8 5.900856 -0.034771 1.012806
1 5.550056 0.830978 0.763355
1 5.761735 -0.589950 0.212384
8 5.340897 -1.673516 -1.119269
1 5.520285 -2.556305 -0.762382
1 4.369959 -1.623285 -1.156584
8 2.548607 -1.271910 -0.594474
1 1.952104 -1.629225 2.642083
17 -0.865494 -1.269452 2.581165
6 -1.470122 0.385529 0.172452
6 -2.357395 -0.557243 -0.359192
6 -3.712356 -0.251067 -0.509711
6 -4.191125 0.998817 -0.124923
6 -3.324964 1.942257 0.413372
6 -1.976875 1.629035 0.559426
17 -0.900100 2.811263 1.245858
1 -3.694215 2.925964 0.723964
1 -5.256115 1.227236 -0.248575
17 -4.811288 -1.418286 -1.172970
17 -1.745128 -2.115430 -0.811237
17 0.462452 1.607211 -1.797123
17 3.526165 1.156816 -1.132116

**PCB 136 3,4 TS1 (C-Cl)**

RM11 energy (hartrees): -3447.00987092

Free Energy (hartrees): -3446.889078

6 2.339893 -0.070433 -0.431243
6 0.910569 0.411502 -0.511752
6 -0.036464 -0.016563 0.355650
6 0.377740 -0.924107 1.412211
6 1.667773 -1.387378 1.584680
6 2.658767 -0.871356 0.759080
1 3.729120 -1.080246 0.905664
8 5.824395 -0.391790 1.031489
1 5.446320 0.498233 1.010371
1 5.717959 -0.711353 0.103980
8 5.321299 -1.356861 -1.458242
1 5.469788 -2.306128 -1.337410
1 4.340439 -1.261164 -1.455504
8 2.626126 -1.037351 -1.284132
1 1.904766 -2.089548 2.390268
17 -0.830857 -1.453738 2.497230
6 -1.460992 0.387173 0.212140
6 -2.400187 -0.512158 -0.308926
6 -3.736601 -0.133563 -0.457754
6 -4.141949 1.148185 -0.093164
6 -3.221618 2.054565 0.417028
6 -1.892398 1.669836 0.564466
17 -0.742281 2.814929 1.190688
1 -3.533119 3.065499 0.702466
1 -5.193386 1.433039 -0.215634
17 -4.902344 -1.245298 -1.099450
17 -1.875114 -2.099764 -0.767457
17 0.519957 1.380215 -1.858746
17 3.451600 1.439272 -0.517066

**PCB 136 3,4 INT (C-Cl)**

RM11 energy (hartrees): -3447.08085712

Free Energy (hartrees): -3446.958509

6 2.242214 0.589209 -0.575939
6 0.770368 0.735889 -0.679728
6 -0.073568 0.209679 0.233020
6 0.464250 -0.558895 1.370684
6 1.768089 -0.813572 1.513800
6 2.745038 -0.384569 0.475818
1 3.087221 -1.285266 -0.085862
8 4.750939 -2.503619 -0.563047
1 5.060627 -2.380075 0.345262
1 5.049100 -1.686510 -1.027320
8 5.444178 -0.167558 -1.805583
1 5.301996 -0.322482 -2.751265
1 4.711782 0.423082 -1.548011
8 2.998890 1.175373 -1.319174
1 2.145035 -1.391882 2.366076
17 -0.684235 -1.119861 2.552283
6 -1.544657 0.397827 0.110580
6 -2.345015 -0.660926 -0.332324
6 -3.727598 -0.500308 -0.451623
6 -4.317636 0.718352 -0.124750
6 -3.535847 1.776131 0.322540
6 -2.159103 1.609064 0.437527
17 -1.183480 2.931514 1.008380
1 -3.994306 2.736012 0.585437
1 -5.403341 0.833585 -0.223217
17 -4.720267 -1.810143 -1.006584
17 -1.587863 -2.170819 -0.723537
17 0.208101 1.695496 -1.999844
17 4.248368 0.258088 1.230698

**PCB 136 3,4 TS2 (C-Cl)**

RM11 energy (hartrees): -3447.07263975

Free Energy (hartrees): -3446.952060

6 2.765196 -0.114431 0.684989
6 2.354636 0.623099 -0.516676
6 0.893488 0.698492 -0.684916
6 0.013459 0.342985 0.291426
6 0.506982 -0.155387 1.558736
6 1.829946 -0.320897 1.775537
1 2.205427 -0.688656 2.738205
17 -0.655474 -0.524261 2.806123
6 -1.454610 0.454833 0.068674
6 -2.208630 -0.691996 -0.204111
6 -3.586813 -0.598904 -0.415864
6 -4.219724 0.640083 -0.353449
6 -3.485344 1.786386 -0.076255
6 -2.113465 1.684948 0.133547
17 -1.196922 3.120470 0.489522
1 -3.976542 2.764217 -0.021064
1 -5.300860 0.699534 -0.523555
17 -4.524979 -2.018048 -0.757461
17 -1.403530 -2.226926 -0.271235
17 0.343444 1.368696 -2.184896
8 3.140725 1.034462 -1.369206
1 2.820615 -1.270074 0.106452
17 4.463413 0.136687 1.148482
8 3.167960 -2.395488 -0.602703
1 3.593013 -3.010216 0.017404
1 3.905429 -1.995079 -1.157869
8 4.965496 -1.001245 -1.914191
1 4.816515 -1.108019 -2.866075
1 4.531076 -0.151308 -1.693629

**PCB 136 3,4 Product Complex (C-Cl)**

RM11 energy (hartrees): -3447.12702341

Free Energy (hartrees): -3447.005032

6 2.794447 0.184560 0.752760
6 2.343215 0.664578 -0.489211
6 0.950577 0.755741 -0.654194
6 0.044821 0.387624 0.343887
6 0.557241 -0.092654 1.552194
6 1.925755 -0.195683 1.767375
1 2.316212 -0.568996 2.720493
17 -0.539126 -0.574019 2.817970
6 -1.421256 0.477671 0.109651
6 -2.169380 -0.672424 -0.171251
6 -3.547191 -0.586227 -0.392262
6 -4.187602 0.649113 -0.338473
6 -3.460241 1.800306 -0.063768
6 -2.089919 1.704494 0.157244
17 -1.182163 3.149902 0.498890
1 -3.955365 2.776714 -0.019875
1 -5.267899 0.701963 -0.515915
17 -4.478707 -2.009572 -0.738226
17 -1.364569 -2.208352 -0.244831
17 0.352635 1.342567 -2.176126
8 3.171619 1.014953 -1.476915
1 2.020955 -2.191300 -0.512918
17 4.510508 0.055218 1.014954
8 2.880782 -2.640000 -0.502612
1 3.135666 -2.643904 0.432551
1 4.014472 -1.699518 -1.426915
8 4.575805 -1.083887 -1.961236
1 4.348981 -1.281841 -2.882634
1 3.842699 0.268105 -1.647143

Pathway Toward C-H

**PCB 136 3,4 Reactant Complex (C-H)**

RM11 energy (hartrees): -3447.04981093

Free Energy (hartrees): -3446.926751

6 2.312252 0.090383 -0.416920
6 0.894476 0.523878 -0.524515
6 -0.030610 0.041100 0.330972
6 0.386440 -0.795916 1.466401
6 1.665117 -1.121355 1.714359
6 2.703889 -0.754408 0.741748
1 3.752372 -0.698956 1.071210
8 5.900053 -0.030777 1.014398
1 5.546649 0.834030 0.765372
1 5.761465 -0.586456 0.214238
8 5.342491 -1.673181 -1.115325
1 5.517168 -2.554445 -0.752388
1 4.371983 -1.620645 -1.159267
8 2.548483 -1.272872 -0.596712
1 1.952790 -1.631705 2.639788
17 -0.864769 -1.270608 2.580526
6 -1.469960 0.385445 0.172711
6 -2.357826 -0.556671 -0.359110
6 -3.712733 -0.249892 -0.508781
6 -4.190855 0.999902 -0.122884
6 -3.324120 1.942675 0.415627
6 -1.976059 1.628880 0.560769
17 -0.898438 2.810235 1.247390
1 -3.692877 2.926292 0.727084
1 -5.255827 1.228778 -0.245858
17 -4.812396 -1.416218 -1.172377
17 -1.746297 -2.114792 -0.812375
17 0.461927 1.606715 -1.797697
17 3.525736 1.156057 -1.133781

**PCB 136 3,4 TS1 (C-H)**

RM11 energy (hartrees): -3446.97691403

Free Energy (hartrees): -3446.856201

6 2.217135 0.317223 -0.416911
6 0.872733 0.631940 -0.533036
6 -0.034704 0.043398 0.359631
6 0.393628 -0.902785 1.341917
6 1.699762 -1.258627 1.434951
6 2.726405 -0.765258 0.479559
1 3.637335 -0.402976 1.010545
8 5.782107 0.546488 1.027773
1 5.522537 1.338860 0.539166
1 5.726504 -0.181023 0.361013
8 5.498023 -1.522939 -0.713790
1 5.852532 -2.246844 -0.177654
1 4.496758 -1.667450 -0.689850
8 2.947263 -1.804659 -0.428991
1 2.025011 -2.017439 2.159374
17 -0.799080 -1.600764 2.397256
6 -1.477645 0.384264 0.252549
6 -2.365752 -0.513084 -0.346813
6 -3.720783 -0.189637 -0.446639
6 -4.186120 1.024166 0.054860
6 -3.309518 1.921915 0.651975
6 -1.960271 1.598528 0.743095
17 -0.843611 2.708210 1.480622
1 -3.670848 2.877400 1.048113
1 -5.252372 1.264292 -0.028947
17 -4.830561 -1.290562 -1.193937
17 -1.748522 -2.007390 -0.967518
17 0.306487 1.734041 -1.739776
17 3.369061 1.017158 -1.432678

**PCB 136 3,4 Product Complex (C-H) == PCB 136 4,5-arene oxide**

RM11 energy (hartrees): -3447.04886646

Free Energy (hartrees): -3446.924842

6 2.190330 0.907646 -0.540027
6 0.845810 1.006404 -0.512739
6 0.017178 0.039772 0.222455
6 0.606597 -0.984445 0.871853
6 2.049945 -1.271179 0.750974
6 2.874069 -0.276343 0.015088
1 3.943254 -0.169685 0.250691
8 6.170988 -0.171532 0.173905
1 5.974691 0.179161 -0.706250
1 5.827025 -1.091863 0.143972
8 4.991750 -2.665349 0.164043
1 4.735737 -2.815058 1.086344
1 4.148661 -2.490812 -0.292045
8 2.510322 -1.523546 -0.569399
1 2.514432 -1.888981 1.533965
17 -0.311087 -2.055120 1.877227
6 -1.452645 0.256195 0.309728
6 -2.340616 -0.512596 -0.451947
6 -3.720178 -0.306829 -0.350381
6 -4.221272 0.666008 0.510595
6 -3.353235 1.435153 1.275835
6 -1.982990 1.222731 1.169327
17 -0.893050 2.181505 2.130214
1 -3.738244 2.200930 1.958409
1 -5.304942 0.816431 0.578812
17 -4.825151 -1.258622 -1.291327
17 -1.709731 -1.728492 -1.514688
17 0.031279 2.333397 -1.272576
17 3.204033 2.118402 -1.242194

*Structures for 4*'*,5*'*-arene oxide mechanisms*

**PCB 132 4**'**,5**'**-arene oxide**

Path Toward C-H4

```
PCB 132 4,5 Reactant Complex (C-H4)
RM11 Energy	-3447.04568874 Hartree
Free Energy	-3446.922511 Hartree
6 2.575667 1.420427 0.004645
6 3.267588 0.107242 -0.065748
6 2.446984 -1.082326 -0.361673
6 1.101991 -1.001498 -0.411268
6 0.397570 0.290072 -0.341523
6 1.116414 1.426147 -0.222107
17 0.372044 2.991092 -0.294367
6 -1.084568 0.315546 -0.488816
6 -1.904426 -0.105590 0.562797
6 -3.298234 -0.093175 0.434914
6 -3.861859 0.350735 -0.767021
6 -3.053850 0.774590 -1.817955
6 -1.672037 0.756264 -1.675065
1 -1.030121 1.091150 -2.499337
1 -3.519562 1.118633 -2.748251
17 -5.586398 0.379255 -0.962302
17 -4.303585 -0.614441 1.741501
17 -1.171852 -0.650546 2.038201
17 0.139286 -2.416258 -0.689661
17 3.308409 -2.555568 -0.635917
1 4.327708 0.066326 -0.355620
8 3.007517 0.745832 1.179601
1 3.131653 2.339034 -0.235262
8 6.543981 -0.009562 -0.576930
1 6.283445 0.684923 0.067918
1 6.412004 -0.836633 -0.092240
8 5.600941 1.901064 1.178149
1 4.753532 1.533080 1.486505
1 5.350006 2.684140 0.665909
```

**PCB 132 4**'**,5**' **TS1 (C-H4)**

```
RM11 Energy	-3446.97891149 Hartree
Free Energy	-3446.855792 Hartree
6 2.433797 1.578180 -0.133361
6 3.246720 0.344029 -0.218715
6 2.436796 -0.876489 -0.548014
6 1.071152 -0.861241 -0.629903
6 0.372205 0.367661 -0.440008
6 1.065607 1.571515 -0.187275
17 0.174871 3.035552 0.100566
6 -1.110439 0.375003 -0.507977
6 -1.865435 -0.183366 0.526755
6 -3.262179 -0.190566 0.456019
6 -3.884040 0.364334 -0.669520
6 -3.132503 0.926020 -1.698143
6 -1.746902 0.941769 -1.612918
1 -1.149226 1.388963 -2.416803
1 -3.644861 1.356688 -2.565549
17 -5.612146 0.364980 -0.793216
17 -4.196228 -0.871523 1.738181
17 -1.052664 -0.840705 1.908666
17 0.175465 -2.305629 -0.951135
17 3.324442 -2.308618 -0.704160
1 4.030093 0.446839 -1.013616
8 3.754214 0.237337 1.074114
1 2.989789 2.498861 0.094415
8 6.149529 -0.599813 0.160370
1 6.406597 0.336418 0.192541
1 5.284357 -0.553693 0.626743
8 5.622279 2.136877 0.775315
1 4.912270 1.481358 1.008662
1 5.380550 2.449307 -0.107754
```

**PCB 132 4**'**,5**' **INT (C-H4)**

```
RM11 Energy	-3447.08933243 Hartree
Free Energy	-3446.965611 Hartree
6 2.694658 -0.993527 -1.211374
6 3.435759 0.122276 -0.526820
6 2.618833 1.197492 0.077999
6 1.285683 1.037213 0.240621
6 0.564435 -0.167824 -0.196611
6 1.255122 -1.109398 -0.862849
17 0.477059 -2.554769 -1.419756
6 -0.866860 -0.347006 0.168698
6 -1.877949 0.320568 -0.527953
6 -3.222579 0.143822 -0.181695
6 -3.539903 -0.718740 0.874377
6 -2.539236 -1.388516 1.573082
6 -1.209279 -1.199774 1.218790
1 -0.409703 -1.720961 1.760680
1 -2.814074 -2.057370 2.396381
17 -5.196938 -0.966467 1.328078
17 -4.467979 0.975852 -1.046132
17 -1.444397 1.379806 -1.831725
17 0.357832 2.252749 1.042787
17 3.481128 2.564308 0.678037
8 4.652595 0.159104 -0.508640
1 3.230009 -1.937999 -1.009610
1 2.782730 -0.804868 -2.302898
8 3.000999 -1.447752 1.711757
1 3.799412 -1.866389 1.323264
1 3.344306 -0.667853 2.170046
8 5.288557 -2.421163 0.455624
1 5.340705 -1.603551 -0.071887
1 5.083306 -3.117185 -0.185390
```

**PCB 132 4**'**,5**' **TS2 (C-H4)**

```
RM11 Energy	-3447.07459239 Hartree
Free Energy	-3446.955019 Hartree
6 -2.697615 -0.998462 0.683264
6 -3.315996 0.296241 0.432071
6 -2.461626 1.285138 -0.233171
6 -1.135946 1.042703 -0.450756
6 -0.485082 -0.190845 -0.067491
6 -1.272585 -1.136936 0.517209
17 -0.578666 -2.641849 1.042966
6 0.972083 -0.386536 -0.298552
6 1.916825 0.275461 0.495013
6 3.289038 0.099108 0.279401
6 3.707915 -0.760192 -0.742891
6 2.778550 -1.430709 -1.532265
6 1.420760 -1.240992 -1.307290
1 0.683979 -1.765720 -1.928065
1 3.130886 -2.098931 -2.326021
17 5.401130 -1.001024 -1.042623
17 4.446768 0.926441 1.263210
17 1.371705 1.328436 1.762478
17 -0.160095 2.242105 -1.232238
17 -3.209816 2.781478 -0.672733
8 -4.508450 0.532416 0.679385
1 -3.236524 -1.669798 -0.314992
1 -3.155033 -1.576346 1.501457
8 -4.008817 -2.342462 -1.182911
1 -4.881130 -2.196657 -0.694921
1 -3.787312 -3.280602 -1.062130
8 -6.021441 -1.698760 0.339066
1 -5.556405 -0.867870 0.599741
1 -6.794185 -1.402485 -0.164664
```

**PCB 132 4**'**,5**' **Product Complex (C-H4) == 4’-132**

6 -2.632825 -0.967712 0.538661

6 -3.237485 0.257832 0.232754

6 -2.430342 1.300254 -0.263819

6 -1.062337 1.104547 -0.448789

6 -0.446358 -0.119382 -0.153233

6 -1.269779 -1.135714 0.339996

17 -0.567075 -2.681578 0.741759

6 1.012945 -0.326048 -0.363953

6 1.951706 0.227669 0.514178

6 3.324454 0.031839 0.316043

6 3.748774 -0.734790 -0.774999

6 2.823892 -1.296091 -1.650517

6 1.466210 -1.090504 -1.440752

1 0.733032 -1.533405 -2.126241

1 3.179390 -1.894187 -2.497097

17 5.442752 -0.998431 -1.051839

17 4.476854 0.723448 1.405508

17 1.402951 1.165416 1.867236

17 -0.087543 2.404586 -1.062818

17 -3.184739 2.815154 -0.637880

8 -4.541719 0.470502 0.389960

1 -4.998903 -0.346566 0.764833

1 -3.242908 -1.785533 0.937071

1 -4.854484 -1.337996 -1.578206

8 -4.992509 -2.273100 -1.362177

1 -4.100477 -2.652158 -1.360456

1 -5.495530 -2.103054 0.274428

8 -5.632777 -1.755054 1.191330

1 -6.591635 -1.683411 1.305943

Pathway Toward C-H5

**PCB 132 4**'**,5**' **Reactant Complex (C-H5)**

| RM11 Energy | -3447.04513244 Hartree |
| --- | --- |
| Free Energy | -3446.920906 Hartree |

6 -3.000718 -0.707673 -0.225617

6 -3.507560 0.687820 -0.139223

6 -2.527499 1.779829 -0.319109

6 -1.205278 1.520488 -0.383494

6 -0.681710 0.145912 -0.447416

6 -1.552734 -0.883571 -0.451019

17 -1.021743 -2.526468 -0.661345

6 0.786721 -0.069690 -0.574657

6 1.622093 0.121436 0.530581

6 3.003432 -0.078964 0.428501

6 3.538696 -0.478040 -0.801901

6 2.714811 -0.672858 -1.906655

6 1.345485 -0.468748 -1.789174

1 0.691145 -0.622817 -2.656149

1 3.157953 -0.986008 -2.858737

17 5.247187 -0.736788 -0.964288

17 4.027494 0.158283 1.801197

17 0.923105 0.611801 2.041130

17 -0.053929 2.810049 -0.518253

17 -3.166945 3.380653 -0.438849

1 -4.548344 0.905413 -0.418600

8 -3.338162 -0.110509 1.018964

1 -3.660520 -1.525180 -0.550701

8 -4.202679 -3.744143 -0.590005

1 -3.401318 -3.889586 -1.110364

1 -3.869585 -3.508694 0.303248

8 -3.309394 -2.879111 1.879525

1 -2.349444 -2.753231 1.913364

1 -3.670303 -1.983463 1.972072

**PCB 132 4**'**,5**' **TS1 (C-H5)**

| RM11 Energy | -3446.98219375 Hartree |
| --- | --- |
| Free Energy | -3446.861625 Hartree |

6 -3.020285 -0.521870 0.097691

6 -3.394470 0.886857 -0.147126

6 -2.475318 1.881455 -0.345867

6 -1.114613 1.528390 -0.495340

6 -0.629339 0.185413 -0.385329

6 -1.556767 -0.778048 -0.115420

17 -1.086799 -2.397916 0.134441

6 0.820926 -0.119159 -0.534380

6 1.711727 0.156972 0.508491

6 3.074219 -0.132879 0.378428

6 3.530482 -0.710700 -0.812870

6 2.648640 -0.995630 -1.851090

6 1.298583 -0.699685 -1.709167

1 0.597763 -0.921684 -2.523397

1 3.031136 -1.450551 -2.771613

17 5.213439 -1.079974 -1.009364

17 4.171382 0.214695 1.667228

17 1.108836 0.871558 1.969319

17 0.023689 2.751074 -0.785119

17 -2.967677 3.541042 -0.427044

1 -4.461105 1.125547 -0.043307

8 -3.345622 -0.653350 1.444337

1 -3.572562 -1.212384 -0.587255

8 -4.370369 -3.238935 -1.383488

1 -3.458041 -3.466416 -1.609251

1 -4.368296 -3.215702 -0.394841

8 -4.235880 -3.054129 1.321537

1 -3.459264 -3.587388 1.541885

1 -3.926139 -2.094931 1.459708

**PCB 132 4**'**,5**' **INT (C-H5)**

| RM11 Energy | -3447.08724645 Hartree |
| --- | --- |
| Free Energy | -3446.963898 Hartree |

6 -3.561034 0.537212 0.636361

6 -2.969892 -0.814639 0.928764

6 -1.505873 -0.962299 0.730356

6 -0.763265 -0.017553 0.114513

6 -1.388268 1.230727 -0.352675

6 -2.688114 1.477949 -0.119043

17 -3.446725 2.937630 -0.643821

17 -0.391848 2.347061 -1.229911

6 0.676897 -0.252855 -0.177318

6 1.671353 0.418172 0.537000

6 3.023673 0.199701 0.251005

6 3.361251 -0.703349 -0.764254

6 2.373854 -1.374560 -1.480673

6 1.035671 -1.149298 -1.184038

1 2.665540 -2.075039 -2.271112

17 5.028196 -1.000835 -1.143264

17 4.251288 1.032020 1.138548

17 1.206622 1.527726 1.786186

17 -0.814789 -2.447977 1.294468

8 -3.656742 -1.729707 1.340225

1 -3.818473 0.994991 1.614872

1 -4.514085 0.382326 0.100310

1 0.245750 -1.673229 -1.738592

8 -3.074904 -1.223934 -1.838728

1 -2.147842 -1.048301 -2.050869

1 -3.072465 -2.141222 -1.487229

8 -3.097272 -3.709777 -0.628406

1 -2.190778 -4.043472 -0.564012

1 -3.290493 -3.349579 0.254422

**PCB 132 4**'**,5**' **TS2 (C-H5)**

| RM11 Energy | -3447.07256843 Hartree |
| --- | --- |
| Free Energy | -3446.954022 Hartree |

6 -3.298396 0.287534 0.328625

6 -2.709761 -0.996431 0.677626

6 -1.283429 -1.134543 0.358475

6 -0.524827 -0.095066 -0.094482

6 -1.117852 1.205857 -0.307984

6 -2.443128 1.384441 -0.057354

17 -3.196599 2.927728 -0.279120

17 -0.105623 2.501404 -0.872955

6 0.926593 -0.288402 -0.376767

6 1.887394 0.187224 0.519740

6 3.252605 0.010031 0.265453

6 3.641563 -0.654091 -0.903614

6 2.690070 -1.134561 -1.798810

6 1.338806 -0.952996 -1.531601

1 0.583657 -1.334361 -2.229943

1 3.019616 -1.654112 -2.705682

17 5.325590 -0.889876 -1.252561

17 4.436085 0.600019 1.379305

17 1.369799 1.000707 1.961990

17 -0.577360 -2.688936 0.675555

8 -3.370043 -1.940028 1.137985

1 -4.204447 0.550895 0.897301

1 -3.914039 -0.111452 -0.766485

8 -4.716433 -0.652217 -1.698482

1 -4.168749 -1.299442 -2.173005

1 -5.298997 -1.184803 -1.067122

8 -5.939843 -1.883857 0.237044

1 -6.163280 -2.802565 0.025471

1 -5.083196 -1.937150 0.723623

**PCB 132 4**'**,5**' **Product Complex (C-H5) == 5**'**-132**

| RM11 Energy | -3447.12411814 Hartree |
| --- | --- |
| Free Energy | -3447.000586 Hartree |

6 -3.238511 0.371556 0.283604

6 -2.685360 -0.887224 0.545165

6 -1.307005 -1.064704 0.326520

6 -0.496318 -0.029725 -0.140005

6 -1.072136 1.221868 -0.395372

6 -2.439292 1.408351 -0.181886

17 -3.177725 2.949837 -0.496552

17 -0.075008 2.523260 -0.971481

6 0.957136 -0.260484 -0.377991

6 1.914937 0.225960 0.517064

6 3.280229 0.005413 0.295101

6 3.674342 -0.714002 -0.838560

6 2.727605 -1.205298 -1.732918

6 1.377416 -0.979140 -1.498005

1 0.626103 -1.369274 -2.195854

1 3.060225 -1.767343 -2.612925

17 5.358109 -1.006840 -1.145282

17 4.459206 0.609571 1.406967

17 1.399427 1.109832 1.918123

17 -0.623271 -2.627787 0.665938

8 -3.414198 -1.912772 0.987825

1 -4.384250 -1.656764 1.051117

1 -4.309685 0.533287 0.450003

1 -4.514970 -0.917046 -1.931239

8 -5.011497 -1.700993 -1.651191

1 -4.333531 -2.296849 -1.296234

1 -5.786424 -1.328044 -0.140289

8 -5.920617 -1.181902 0.828937

1 -6.537759 -1.871881 1.113424

**PCB 136 4,5-arene oxide**

Path Toward C-H4

**PCB 136 4,5 Reactant Complex (C-H4)**

RM11 energy (hartrees): -3447.04910403

Free Energy (hartrees): -3446.925846

6 -2.174697 1.242694 -0.594171
6 -2.878427 0.002439 -0.170986
6 -2.078063 -1.051368 0.481558
6 -0.734359 -0.959001 0.545326
6 -0.023436 0.259590 0.133144
6 -0.724310 1.315065 -0.329165
17 0.040622 2.834345 -0.652788
6 1.453392 0.325158 0.304442
6 2.294848 -0.298992 -0.623816
6 3.683164 -0.237191 -0.477819
6 4.240659 0.444566 0.601052
6 3.419339 1.057910 1.538926
6 2.037738 0.990365 1.385824
17 1.013703 1.744671 2.573361
1 3.849532 1.589582 2.395042
1 5.331111 0.488433 0.703388
17 4.726501 -1.005949 -1.631858
17 1.580304 -1.161715 -1.947535
17 0.219341 -2.232939 1.231182
17 -2.956155 -2.388018 1.136081
1 -3.949049 0.042340 0.078728
8 -2.570003 0.255568 -1.537449
1 -2.729688 2.190878 -0.649798
8 -6.173163 -0.063538 0.164762
1 -5.880280 0.429633 -0.633502
1 -5.973500 -0.986439 -0.046717
8 -5.148801 1.325349 -1.987059
1 -4.277869 0.911012 -2.124767
1 -4.942965 2.211393 -1.653827

**PCB 136 4,5 TS1 (C-H4)**

RM11 energy (hartrees): -3446.98118077

Free Energy (hartrees): -3446.858642

6 -2.038006 1.372770 -0.583854
6 -2.866537 0.244557 -0.103366
6 -2.080840 -0.791371 0.648046
6 -0.716570 -0.750617 0.760518
6 -0.004830 0.340273 0.186839
6 -0.671776 1.383062 -0.488709
17 0.254032 2.662408 -1.208929
6 1.478327 0.362424 0.278209
6 2.244927 -0.466758 -0.545633
6 3.638602 -0.434711 -0.460015
6 4.262209 0.417285 0.449068
6 3.506970 1.244626 1.271406
6 2.120156 1.216203 1.176917
17 1.153871 2.245112 2.191218
1 3.992609 1.916079 1.988307
1 5.356856 0.428817 0.506855
17 4.600703 -1.450481 -1.481440
17 1.436476 -1.508366 -1.668283
17 0.162699 -2.005063 1.561857
17 -2.985453 -2.083253 1.257129
1 -3.670501 0.615006 0.584124
8 -3.340344 -0.297744 -1.296205
1 -2.575833 2.164129 -1.125053
8 -5.759543 -0.771637 -0.164291
1 -5.989629 0.106713 -0.511278
1 -4.897523 -0.910168 -0.614393
8 -5.219793 1.588322 -1.646580
1 -4.508853 0.894357 -1.659631
1 -4.965046 2.181032 -0.925238

**PCB 136 4,5 INT (C-H4)**

RM11 energy (hartrees): -3447.0923649

Free Energy (hartrees): -3446.967406

6 -2.201922 0.547708 -1.508362
6 -2.899733 -0.569664 -0.780634
6 -2.071958 -1.418051 0.106708
6 -0.795331 -1.076919 0.394306
6 -0.137469 0.112057 -0.168688
6 -0.834157 0.874232 -1.029554
17 -0.125478 2.294785 -1.720413
6 1.270726 0.420358 0.199698
6 2.317555 -0.308270 -0.381039
6 3.647610 -0.026195 -0.059438
6 3.945015 0.990699 0.844456
6 2.920730 1.723312 1.429940
6 1.598247 1.433043 1.106215
17 0.327585 2.360129 1.848738
1 3.145100 2.524950 2.142228
1 4.992252 1.205171 1.087203
17 4.941626 -0.931144 -0.779120
17 1.933065 -1.573117 -1.504353
17 0.132189 -2.025214 1.499387
17 -2.864912 -2.770956 0.823208
1 -2.859730 1.433913 -1.489330
8 -4.085336 -0.789156 -0.948139
1 -2.128615 0.232318 -2.571178
8 -3.180273 1.287023 1.197072
1 -3.922912 1.537226 0.606535
1 -3.528993 0.542015 1.706156
8 -5.239589 1.755321 -0.616440
1 -5.068854 0.875355 -0.999510
1 -4.991813 2.383471 -1.310286

**PCB 136 4,5 TS2 (C-H4)**

RM11 energy (hartrees): -3447.0786208

Free Energy (hartrees): -3446.959064

6 2.302252 -0.592166 -1.036299
6 2.860254 0.678517 -0.589462
6 1.980346 1.493210 0.255253
6 0.685310 1.128005 0.486677
6 0.094123 -0.068567 -0.066721
6 0.898947 -0.848464 -0.841104
17 0.264796 -2.295381 -1.560889
6 -1.323748 -0.416622 0.209696
6 -2.368234 0.265777 -0.429555
6 -3.701200 -0.066493 -0.171892
6 -4.005935 -1.087744 0.724434
6 -2.985458 -1.777137 1.366758
6 -1.661387 -1.436150 1.106991
17 -0.394262 -2.307022 1.926024
1 -3.214923 -2.581735 2.074070
1 -5.055351 -1.340172 0.915846
17 -4.992820 0.781144 -0.964061
17 -1.981979 1.534866 -1.547263
17 -0.323542 2.112998 1.492899
17 2.654315 2.952961 0.892805
8 4.027979 1.019155 -0.828765
1 2.906046 -1.381043 -0.178163
1 2.756332 -0.998408 -1.953523
8 3.747686 -2.146778 0.545652
1 4.584581 -1.882480 0.047346
1 3.558779 -3.063683 0.286039
8 5.646836 -1.169997 -0.949664
1 5.142187 -0.327268 -1.041833
1 6.440095 -0.929865 -0.447981

**PCB 136 4,5 Product Complex (C-H4)**

RM11 energy (hartrees): -3447.12830209

Free Energy (hartrees): -3447.003047

6 2.267692 -0.470033 -1.087784
6 2.820742 0.692551 -0.533342
6 1.984306 1.544892 0.213763
6 0.649650 1.204861 0.434195
6 0.100491 0.023797 -0.079233
6 0.941585 -0.788986 -0.844019
17 0.310758 -2.271356 -1.510704
6 -1.299800 -0.373929 0.218990
6 -2.382511 0.228286 -0.434600
6 -3.693825 -0.160670 -0.144517
6 -3.936061 -1.156776 0.797676
6 -2.874943 -1.766740 1.455208
6 -1.573754 -1.370434 1.162341
17 -0.249512 -2.136934 1.997015
1 -3.055057 -2.551309 2.198536
1 -4.969301 -1.452486 1.013160
17 -5.036254 0.586037 -0.953840
17 -2.074469 1.467605 -1.607607
17 -0.358011 2.249925 1.386280
17 2.666258 2.995768 0.872007
8 4.103074 1.011475 -0.689993
1 3.278492 -1.832858 1.057263
1 2.895185 -1.130207 -1.696218
8 3.817037 -2.531470 0.653983
1 4.829657 -1.749768 -0.527808
1 3.181395 -3.037636 0.124768
8 5.277055 -1.231223 -1.243173
1 4.624083 0.198237 -0.988538
1 6.217701 -1.222897 -1.013344

*Structures for 5*'*,6*'*-arene oxide mechanisms*

**PCB 132 5**'**,6**'**-arene oxide**

Path Toward C-Cl

**PCB 132 5**'**,6**' **Reactant Complex (C-Cl)**

```
RM11 Energy	-3447.04810026 Hartree
Free Energy	-3446.923782 Hartree
6 -1.482139 0.761777 0.245496
6 -2.968097 0.771478 0.246997
6 -3.655353 -0.518851 0.117891
6 -2.946373 -1.635051 -0.119764
6 -1.473373 -1.642972 -0.101068
6 -0.747260 -0.530996 0.156123
6 0.725769 -0.550994 0.350001
6 1.565845 0.158264 -0.515923
6 2.952004 0.172389 -0.323626
6 3.490464 -0.536788 0.756180
6 2.664710 -1.246078 1.624039
6 1.291673 -1.251046 1.417869
1 0.636768 -1.806223 2.100895
1 3.110176 -1.791224 2.463721
17 5.204083 -0.541677 1.027169
17 3.978799 1.046990 -1.405253
17 0.867335 1.020769 -1.850050
17 -0.674198 -3.163285 -0.345297
17 -3.775028 -3.144745 -0.344894
1 -4.743714 -0.561215 0.236161
1 -3.475041 1.606393 0.753587
8 -2.173833 1.184305 -0.886451
17 -0.639267 2.052723 1.127716
1 -2.329057 3.088527 -1.027485
8 -2.478230 4.015581 -0.769080
1 -1.590746 4.363191 -0.596669
8 -3.640291 3.627006 1.694542
1 -3.227937 3.823984 0.823082
1 -2.898139 3.351494 2.249673
```

**PCB 132 5**'**,6**' **TS1 (C-Cl)**

```
RM11 Energy	-3447.00242833 Hartree
Free Energy	-3446.881792 Hartree
6 1.680825 0.856110 0.058290
6 3.107668 0.532113 0.187709
6 3.557858 -0.783485 0.213915
6 2.675333 -1.776761 -0.155947
6 1.270472 -1.518835 -0.465279
6 0.770172 -0.267456 -0.389104
6 -0.678221 0.016264 -0.542729
6 -1.555433 -0.284839 0.507876
6 -2.924694 -0.016191 0.390567
6 -3.407254 0.545327 -0.797159
6 -2.544275 0.830161 -1.850409
6 -1.186624 0.568497 -1.720314
1 -0.506205 0.789679 -2.549510
1 -2.946163 1.260411 -2.774441
17 -5.098498 0.886741 -0.975064
17 -4.004050 -0.380011 1.691173
17 -0.948180 -1.018210 1.956858
17 0.241796 -2.858067 -0.844976
17 3.236155 -3.380944 -0.229953
1 4.597689 -1.026592 0.455385
1 3.773887 1.396059 0.343219
8 1.517193 1.232066 1.305852
17 1.501408 2.258996 -1.227896
1 1.965875 2.884246 1.575916
8 2.228292 3.834205 1.575983
1 1.610382 4.246374 0.954413
8 4.431717 3.394011 0.033648
1 3.658401 3.621249 0.602906
1 4.046832 3.296314 -0.848658
```

**PCB 132 5**'**,6**' **INT (C-Cl)**

```
RM11 Energy	-3447.07926173 Hartree
Free Energy	-3446.957383 Hartree
6 2.769183 -0.658777 0.279882
6 1.277254 -0.641470 0.556949
6 0.523418 0.585618 0.218768
6 1.191714 1.705515 -0.144064
6 2.664154 1.747875 -0.241504
6 3.399899 0.654246 -0.020384
1 4.493134 0.689064 -0.103853
17 3.427094 3.251449 -0.654028
17 0.322599 3.147346 -0.517412
6 -0.951614 0.524518 0.383630
6 -1.722621 -0.287760 -0.455858
6 -3.109536 -0.381480 -0.288641
6 -3.718207 0.354820 0.734161
6 -2.961404 1.168603 1.572137
6 -1.586528 1.249617 1.393984
1 -0.985376 1.887041 2.054403
1 -3.461544 1.734332 2.366125
17 -5.434640 0.262598 0.972323
17 -4.054557 -1.385083 -1.332947
17 -0.947483 -1.171853 -1.732810
8 0.710377 -1.621071 1.004312
1 2.914219 -1.339151 -0.591293
17 3.633706 -1.465891 1.641516
1 1.548872 -3.257895 0.792079
8 2.041489 -4.055372 0.514860
1 2.712651 -4.174620 1.202546
8 3.570916 -3.079492 -1.533118
1 3.010751 -3.492376 -0.834854
1 4.436547 -2.982467 -1.110845
```

**PCB 132 5**'**,6**' **TS2 (C-Cl)**

```
RM11 Energy	-3447.07230852 Hartree
Free Energy	-3446.951013 Hartree
6 1.277113 -0.672529 0.849999
6 2.707133 -0.801898 0.556092
6 3.481911 0.379990 0.240032
6 2.860006 1.518330 -0.142718
6 1.412731 1.582805 -0.126167
6 0.644247 0.548697 0.325960
6 -0.837517 0.611895 0.414290
6 -1.630505 -0.274675 -0.325419
6 -3.026962 -0.249264 -0.221198
6 -3.625941 0.679696 0.637157
6 -2.848977 1.566939 1.375826
6 -1.465467 1.527772 1.261544
1 -0.849300 2.222538 1.845786
1 -3.339515 2.285028 2.042506
17 -5.354252 0.738541 0.792178
17 -3.997567 -1.344305 -1.144187
17 -0.874334 -1.404541 -1.406818
17 0.658960 3.028738 -0.711865
17 3.785506 2.904199 -0.641982
1 4.577302 0.314707 0.237718
17 3.546737 -2.060498 1.489613
8 0.608413 -1.561164 1.388346
1 2.565072 -1.368151 -0.613828
1 0.908296 -3.154580 0.586352
8 1.187417 -3.845322 -0.052501
1 1.917015 -4.306366 0.388682
8 2.377614 -2.180525 -1.688149
1 1.895314 -2.923560 -1.208659
1 3.245749 -2.536372 -1.940462
```

**PCB 132 5**'**,6**' **Product Complex (C-Cl)**

```
RM11 Energy	-3447.12607507 Hartree
Free Energy	-3446.999881 Hartree
6 -1.318046 -0.669406 -0.930567
6 -2.721338 -0.701093 -0.892782
6 -3.463872 0.353351 -0.380230
6 -2.807602 1.480363 0.101970
6 -1.411187 1.549546 0.056533
6 -0.664768 0.481679 -0.453253
6 0.822391 0.529030 -0.513017
6 1.605076 -0.295025 0.303053
6 3.003725 -0.260953 0.229754
6 3.613908 0.617008 -0.672702
6 2.846008 1.445809 -1.485246
6 1.460743 1.397830 -1.400786
1 0.850926 2.049247 -2.039179
1 3.345448 2.126133 -2.184161
17 5.344716 0.686552 -0.789825
17 3.962911 -1.283756 1.243161
17 0.837031 -1.363389 1.437414
17 -0.594223 2.957403 0.661648
17 -3.746538 2.792953 0.746148
1 -4.558478 0.301185 -0.356653
17 -3.548409 -2.104021 -1.509712
8 -0.588924 -1.697399 -1.390561
1 -0.862872 -2.534559 -0.884498
1 -1.938280 -0.926765 1.864297
8 -2.340605 -1.795621 2.020121
1 -3.273627 -1.670722 1.788075
8 -1.313694 -3.571491 0.220099
1 -2.076223 -4.078901 -0.095353
1 -1.674650 -2.975970 0.923660
```

Pathway Toward C-H

**PCB 132 5**'**,6**' **Reactant Complex (C-H same as C-Cl)**

See above

**PCB 132 5**'**,6**' **TS1 (C-H)**

```
RM11 Energy	-3446.98129812 Hartree
Free Energy	-3446.860054 Hartree
6 -1.473101 -0.642129 -0.406706
6 -2.921028 -0.649736 -0.054869
6 -3.485275 0.725806 0.067862
6 -2.708879 1.830214 0.087609
6 -1.283697 1.698391 -0.091168
6 -0.645857 0.468837 -0.315421
6 0.830575 0.371339 -0.485680
6 1.624203 -0.186150 0.522802
6 3.011008 -0.292592 0.361485
6 3.591221 0.170830 -0.825200
6 2.807388 0.726007 -1.832937
6 1.433096 0.823248 -1.660622
1 0.810201 1.258947 -2.452026
1 3.285343 1.079501 -2.753301
17 5.306148 0.056749 -1.057423
17 3.987685 -0.981636 1.609491
17 0.875414 -0.753001 1.980508
17 -0.325678 3.095682 0.019843
17 -3.391664 3.401803 0.361605
1 -4.565411 0.805162 0.251524
1 -3.494649 -1.245475 -0.801043
8 -2.857949 -1.210487 1.223720
17 -0.804674 -2.160414 -0.767622
1 -3.235706 -2.704521 0.942023
8 -3.493655 -3.633270 0.624102
1 -2.656460 -4.115534 0.575535
8 -4.107535 -3.160221 -1.976790
1 -3.917084 -3.382920 -1.032421
1 -3.233985 -3.133834 -2.390305
```

**PCB 132 5**'**,6**' **Product Complex (C-H) == PCB 132 4**'**,5**'**-arene oxide**

See above

**PCB 136 5,6-arene oxide**

Pathway Toward C-Cl

**PCB 136 5,6 Reactant Complex (C-Cl)**

RM11 energy (hartrees): -3447.0510487

Free Energy (hartrees): -3446.926098

6 -1.068308 0.926354 0.155868
6 -2.547383 1.011739 0.031693
6 -3.270974 -0.219589 -0.307442
6 -2.589647 -1.335332 -0.619123
6 -1.127566 -1.420998 -0.471337
6 -0.389337 -0.386292 -0.011575
6 1.058450 -0.500902 0.298553
6 1.996993 0.177602 -0.490376
6 3.362118 0.098957 -0.204944
6 3.803261 -0.653563 0.880442
6 2.887163 -1.320069 1.684046
6 1.528862 -1.236751 1.392481
17 0.398718 -2.051959 2.434009
1 3.224689 -1.904644 2.547140
1 4.876540 -0.708873 1.096249
17 4.519569 0.934527 -1.190824
17 1.427618 1.127031 -1.826286
17 -0.362317 -2.938437 -0.803001
17 -3.452136 -2.763414 -1.100809
1 -4.366500 -0.219704 -0.288175
1 -3.064169 1.810067 0.584956
8 -1.636405 1.504551 -0.975354
17 -0.242639 2.065057 1.238655
1 -1.700700 3.418656 -0.920291
8 -1.825113 4.322713 -0.580258
1 -0.944131 4.590402 -0.279366
8 -3.228241 3.729516 1.711554
1 -2.733832 3.999508 0.904819
1 -2.545443 3.375715 2.297781

**PCB 136 5,6 TS1 (C-Cl)**

RM11 energy (hartrees): -3447.00375231

Free Energy (hartrees): -3446.883376

6 -1.254669 0.903893 -0.218368
6 -2.699631 0.654418 -0.392079
6 -3.190197 -0.599549 -0.721763
6 -2.336382 -1.681688 -0.603806
6 -0.936885 -1.550702 -0.209884
6 -0.397276 -0.332228 0.003099
6 1.024466 -0.173678 0.399701
6 2.004460 0.114305 -0.557164
6 3.336244 0.294302 -0.167229
6 3.698208 0.166974 1.170207
6 2.743610 -0.169990 2.122088
6 1.421614 -0.348056 1.730830
17 0.248489 -0.845489 2.914467
1 3.024663 -0.303025 3.172813
1 4.745200 0.313394 1.459893
17 4.555161 0.662660 -1.345582
17 1.570123 0.161713 -2.233106
17 0.019646 -2.979991 -0.024420
17 -2.938370 -3.235207 -0.929120
1 -4.234666 -0.749683 -1.015388
1 -3.331415 1.556546 -0.360919
8 -1.046786 1.578860 -1.312515
17 -1.049057 1.939475 1.398698
1 -1.400347 3.257742 -1.149776
8 -1.603467 4.189993 -0.899868
1 -1.019325 4.353053 -0.144755
8 -3.911102 3.479045 0.374343
1 -3.104104 3.802565 -0.091668
1 -3.572409 3.164480 1.224400

**PCB 136 5,6 INT (C-Cl)**

RM11 energy (hartrees): -3447.08252753

Free Energy (hartrees): -3446.960566

6 -0.971621 0.698923 0.477851
6 -2.442395 0.676265 0.106200
6 -2.973599 -0.610497 -0.417318
6 -2.162871 -1.625182 -0.733635
6 -0.707620 -1.527894 -0.515232
6 -0.135610 -0.441078 0.052312
6 1.319981 -0.331393 0.322069
6 2.105083 0.549531 -0.434031
6 3.473236 0.681101 -0.181543
6 4.069851 -0.063963 0.832260
6 3.306581 -0.936745 1.597038
6 1.944637 -1.061441 1.339274
17 1.001279 -2.147951 2.319002
1 3.766132 -1.523085 2.400467
1 5.143850 0.048305 1.020783
17 4.445390 1.766738 -1.123642
17 1.353942 1.471945 -1.697234
17 0.274248 -2.857259 -0.996217
17 -2.805019 -3.088822 -1.409030
1 -4.055071 -0.685890 -0.585399
1 -2.578425 1.465353 -0.669284
8 -0.485165 1.645467 1.068674
17 -3.434484 1.242046 1.501431
1 -1.399446 3.249262 1.067942
8 -1.917491 4.062857 0.909167
1 -2.574837 4.072114 1.619993
8 -3.392574 3.300015 -1.261206
1 -2.860588 3.641833 -0.504634
1 -4.234752 3.045801 -0.857717

**PCB 136 5,6 TS2 (C-Cl)**

RM11 energy (hartrees): -3447.07580115

Free Energy (hartrees): -3446.953338

6 -0.896358 0.745040 0.842777
6 -2.298884 1.003814 0.501578
6 -3.113009 -0.067424 -0.037892
6 -2.527245 -1.161934 -0.575135
6 -1.089193 -1.314137 -0.494274
6 -0.303854 -0.422264 0.176288
6 1.166089 -0.577841 0.316199
6 2.026851 0.324691 -0.326248
6 3.412950 0.212422 -0.188875
6 3.957628 -0.799728 0.597123
6 3.122627 -1.695341 1.251959
6 1.742875 -1.574887 1.111150
17 0.718203 -2.689921 1.971411
1 3.539848 -2.489927 1.880675
1 5.046558 -0.877709 0.695821
17 4.475352 1.325814 -0.993118
17 1.348526 1.586936 -1.306240
17 -0.361872 -2.673466 -1.277147
17 -3.485495 -2.390218 -1.348215
1 -4.201617 0.062791 -0.084997
1 -2.064281 1.716181 -0.555955
8 -0.209777 1.506832 1.531505
17 -3.121993 2.152073 1.579923
1 -0.393674 3.214094 0.940796
8 -0.595594 4.000885 0.390809
1 -1.333581 4.433582 0.846350
8 -1.774456 2.648080 -1.517204
1 -1.284287 3.283949 -0.911116
1 -2.609453 3.085493 -1.752768

**PCB 136 5,6 Product Complex (C-Cl)**

RM11 energy (hartrees): -3447.12985661

Free Energy (hartrees): -3447.003361

6 -0.945236 0.681068 0.914073
6 -2.340730 0.801469 0.809099
6 -3.099846 -0.098533 0.075449
6 -2.470790 -1.154117 -0.576883
6 -1.084835 -1.307293 -0.479754
6 -0.324512 -0.399200 0.263803
6 1.153968 -0.527802 0.368153
6 1.998153 0.363591 -0.307163
6 3.387571 0.250647 -0.202518
6 3.950162 -0.755229 0.578663
6 3.130547 -1.646341 1.259088
6 1.748554 -1.523791 1.150326
17 0.735874 -2.638882 2.024493
1 3.561780 -2.438961 1.880848
1 5.041092 -0.833408 0.651197
17 4.431306 1.355432 -1.041874
17 1.301763 1.616429 -1.287249
17 -0.289916 -2.611806 -1.300917
17 -3.430273 -2.270174 -1.499923
1 -4.187227 0.021370 0.007710
1 -1.455336 1.547966 -1.774797
8 -0.192661 1.561728 1.588487
17 -3.134168 2.124164 1.617067
1 -0.393186 2.497816 1.244828
8 -0.701740 3.767282 0.367504
1 -1.437172 4.257646 0.763653
8 -1.778778 2.462479 -1.771272
1 -1.078581 3.351149 -0.448059
1 -2.731919 2.379901 -1.614263

Pathway Toward C-H

**PCB 136 5,6 Reactant Complex (C-H same as C-Cl)**

See above

**PCB 136 5,6 TS1 (C-H)**

RM11 energy (hartrees): -3446.98454646

Free Energy (hartrees): -3446.862922

6 -1.134267 0.689125 0.298156
6 -2.553909 0.722099 -0.151392
6 -3.068916 -0.630166 -0.513658
6 -2.264000 -1.708623 -0.633478
6 -0.862008 -1.582427 -0.328594
6 -0.276538 -0.384675 0.111214
6 1.182680 -0.280970 0.381889
6 2.020483 0.404772 -0.507401
6 3.388111 0.519562 -0.242487
6 3.926209 -0.049974 0.908682
6 3.108356 -0.730911 1.801503
6 1.747746 -0.837666 1.533973
17 0.723483 -1.685626 2.656178
1 3.524697 -1.178896 2.710467
1 5.000745 0.045830 1.103125
17 4.429799 1.367324 -1.339335
17 1.336591 1.113681 -1.933297
17 0.142109 -2.931613 -0.547253
17 -2.882472 -3.232810 -1.184098
1 -4.130405 -0.700882 -0.787476
1 -3.193343 1.195086 0.628078
8 -2.414833 1.460309 -1.329559
17 -0.528311 2.149236 0.915630
1 -2.849387 2.898923 -0.863758
8 -3.144331 3.768474 -0.434476
1 -2.322822 4.259634 -0.293321
8 -3.868520 2.932072 2.045064
1 -3.634506 3.287120 1.152621
1 -3.017582 2.862215 2.498571

**PCB 136 5,6 Product Complex (C-H) == PCB 136 4,5-arene oxide**

RM11 energy (hartrees): -3447.04958182

Free Energy (hartrees): -3446.927603

6 1.031233 -0.821957 0.755704
6 2.472170 -1.096673 0.584745
6 3.274886 -0.044305 -0.101167
6 2.574139 1.180210 -0.540100
6 1.230038 1.270991 -0.470820
6 0.422315 0.244810 0.200502
6 -1.042728 0.457705 0.352899
6 -1.958323 -0.197317 -0.476362
6 -3.332883 -0.000156 -0.314331
6 -3.800738 0.856945 0.678265
6 -2.904928 1.517680 1.510113
6 -1.539918 1.312122 1.341135
17 -0.413482 2.139639 2.377430
1 -3.264628 2.194429 2.293163
1 -4.880742 1.004614 0.794637
17 -4.468740 -0.814282 -1.341957
17 -1.366139 -1.255321 -1.719679
17 0.391527 2.658452 -1.082004
17 3.559565 2.457890 -1.155971
1 4.346411 0.053907 0.124066
1 2.943494 -1.784140 1.298889
8 2.901398 -1.242155 -0.759558
17 0.126163 -1.958578 1.706874
1 1.546093 -2.623187 -1.431149
8 1.154517 -3.429827 -1.059577
1 0.236545 -3.201486 -0.856326
8 2.612365 -4.157692 1.192331
1 2.086510 -3.918173 0.399461
1 1.997301 -4.041503 1.928426
